# A Computational Approach for Structural and Functional Analyses of Disease-associated Mutations in the Human *CYLD* Gene

**DOI:** 10.1101/2023.11.16.567342

**Authors:** Arpita Singha Roy, Tasmiah Feroz, Md. Kobirul Islam, Md. Adnan Munim, Dilara Akhter Supti, Nusrat Jahan Antora, Hasan Al Reza, Supriya Gosh, Newaz Mohammad Bahadur, Mohammad Rahanur Alam, Md Shahadat Hossain

**Author notes:** Corresponding authors: Mohammad Rahanur Alam, Md Shahadat Hossain. Contributed equally (Arpita Singha Roy, Tasmiah Feroz).

## Abstract

Tumor suppressor Cylindromatosis protein (CYLD) regulates NF- κB and JNK signaling pathway by cleaving K63 linked poly-ubiquitin chain from its substrate molecules and thus preventing the progression of tumorigenesis and metastasis of the cancer cells. Mutations in CYLD can cause aberrant structure and abnormal functionality leading to tumor formation. In this study, we utilized several computational tools such as PANTHER, PROVEAN, PREDICT- SNP, POLYPHEN 2, PHD SNP, PON P2, and SIFT to find out deleterious nsSNPs. We also highlighted the damaging impact of those deleterious nsSNPs on the structure and function of the CYLD utilizing Consurf, I-Mutant, SDM, Phyre2, HOPE, Swiss PDB Viewer, and Mutation 3D. We shortlisted 18 high-risk nsSNPs from a total of 446 nsSNPs recorded in the NCBI database. Based on the conservation profile, stability status, and structural impact analysis we finalized 13nsSNPs. Molecular docking analysis and molecular dynamic simulation concluded the study with the findings of two significant nsSNPs (R830K, H827R) which have a remarkable impact on binding affinity, RMSD, RMSF, Radius of gyration, and hydrogen bond formation during CYLD-ubiquitin interaction. The principal component analysis compared native and two mutants R830K, H827R of CYLD that signifies structural and energy profile fluctuations during molecular dynamic (MD) simulation. Finally, the Protein-protein interaction network showed CYLD interacts with 20 proteins involved in several biological pathways that mutations can impair. Considering all these in silico analyses, our study recommended conducting large-scale association studies of nsSNPs of CYLD with cancer as well as designing precise medications against diseases associated with these polymorphisms.

## Introduction

CYLD or, cylindromatosis lysine 63 deubiquitinase is a tumor suppressor protein that generally performs deubiquitinase activities essential for a variety of cellular and signaling processes (1). CYLD is mainly a cytoplasmic protein that belongs to the ubiquitin specific protease (USP) family (2) and is abundant in the brain (3), skeletal muscle (4), and immune cells (5). CYLD processes larger substrate molecule by cleaving lysin63 linked ubiquitin chains from that molecule (6, 7) and thereby involved in corresponding cellular events namely: cell cycle control (8), cellular differentiation (9), oncogenesis (10), cellular proliferation (11, 12), and apoptosis (13). Mutation in CYLD can give rise to the constant activation and deregulation of cell survival proteins associated with tumorigenesis (10). Several studies have demonstrated that mutated CYLD gene greatly contributes to the familial cylindromatosis, Brooke-Spiegler syndrome, and multiple familial trichoepitheliomas (14, 15).

CYLD gene encodes a 956 amino acid long protein with a weight of about 110kD, and is localized on the chromosome 16q12-q13 (1) with 19 introns and 21 exons (16). It contains a C terminal conserved catalytic domain (USP) along with three N terminal Cap-Gly domains (1). Cap-Gly domain is crucial for the interaction with proteins involved in the NF-κB pathway (17), while USP domain is important for hydrolyzing the ubiquitin chain. This carboxyl terminal (USP) catalytic domain changes its target proteins by deubiquitinating Lys63-linked ubiquitin chains of specific substrates that are vital in various cellular signaling events, especially in NF- κB pathway (2, 18). By deubiquitinating TRAF2/TRAF6, NF-κB essential modifier (NEMO), CYLD act as a key regulator in the typical p65/NFκB pathway (19, 20). CYLD also contributes greatly by preventing the Bcl3 from being localized in the nucleus and thus controls tumor development and proliferation (11). Therefore, any mutation disrupting the deubiquitinating (DUB) activity of CYLD may leads to oncogenic function gain, as DUB activity is fundamental for CYLD as a tumor suppressor (21, 22).

Several polymorphisms have been identified as being responsible for the impaired activity of the CYLD gene, which finally leads to the tumorigenesis (23, 24). The consequence of missense mutation on CYLD and the manner in which it is associated with cancer formation is not fully explored yet using computational approaches. Therefore, *in-silico* analysis on nsSNPs of CYLD will help to demonstrate the potential role of mutation contributing towards the molecular mechanisms of various cancer types.

By considering all these facts, we have conducted an extensive analysis and explored numerous bioinformatics tools to investigate the functional and structural effect of various nsSNPs on the CYLD protein and narrow down the list of the high risk nsSNPs for our present study. In addition, we performed structural stability analysis, conservation analysis, protein-protein interactions followed by molecular docking analysis with its interacting molecules. Cancer associated nsSNP identification further validated by molecular dynamic simulation analysis where root mean square deviations (RMSD), root-mean-square fluctuation (RMSF), radius of gyration (Rg) analysis, H-bond analysis were taken into consideration. This study will help us to identify cancer prone genotypes related to this CYLD protein as well as future research on CYLD mutations.

## Methodology

### Assortment of nsSNPs

The information about human CYLD protein along with its amino acid sequence were assembled from National Center for Biotechnology Information (NCBI). Details of SNPs (reference ID, location, residual variations, global minor allele frequency) were retrieved from dbSNP(25) (https://www.ncbi.nlm.nih.gov/projects/SNP/) a publicly accessible database for genetic variations available in NCBI for further computational analyses.

### Screening of most deleterious nsSNPs

We exploited seven different *in silico* nsSNP prediction tools (SIFT, PANTHER, POLYPHEN 2, PROVEAN, PHD SNP, PON P2 and PREDICT SNP) for the assessment of most deleterious nsSNPs having significant effect on the structure and function of the CYLD protein. [2] SIFT (26) (Sorting Intolerant From Tolerant) (https://sift.jcvi.org/www/SIFT_seq_submit2.html) a sequence homology based algorithm, determines the effect of amino acid substitution over the physical and functional properties of a protein. SIFT provides prediction score against our submitted rsID for query nsSNPs where prediction score <0.05 is regarded as intolerant and >0.05 regarded as tolerant. SIFT result obtained from PREDICT SNP(27).

PANTHER(28) (https://www.pantherdb.org/tools) database integrates the evolutionary conservation history with hidden Markov models (HMMs) to analyze the probability of a damaging effect of nsSNPs on the functionality of a protein and their interacting ability with other proteins. PANTHER provides position specific evolutionary conservation scores when protein sequences along with human missense SNPs are submitted as a query.

Polyphen2 (29) (Polymorphism Phenotyping v2) (https://genetics.bwh.harvard.edu/pph2/) is a tool that employs machine learning methods, considering multiple sequence alignment to classify the damaging impact of allele change over the structure and function of a protein categorized as probably damaging with probabilistic score (0.85 to 1.0), and possibly damaging with probabilistic score (0.15 to 1.0). Information about amino acid substitution along with FASTA sequence of a protein is required for the query submission.

PROVEAN (30, 31) (Protein Variation Effect Analyzer) (https://provean.jcvi.org/index.php) predicts the deleterious consequences of single or multiple amino acid changes (insertion and deletion) on the biological function of a protein. PROVEAN considers -2.5 as a cut off value where amino acid substitution score >-2.5 is regarded as deleterious mutation PHD SNP (32) (Predictor of human Deleterious Single Nucleotide Polymorphisms) (https://snps.biofold.org/phd-snp/phd-snp.html) applies support vector machines (SVMs) to distinguish a genetic disease linked point mutation from the neutral polymorphisms. Protein sequences, mutation profile information such as position of mutation, and mutated residues are required as an input file.

PON P2 (33) (http://structure.bmc.lu.se/PON-P2/) a machine learning-based algorithm classifies the amino acid alteration into 3 categories: pathogenic, neutral and unknown groups. This tool predicts probability score of variant tolerance with respect to sequence conservancy, biological and physical properties of amino acids, gene ontology features and functional annotations of alteration sites.

PredictSNP (27) (https://loschmidt.chemi.muni.cz/predictsnp) is a consensus program based on Protein Mutant Database and the UniProt database annotations. It confirms the accuracy of the results acquired from eight renowned prediction tools (MAPP, nsSNPAnalyzer, PANTHER, PhD-SNP, PolyPhen-1, PolyPhen-2, SIFT and SNAP) which signifies the impact of amino acid alteration on functional activity of a protein.

### Identification of deleterious nsSNPs on different CYLD domains

InterPro (34) (https://www.ebi.ac.uk/interpro/) program ascertained the position of mutant nsSNPs on the CYLD protein according to the protein families, domains, and functional regions by integrating information from various protein-signature databases such as Pfam, PROSITE, PRINTS, ProDom, SMART, PIRSF, TIGRFAMs, and PANTHER, the structure-based SUPERFAMILY and Gene3D. Protein ID or FASTA sequences are used for the query searching.

### Structural stability determination

I-Mutant and SDM tools specify the structural stability alteration of a protein as a result of deleterious point mutation. I-Mutant (35) (https://folding.biofold.org/i-mutant/i-mutant2.0.html) is a web server established on a support vector machine utilizing the thermodynamic database ProTherm to offer the changing free energy (DDG value) and the reliability index value (RI) of a protein, and thus evaluate the level of changing structural stability in mutant proteins.

Site Director Mutator (36, 37) (SDM) is a web-based application that predicts the significant effect of mutation on protein stability. This tool computes stability score considering amino acid substitution patterns among homologous proteins from the same family and estimates free energy variation by comparing wild type and mutant type proteins.

### Evolutionary conservation analysis

Consurf (38) (https://consurf.tau.ac.il) applies either an empirical Bayesian method or a maximum likelihood (ML) method for the interpretation of the evolutionary conservancy of a particular amino acid at a specific position of a protein that signifies its structural and functional importance. In order to assess conservation score (ranges from 1 to 9) of an amino acid in a protein, Consurf analyzes the phylogenetic relationship, multiple sequence alignment and sequence homology of the protein. Conserved nsSNPs were considered for the further investigations.

### Structural effect of nsSNPs on CYLD

Project HOPE (39) (https://www3.cmbi.umcn.nl/hope) is a web browser that uses Distributed Annotation System along with UniProt database to analyze the impact of a point mutation on the structure of a protein. Significant findings regarding structural variations between mutant and native residues are produced through 3D homology modelling using YASARA program. FASTA sequence or Uniprot id is submitted as query file.

SWISS-PDB VIEWER (40, 41) (v4.1.0) (https://spdbv.vital-it.ch/) computes the energy minimization of a protein for different amino acid substitutions. This tool utilizes its mutation tool and thereby select best rotamer of the mutated protein and calculate the energy minimization state of a native and mutant 3D protein model using Nomad-Ref server. This server performed the energy minimization of a 3D structure of a protein using GROMACS program as a default force field which is built on the methods of steepest descent, conjugate gradient, and L-BFGS (Limited-memory Broyden–Fletcher–Goldfarb–Shanno) algorithm.

### 3D structure modelling to visualize the effect of nsSNPs

Three-homology modeling tools namely: SWISS MODEL, Phyre 2, and I -TASSER were used to create 3D structure of native and mutant proteins.

I-TASSER (42–44) (https://zhanglab.ccmb.med.umich.edu/I-TASSER/) is a web server that operates replica exchanged Monte Carlo Simulations, and thereby builds 3D structure of a full-length protein through splicing continuous threading alignments. This tool offers the comparative analysis of the I-TASSER models using confidence score, TM-score (template modelling score) and RMSD (root mean square deviation) value which is conducted by benchmark scoring system.

Phyre 2 (45) (https://www.sbg.bio.ic.ac.uk/phyre2) is a web server based on advanced distant homology detection algorithms that generates 3D protein model and therefore provides analysis on the influence of amino acid variants on structure and function of a protein. Intensive mode was selected for developing 3D structure of CYLD protein. This mode constructs a full-length sequence model of a query protein combining different template models with high confidence score and sequence similarity. Then TM Align tool (46) (https://zhanglab.ccmb.med.umich.edu/TM-align/) is incorporated to compare the mutant protein structure against the native one. Tm Align calculates template modelling score (TM-Score) and root mean square deviation (RMSD) on the basis of structural similarities between two proteins. It generates TM-Score ranges between 0 and 1, where TM Score 1 signifies perfect similarity between two protein structures. Significant deviation between native and mutant structure is estimated considering higher RMSD value.

SWISS MODEL (47) (https://swissmodel.expasy.org/) server combines sequence alignment and template structure to develop three-dimensional structure of a protein. QMEAN scoring function applies for the model quality assessment to validate the reliability of the resultant models of both wild type and mutant type proteins.

### Molecular docking analysis

HADDOCK (48, 49) (High Ambiguity Driven Protein-Protein Docking) (https://wenmr.science.uu.nl/) a web tool was used to perform molecular docking analysis to understand the effect of deleterious point mutation over the binding affinity of CYLD with its interacting proteins. Protein-protein docking was carried out by the HADDOCK, with default settings for all parameters. The PDB structure of wild type CYLD protein (PDB id-2VHF) was taken from SWISS-MODEL (50) and Ramachandran Plot was used to validate the structure. Refinement was done before performing docking analysis using refinement interface in HADDOCK. C-PORT (51) server (http://haddock.chem.uu.nl/services/CPORT) identified the active and passive residues of CYLD and Ubiquitin protein. PRODIGY (52) (https://wenmr.science.uu.nl/prodigy/) web server performs the calculation of the binding affinity between protein- protein dock complexes. BIOVIA Discovery Studio (53) was used to perform docking complex analysis along with image generation.

### Identification of the cancer associated nsSNPs

The web tool mutation3D (54) (http://www.mutation3d.org/) enables users to easily identify the cancer causing mutation clusters collected from 975,000 somatic mutations recorded in 6,811 cancer sequencing studies. This tool applies 3D clustering approach to find out amino acid substitution of a protein that can cause cancer when a target protein along with its mutations inserted as a query. We used this tool to look at the nsSNPs that can predispose to cancer.

### Molecular dynamics simulation

YASARA (55) simulation software uses AMBER14 (56) as a force field to analyze the changing outcome of wild type and mutant dock complex by allowing them to interact for a fixed period. The simulating chamber was permitted to contain 20 Å around the protein that is filled with water at 0.997 g/ml density. Initially protein-protein dock complex was cleaned along with the H-bond network optimization. The steepest descent method was used to minimize the energy of the protein-protein complex. In order to evaluate the short-range Coulomb and van der Waals interaction, the cut-off radius was limited to 8 Å. The PME (Particle Mesh Ewalds) method was utilized to assess the long-range electrostatic interactions. Simulations were accomplished under constant pressure in water and Berendsen thermostat process controlled the simulation temperature at 298 K. Counter ions (Na or Cl) were introduced to maintain a concentration of 0.9 % (NaCl) to neutralize the system at pH 7.4. Simulation was executed for 100 ns under constant temperature and pressure by maintaining a time step interval of 2.5 femtoseconds (fs). This tool provides following types of data such as root mean square deviation (RMSD), root mean square fluctuation (RMSF), radius of gyration, total number of hydrogen bonds, and helix, sheet, turn, coil values after the completion of simulations.

### Principal component analysis

Principal component analysis was performed to determine the dynamics of biological system by reducing data complexity along with the retrieval of the coordinated movements found in the simulations. A correlation matrix was built to represent variations detected in MD trajectories and offers the prediction of the first two principal components based on the calculation of the eigenvectors and eigenvalues (57, 58). We performed principal component analysis (PCA) by considering bond distances, bond angles, dihedral angles, planarity, van der Waals energies, and electrostatic energies and thus analyze the structural and energy changes of the wild CYLD- ubiquitin complex and mutant CYLD-ubiquitin complexes (H827R, R830K). Minitab software (Minitab 19, Minitab Inc., State College, PA, USA) a multivariate data analysis tool performed Principal Component Analysis (PCA) to signify variations among different groups by analyzing 100 ns MD simulation data.

### Protein-protein interacting network analysis

STRING (59) (https://string-db.org/), online database helped with better understanding of the molecular interactions of CYLD with other proteins. STRING produced the data in Simple Interaction Format (SIF) or GML format which were visualized by CytoScape (60, 61) a freely accessible java based software. The overall workflow represented by **Fig. 1**.

**Fig. 1:**
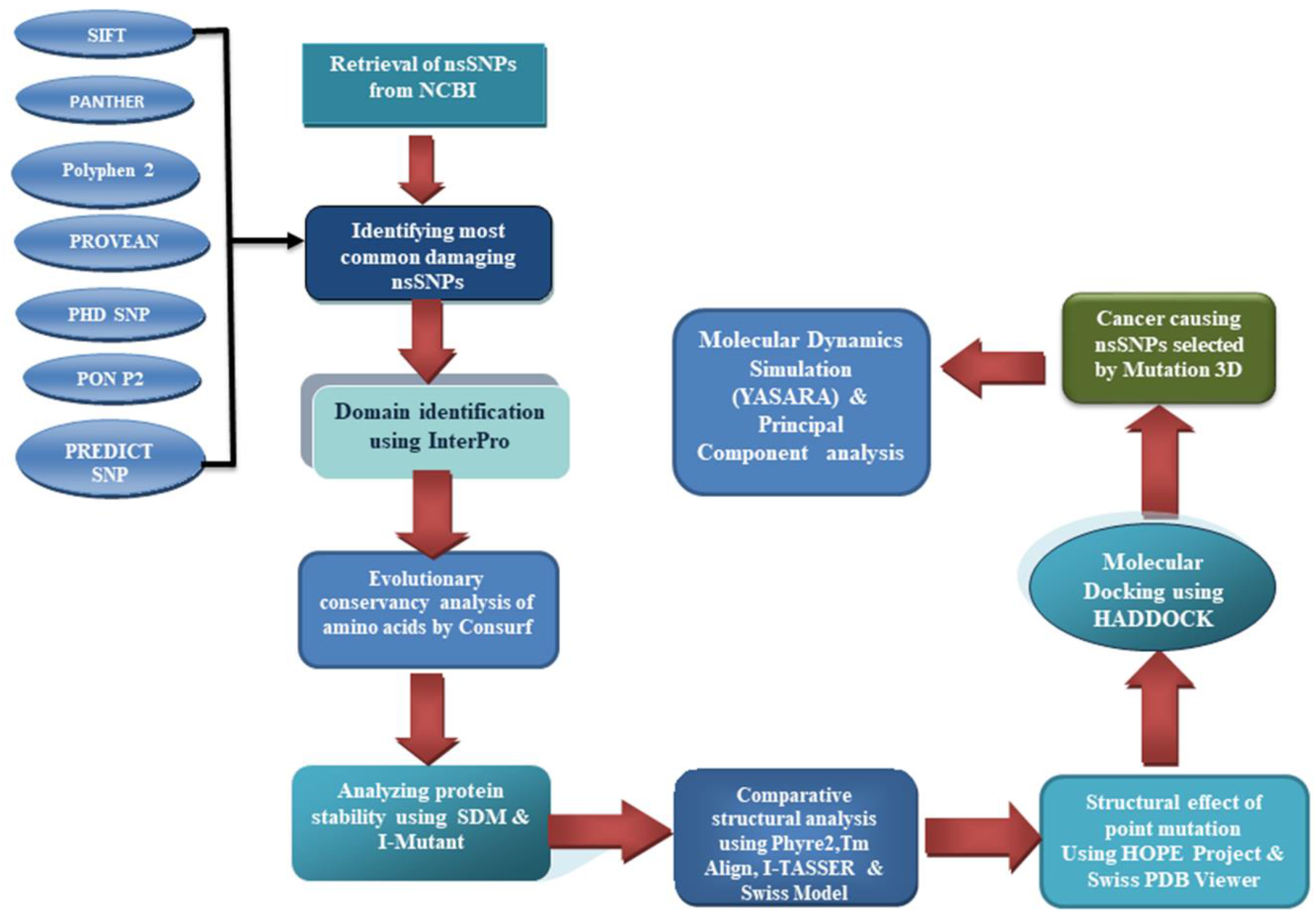
A workflow representing all the *in-silico* tools utilized for the computational analysis.

## Result

### Retrieval of nsSNPs

We retrieved all the reported SNPs found in CYLD from NCBI dbSNP database. This database contains a total of 13653 SNPs, where 13111 SNPs were in the intronic region, 2066 SNPs were in the non-coding region, and 658 SNPs were in the coding region. In case of coding region, 413 SNPs were missense and 245 SNPs were synonymous. A total of 446 missense variants have been found as some reference SNP ID (rsID) contain multiple SNPs at a single position. We considered all missense variants for our further analysis.

### Identification of damaging nsSNPs

All missense variants obtained from dbSNP database were subjected to seven different deleterious SNP prediction tools namely: PANTHER, PROVEAN, PREDICT SNP, PolyPhen-2, PhD-SNP, PON P2, and SIFT,to determine their damaging consequences on the structure and function of CYLD protein. Each server predicted different amount of pathogenic nsSNPs. Finally, we targeted 18 common nsSNPs which were predicted to be deleterious by all the seven *in silico* tools out of 446 nsSNPs **(Table. 1).**

### Domain identification of CYLD

Interpro was used to conduct functional analysis of the CYLD protein by categorizing them into protein families and identifying the active sites and domain of the corresponding protein., This domain identification analysis revealed that CYLD contains two functional domain which are Cap-Gly domain (127-540 Amino acid) and USP domain (592-950 AA) **(S1 Fig)**. Among 18 nsSNPs,16 nsSNPs were positioned at the USP domain whereas 2 nsSNPs were in the Cap-Gly domain.

### Evolutionary conservation analysis of deleterious nsSNP in CYLD

Evolutionary conserved amino acid residues in a protein plays specific roles in various functional biological cascades. Point mutation in such conserved residues result in aberrant structural and functional properties of a protein. ConSurf web server facilitates the analysis of the evolutionary conservancy and solvent accessibility of the amino acid residues of CYLD protein **(Fig. 2)**. Among 18 high-risk nsSNPs, it predicted 16 highly conserved amino acid residues with conservation score 9 **(S1 Table),** whereas 2 nsSNPs (L610F, L781P) were conserved and average conserved respectively. These conserved residues are classified as structural or functional depending on their location in the structure of a protein. Amino acid residues exposed in the surface of a protein are considered functional whereas buried residues are predicted to be structural (62–64) Therefore, these findings further highlight the importance of the deleterious effects of nsSNPs situated at those buried and exposed residues of the CYLD protein .

**Fig. 2.1:**
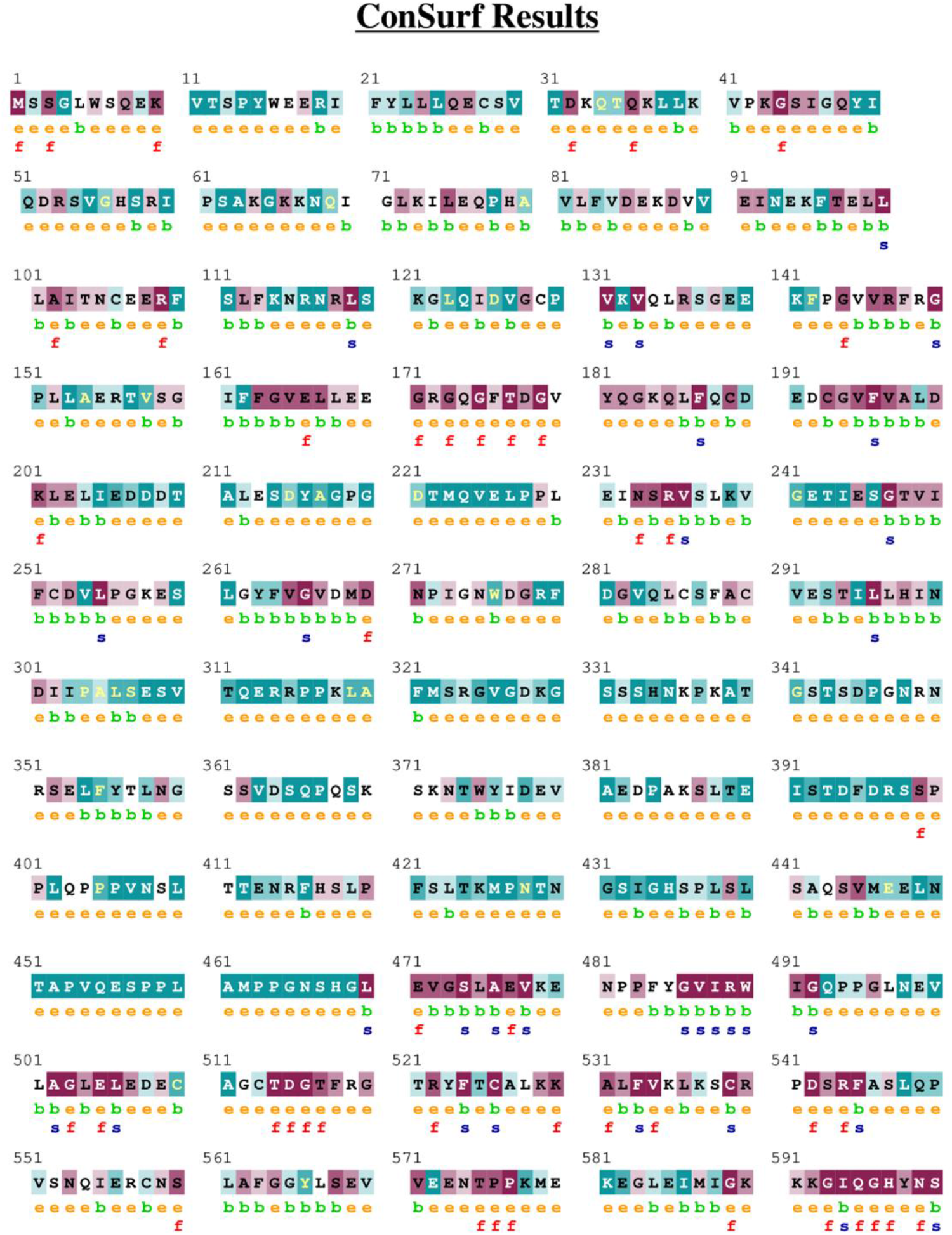
Conservation analysis of amino acid residues of CYLD using Consurf server

**Fig. 2.2:**
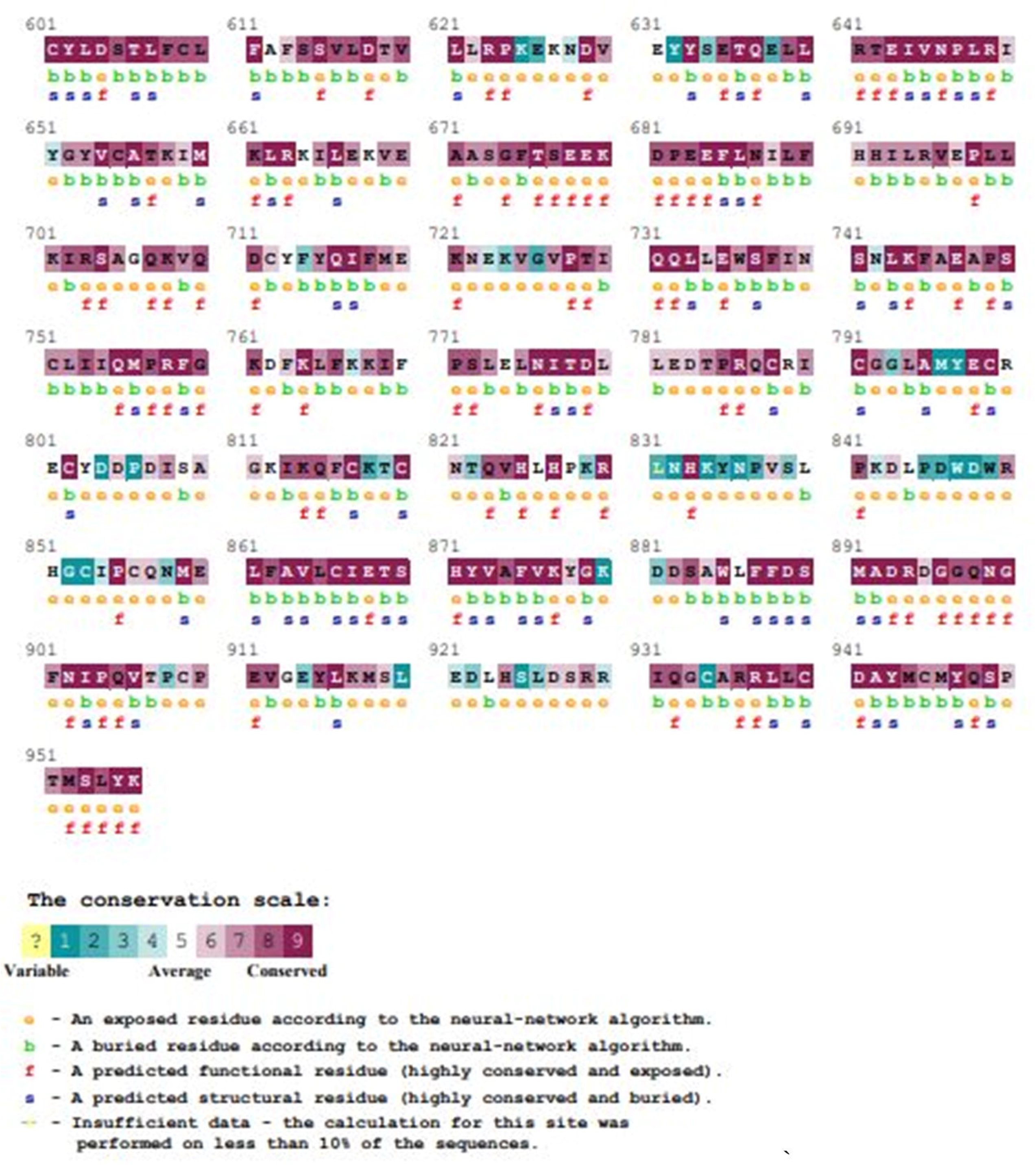
Conservation analysis of amino acid residues of CYLD using Consurf server

### Prediction of changing structural stability

Amino acid substitutions are thought to have damaging impacts on protein stability. Our selected 18 nsSNPs were subjected to the I-mutant and SDM tools to analyze the changes in stability of CYLD protein due to point mutations. I-Mutant carried out the calculations based on free energy change values ΔΔG and reliability index value (RI). It predicted 14 nsSNPs that decreased the stability whereas 4 nsSNPs (R894W, P698L, P698S, L781P) increased the stability of CYLD **(Table 2).** SDM tool identified 4 nsSNPs (R894W, P698L, P698S, P698T) as stabilizing and 14nsSNPs as destabilizing that are specifically responsible for the protein instability and dysfunction **(Table. 2).** Both tools confirmed three common variants: R894W, P698L, and P698S, which increased the stability of the protein after mutation. Therefore, we excluded these three missense variants from further analysis.

**Table 1:**
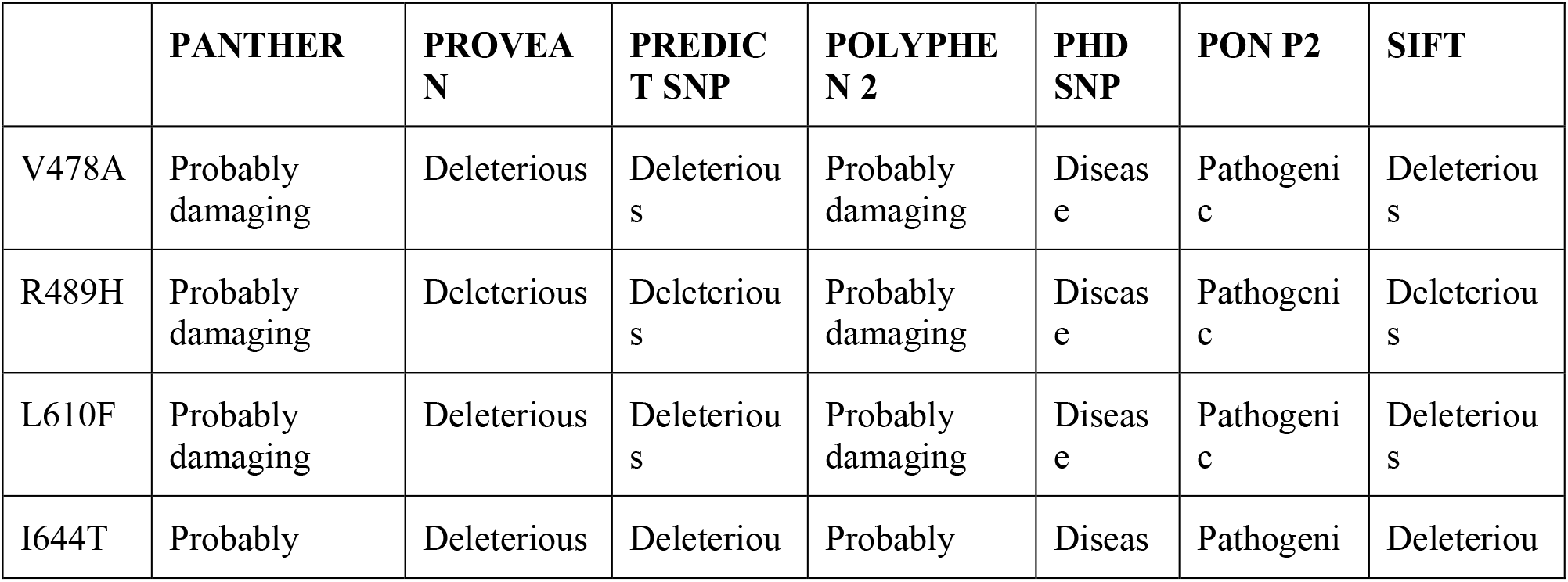
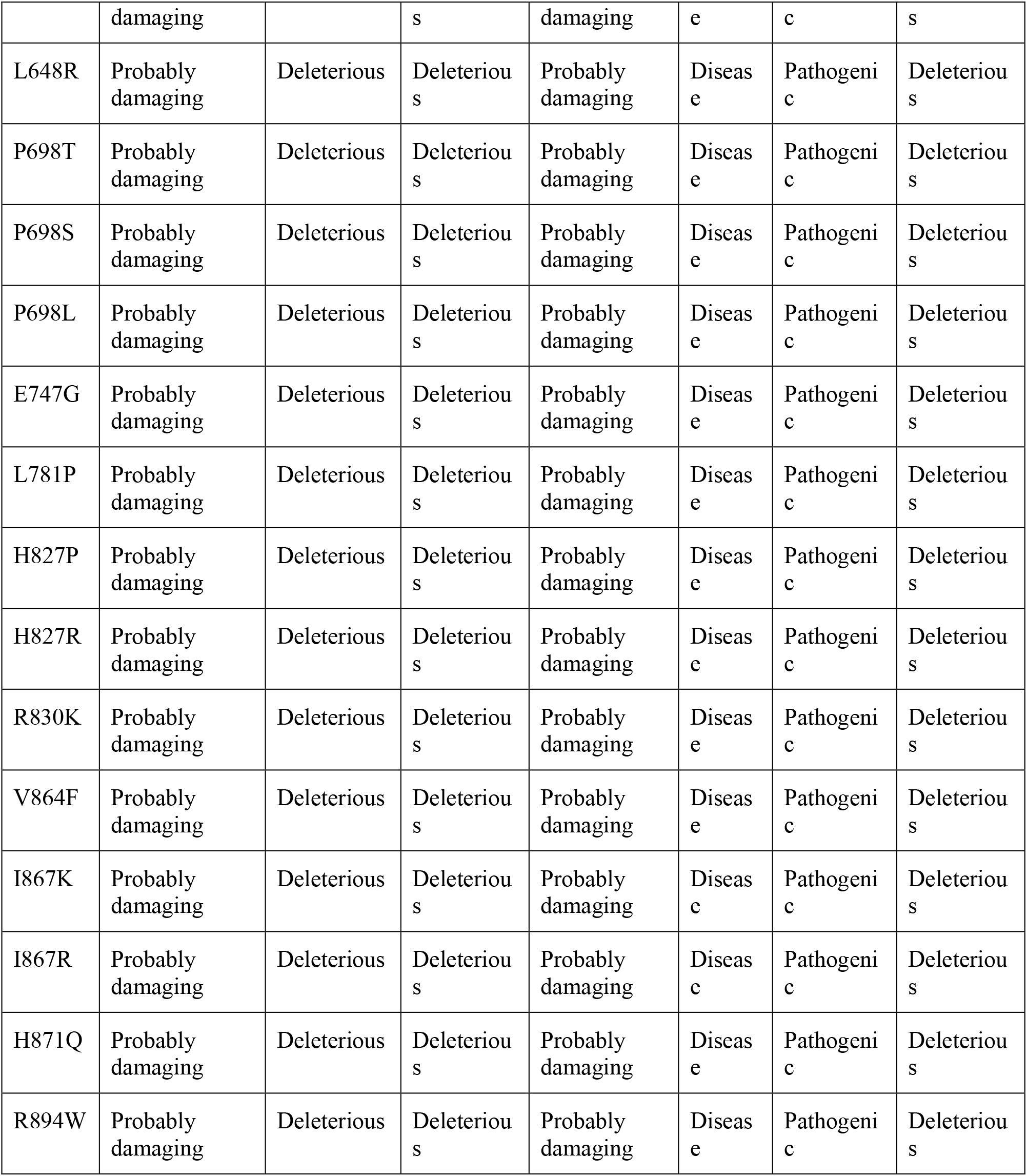
List of highly deleterious nsSNPs screened by seven computational programs.

**Table 2:**
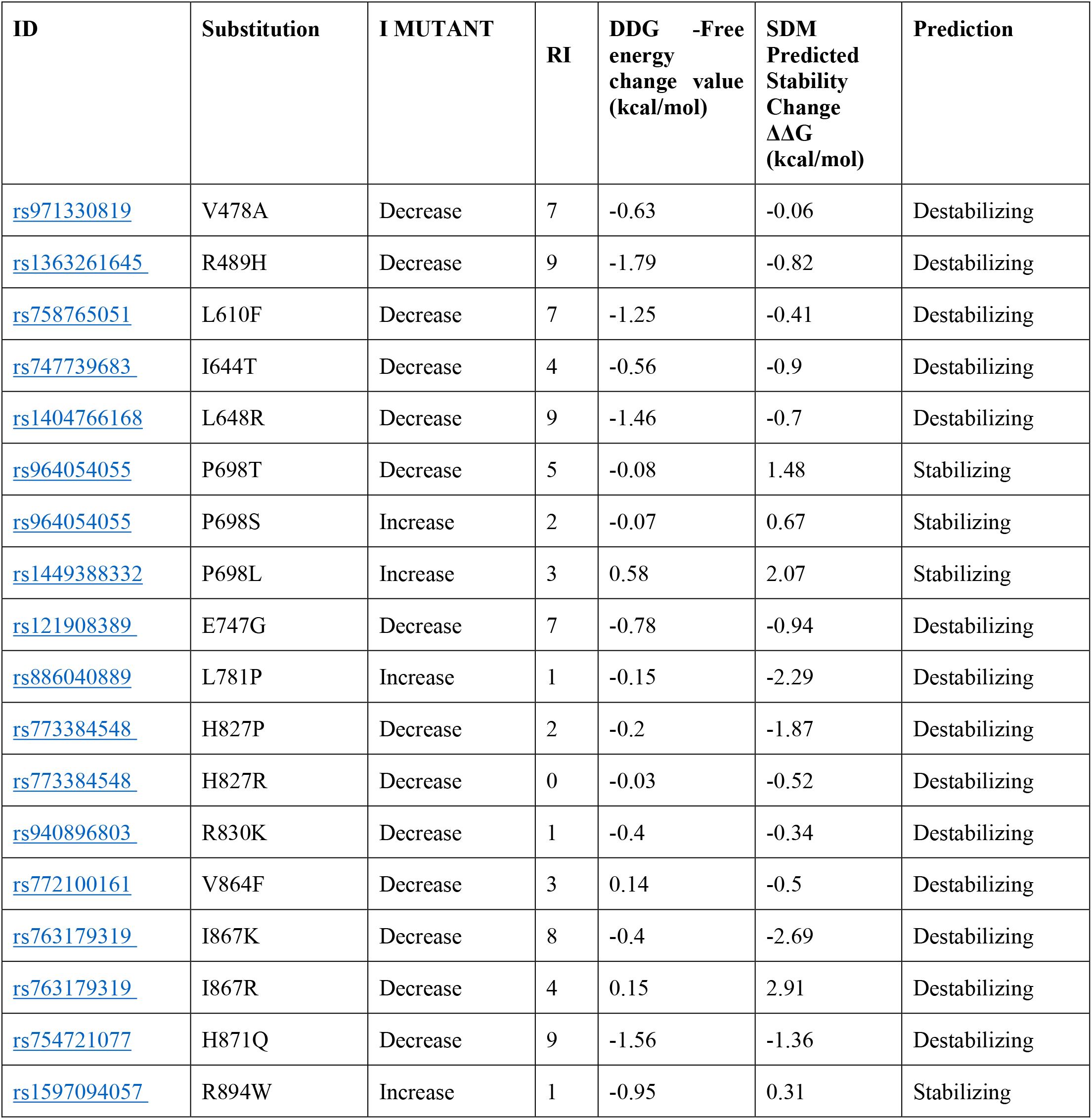
Alterations in the structural stability profile of the CYLD protein by I-MUTANT and SDM tool.

### Comparative structural analysis of wild type and mutant CYLD

We exploited three computational algorithms: Phyre2, I-TASSER and SWISS MODEL to perform comparative structural analysis of wild type and 15 mutants of CYLD. These tools were used to generate 3D structures of wild type protein and mutated proteins as the whole structure of CYLD is not available in Protein Data Bank. Phyre2 utilized 2VHF and 1IXD as template for the 3D protein structure modeling for USP and CAP-Gly domain of CYLD, respectively.

Then we determined variations between native and mutant 3D protein structure with the help of TM- align tool. Two variants, V478A and R489H located in CAP-Gly domain with RMSD value 0 indicated no dissimilarities between these two variants when compared with wild type protein. On the other hand, comparison between native and 13 missense variants exhibit significant TM-score and RMSD value **(S2 Table).** Missense variants located on USP domain showed greater RMSD values and among them H871Q, I867K, H827P variants had the highest TM-Score. We also used I-TASSER for an additional structural study of 13 nsSNPs to verify the relevance of these findings. This server generated the top 5 reliable superimposed models of mutants over the wild type protein based on minimum confidence score (C-score) along with significant TM Score and RMSD value. To conduct comparative analysis of all atom of a protein we incorporated SWISS MODEL. This homology modeling server utilized 2vhf.2. A template to build the structure of CYLD and its mutant. SWISS MODEL server also determined solvation, torsion, qmean, and Cβ value for both native and mutants which are shown in the (**S3 Table)**.

### Analysis of structural effect of point mutation on CYLD protein

The project HOPE server analyzed the physiochemical alterations of CYLD protein structure as a result of amino acid substitutions (**S4 Table)**. We observed functional CYLD mutations that lead to significant change in size and hydrophobicity in all mutant residues. L610F, L648R, H827R, V864F, I867K, and I867R mutant residues are larger whereas I644T, P698T, E747G, L781P, H827P, R830K, and H871Q mutant residues are smaller when compared to wild-type residues. Repulsion was generated between the mutant residue and neighboring residues when a charge was added in H827R **(Fig. 3),** I867K, and I867R position due to mutation. On the other hand, protein-folding problems can arise in L648R missense variant and empty space formed in the core of the protein when mutation occurs in I644T and R830K **(Fig. 3)** position. Moreover, H871Q, I644T, L781P, E747G, H827P, and R830K nsSNPs also resulted in the loss of interactions.

**Fig. 3:**
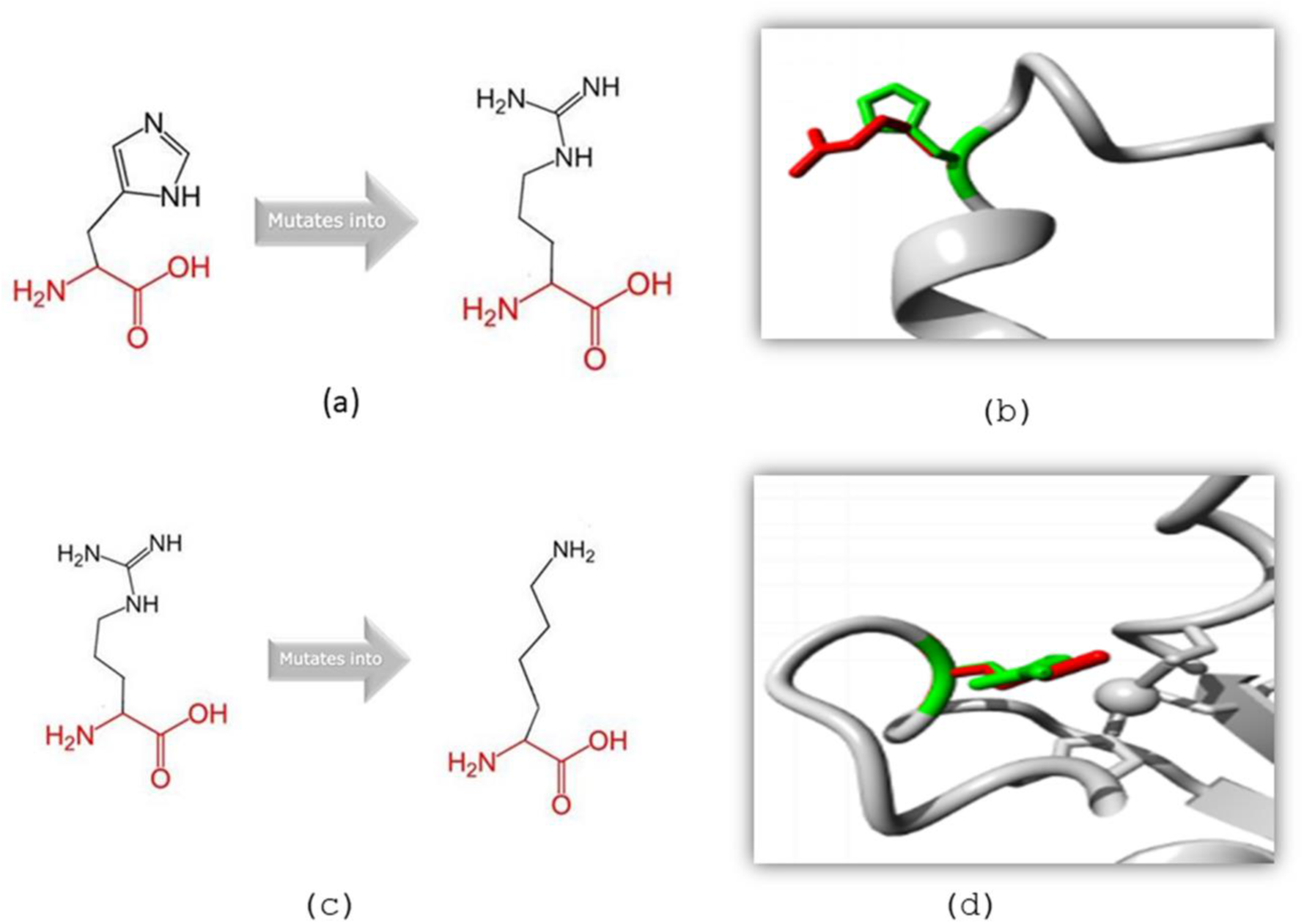
Structural effect of the point mutation on variant H827R (a and b) and R830K (c and d) predicted by HOPE server. (Green color indicates wild and red color indicates mutant residues.)

SWISS-PDB VIEWER calculates energy state variations of a protein when the position of an atom or molecules change. We determined the deviations in the energy minimization state of CYLD structure geometry in wild type and 13 variants. Total energy of the wild type protein was -20130.191 kj/mol which was decreased in case of L610F, L648R, P698T, and I867R variants after energy minimization. Other missense variants showed increase in total energy after energy minimization. Among them, H827R showed significant increase in total energy (-15956.584kj/mol) after energy minimization (**S5 Table)**. Structural changes in h bond in R830K showed in Fig. 4.

**Fig. 4:**
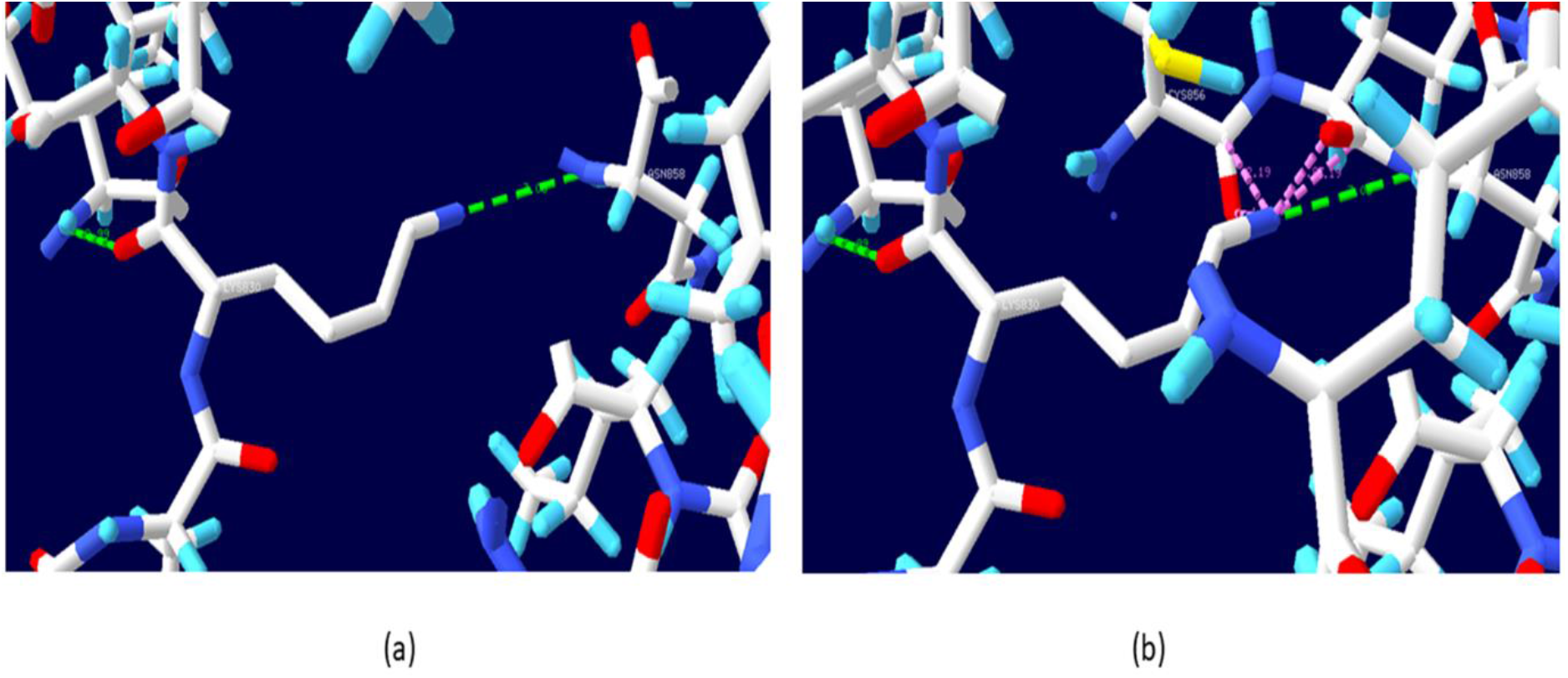
**Structural effect analysis of R830K by Swiss PDB viewer. (**Fig (a) represents R830 where R830 forms two H-bonds (2.99 Å, 3.08 Å) which are indicated by green discontinuous line (b) represents 830K where K830 clashes with C856 (1.43 Å,1.90 Å,2.03 Å) which are indicated by pink discontinuous line along with two H-bonds (green line)).

### Molecular docking analysis

Molecular docking analysis was performed between the CYLD (wild and mutants) and ubiquitin chain using HADDOCK to see how mutant interacts with ubiquitin compared to native CYLD protein. The PDB structure for USP domain of CYLD was taken from SWISS MODEL using PDB ID: 2VHF (583aa-956aa) as a template as some residues were missing in the PDB structure. Ramachandran plot **(S1 Fig)** verified the model where 92.47% amino acid residues are in favored region which assured the good quality of the model. On the other hand, ubiquitin chain was derived from protein data bank (PDB id: 3WXE). Native and mutant structures were refined using refinement algorithm of HADDOCK. Active and passive residues of Ub and CYLD protein was determined by CPORT server that ensures binding of Ub protein in the appropriate binding site of CYLD. The binding affinity between native CYLD and ubiquitin was -14.8 KJ/mol. Among 13 high-risk nsSNPs, binding affinity increased in L610F, I644T, E747G, V864F, I867R, and H871Q whereas binding affinity decreased in P698T, L781P H827P, H827R, R830K, and I867K after mutation. Among them, significant reduction was observed in four nsSNPs namely: R830K, H827R, P698T, and L781P with binding affinity -12.7KJ/mol, -12.8 KJ/mol, -13.0 KJ/mol, and -13.4 KJ/mol respectively. Binding affinity of these variants showed significant deviation compared to native protein. Binding affinity and dissociation constant of all docking complexes found from HADDOCK for both wild type and mutant structures determined by PRODIGY server (**S6 Table)**. We also determined the H-bond interaction between CYLD– ubiquitin dock complex applying BIOVIA Discovery Studio where wild type CYLD formed 24 h bonds with ubiquitin .In case of R830K and H827R, total 22 and 13 h bonds found in CYLD-Ubiquitin dock complex respectively **(Fig. 5).** Interacting residues along with bond category and bond distance represented in the (**S7 table).**

**Fig. 5:**
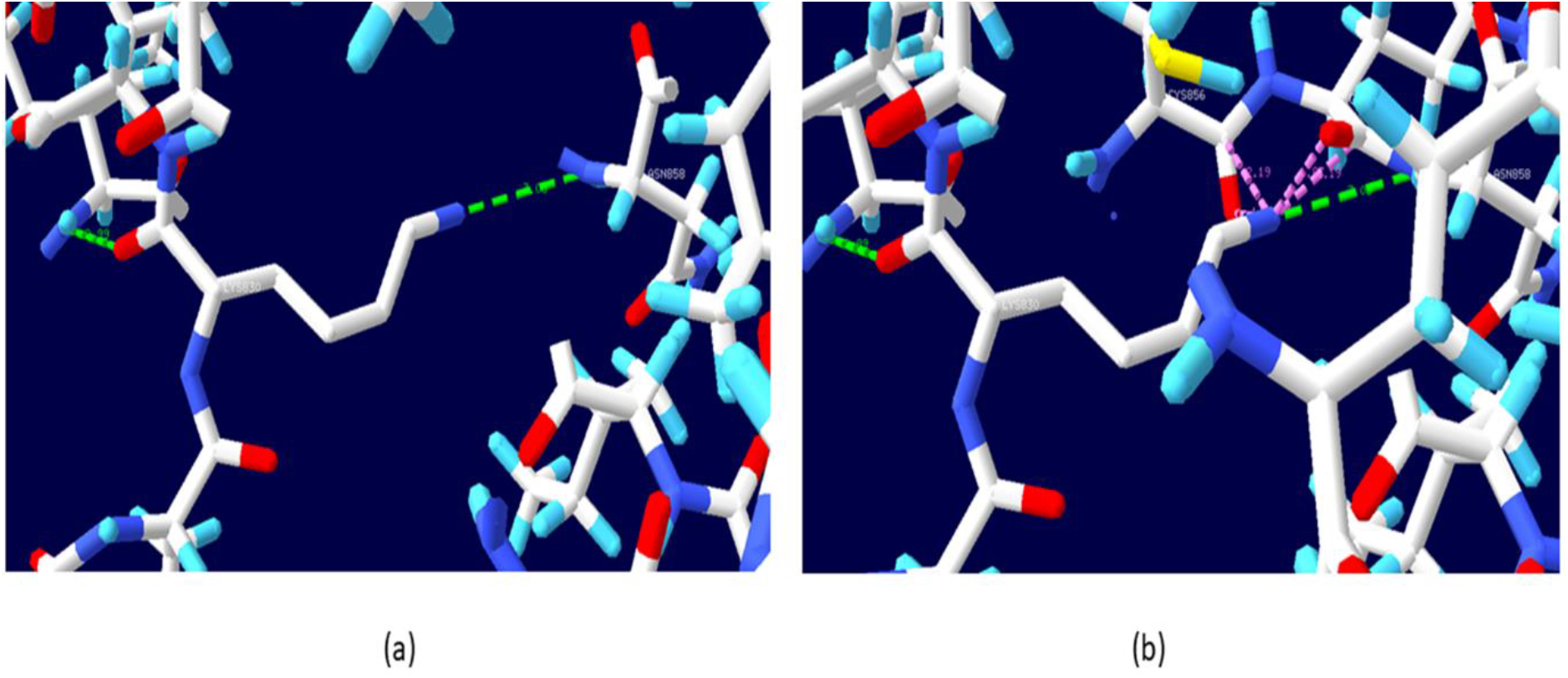
Molecular docking analysis and visualization by BIOVIA Discovery Studio. (Blue indicates USP domain of CYLD and chocolate color indicates ubiquitin. Fig (a) indicates CYLD- ubiquitin docking complex, Fig (b) indicates H-bond interactions between wild CYLD-ubiquitin dock complex, Fig (c) represents H-bond interactions between mutant (H827R) CYLD-ubiquitin dock complex, Fig (d) represents h-bond interactions between mutant (R830K) CYLD-ubiquitin dock complex.)

### Prediction of cancer causing nsSNPs

As CYLD is a tumor suppressor protein, loss of activity due to mutation can result in cancer. Mutation 3D is a server that predicts deleterious nsSNPs which are associated with human cancer. This analysis revealed the association of H827R and R830K with cancer **(Fig. 6)** and we considered these two nsSNPs for further analysis.

**Fig. 6:**
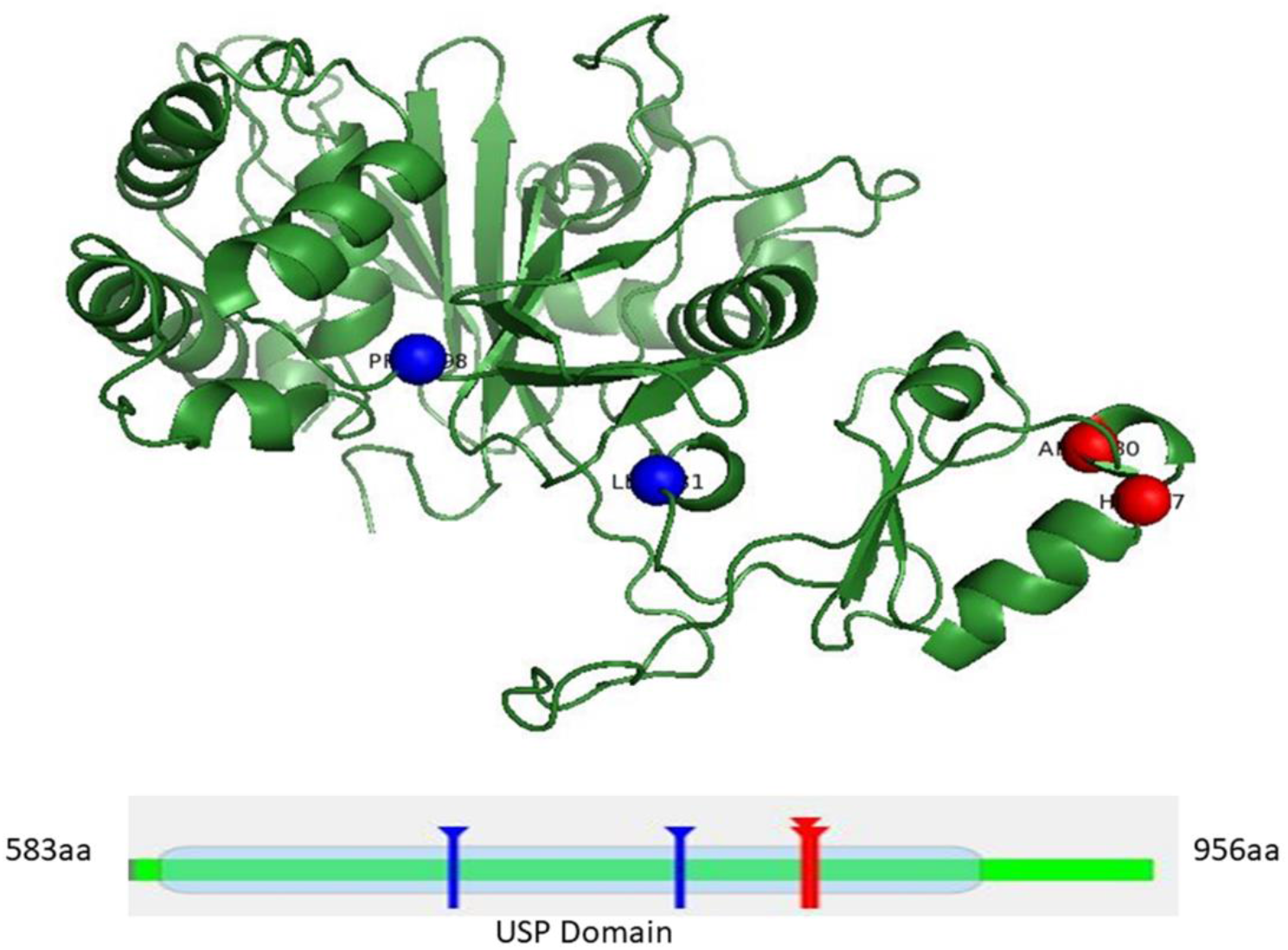
Mutation 3D predicted the association of H827R and R830K (red mark) with cancer

### Molecular Dynamic (MD) simulation

MD simulation was conducted to examine the deviation of the native and mutant CYLD- ubiquitin complex in relation to its initial conformation under physiological conditions. Trajectory analysis from the simulation enables the stability and flexibility of the system to be computed. The simulations were performed for 100 ns to investigate the structural flexibility, stability, and hydrogen bonding between the protein-protein complexes.

The overall changes in the protein stability due to the mutation were calculated by considering the root mean square deviations (RMSD) values. Mutant R830K and H827R complex exhibited great deviation in comparison to native CYLD-ubiquitin complex **(Fig. 7)**. The average RMSD value for native complex was ∼ 3.388 Å, which was increased in R830K and H827R mutant complex to ∼ 5.278 Å,and ∼ 4.9575 Å respectively (Fig. 7). The highest RMSD value for native complex was 4.418 Å at 39 ns meanwhile the highest deviation was noticed for H827R complex with a 6.327 Å RMSD value at 79.75 ns in compared with its initial structure. On the other hand, R830K complex showed deviation at 71 ns with 6.087 Å. Native complex showed mild deviation in RMSD value till 39 ns and then the native complex stayed stable within the range of 2-4 Å for the rest of the time which indicates stability of the protein. On the other hand, mutant H827R showed an increasing tendency till 16.75 ns and thereafter from 16.75 ns to 28 ns RMSD value was decreased and again it started to increase at 29 ns to 100 ns at the range of 5-6 Å which is much higher than wild CYLD. Fluctuations that observed in this RMSD values indicate decreasing stability of the mutant H827R. In case of R830K complex, fluctuation rate is greater than mutant H827R. In R830K, the RMSD value started to increase at 11.5 ns and became unstable throughout the overall simulation period within the range of 5-6 Å which is higher than wild CYLD. Considerable fluctuations observed after 80 ns and it continued up to 100 ns.

**Fig. 7:**
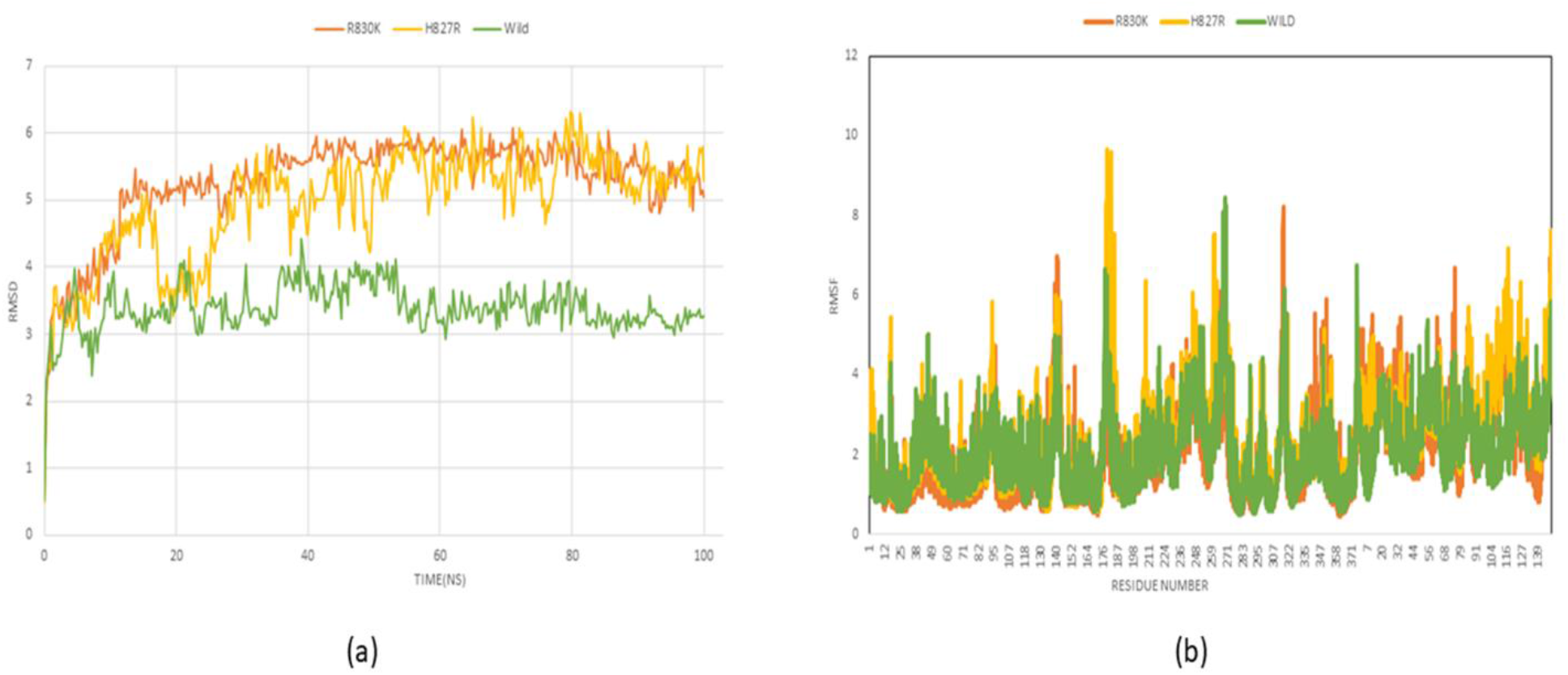
**Molecular dynamic simulation analysis performed by YASARA. (**Fig (a) exhibits RMSD analysis of the Cα atoms of the structure of protein- protein complexes at 0 to 100 ns whereas Fig (b) represents RMSF analysis of the residues of the native and mutant CYLD protein with ubiquitin over the 100 ns simulation.)

Furthermore, to determine the structural flexibility of the protein-protein complexes, we assessed the RMSF value **(Fig 7).** This study revealed that R830K and native both complexes exhibited almost similar level of flexibility during the 100 ns simulation. However, some greater residual fluctuations also observed in case of R830K when compared with wild protein. The highest residual fluctuation for R830K was 8.22 Å observed at position GLN316 (899aa of CYLD). On the contrary, H827R exhibited highest residual fluctuation 9.66 Å at position LYS179 (762aa of CYLD) when compared to native and R830K complexes. Average fluctuation rate for native, mutant R830K and H827R were ∼ 2.041 Å ∼ 2.085 Å and ∼ 2.425 Å respectively. In terms of total residual portions, the RMSFs value of all mutant complexes differed significantly from the native complex structure.

From the radius of gyration (Rg) analysis, compactness and rigidity condition of protein-protein complex were determined. The Rg values of native protein complex ranged from 25.27 Å to 26.38 Å. In case of H827R and R830K, it ranged between 25.17 Å to 26.71 Å, and 24.849 Å to 25.99 Å respectively. The average value for the native structure, H827R and R830K mutants were ∼ 25.65 Å, ∼ 25.70 Å and ∼ 25.37 Å respectively **(Fig. 8).** It was observed that H827R complex had higher radius of gyration value than native and mutant R830K complexes thus showing least compactness.

**Fig. 8:**
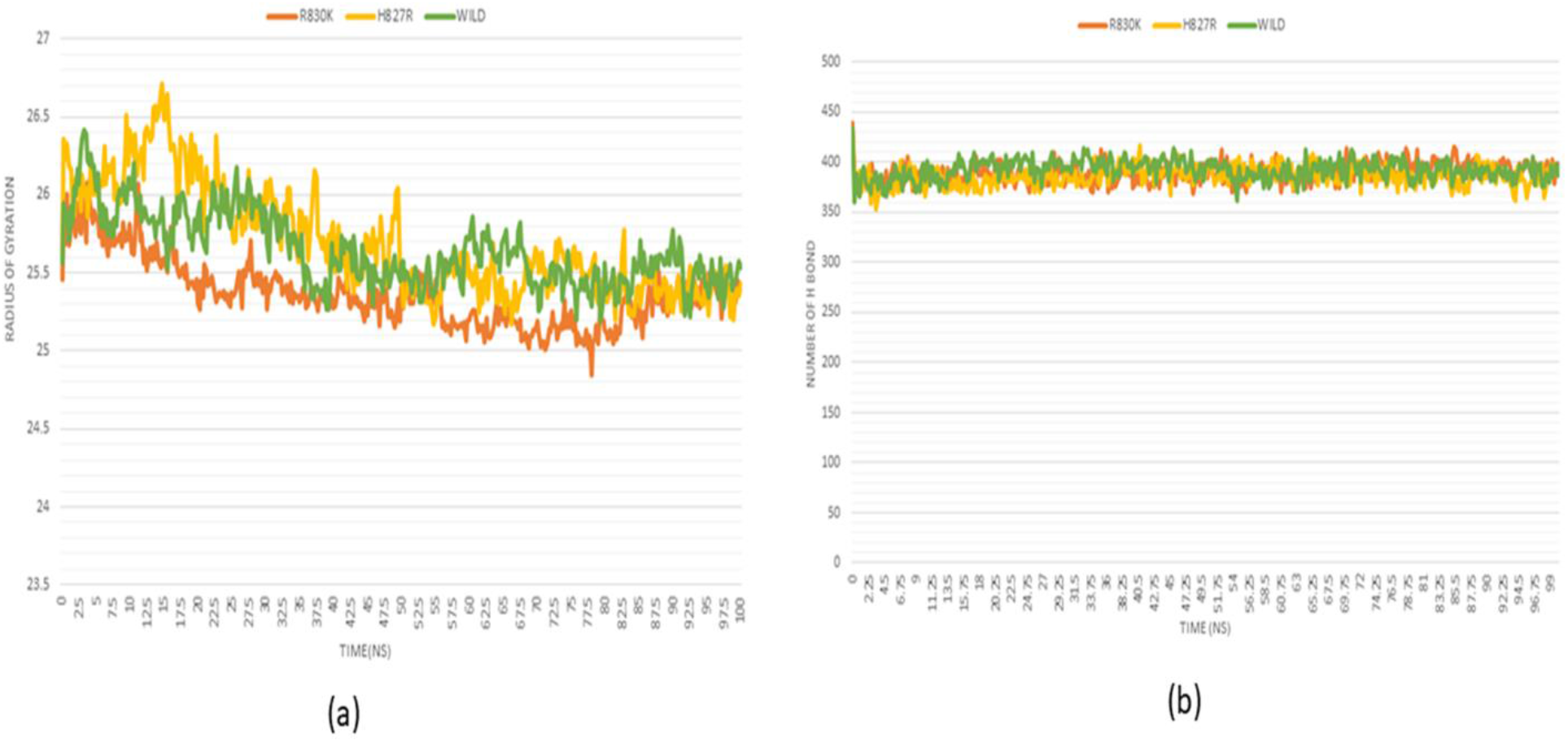
**Molecular dynamic simulation analysis performed by YASARA. (**Fig (a) shows Rg analysis of the backbone structure of the protein-protein complexes over 100 ns and Fig (b) indicates H-bond analysis of the structure of protein-protein complexes over 100ns.)

Following that, we studied the overall number of intra-molecular hydrogen bonds present in the protein to assess the protein stability or the stability between proteins. Native complex of CYLD- ubiquitin displayed an average of ∼ 392 H-bond throughout the 100 ns simulation. The average number of H-bond generated by CYLD-ubiquitin in mutant complexes R830K and H827R was estimated to be ∼ 389 and ∼ 386 respectively during the period of 100 ns simulation **(Fig. 8).** This analysis significantly depicts the impact of amino acid substitution on the backbone structure of the protein-protein complexes.

### Principal component analysis

The principal component analysis model was constructed based on the analysis of the various structural and energy profile such as bond distances, bond angles, dihedral angles, planarity, van der Waals energies, and electrostatic energies getting from MD simulation analysis. Three training sets were taken into consideration for the further analysis. Both the first and second principal components (PC1 and PC2) of this PCA model cover total 88.4% of the proportion variance. The score plot exhibits three different clusters for wild type-ubiquitin complex (green), mutant H827R-ubiquitin complex (blue) and mutant R830K-ubiquitin complex (red) where PC1 covers 66.7% and PC2 covers 21.7% of the variance **(Fig. 9**). Different cluster formation for three training sets signifies fluctuations that occurred during MD simulation. Significant fluctuations were observed when wild type is compared with both mutant types. This analysis indicated that point mutations directed to the alterations of the structural and energy profile of the CYLD-ubiquitin complexes. Therefore, mutations in the 827^th^ and 830^th^ position of the CYLD resulted in the aberrant interaction pattern of the CYLD with ubiquitin.

**Fig. 9:**
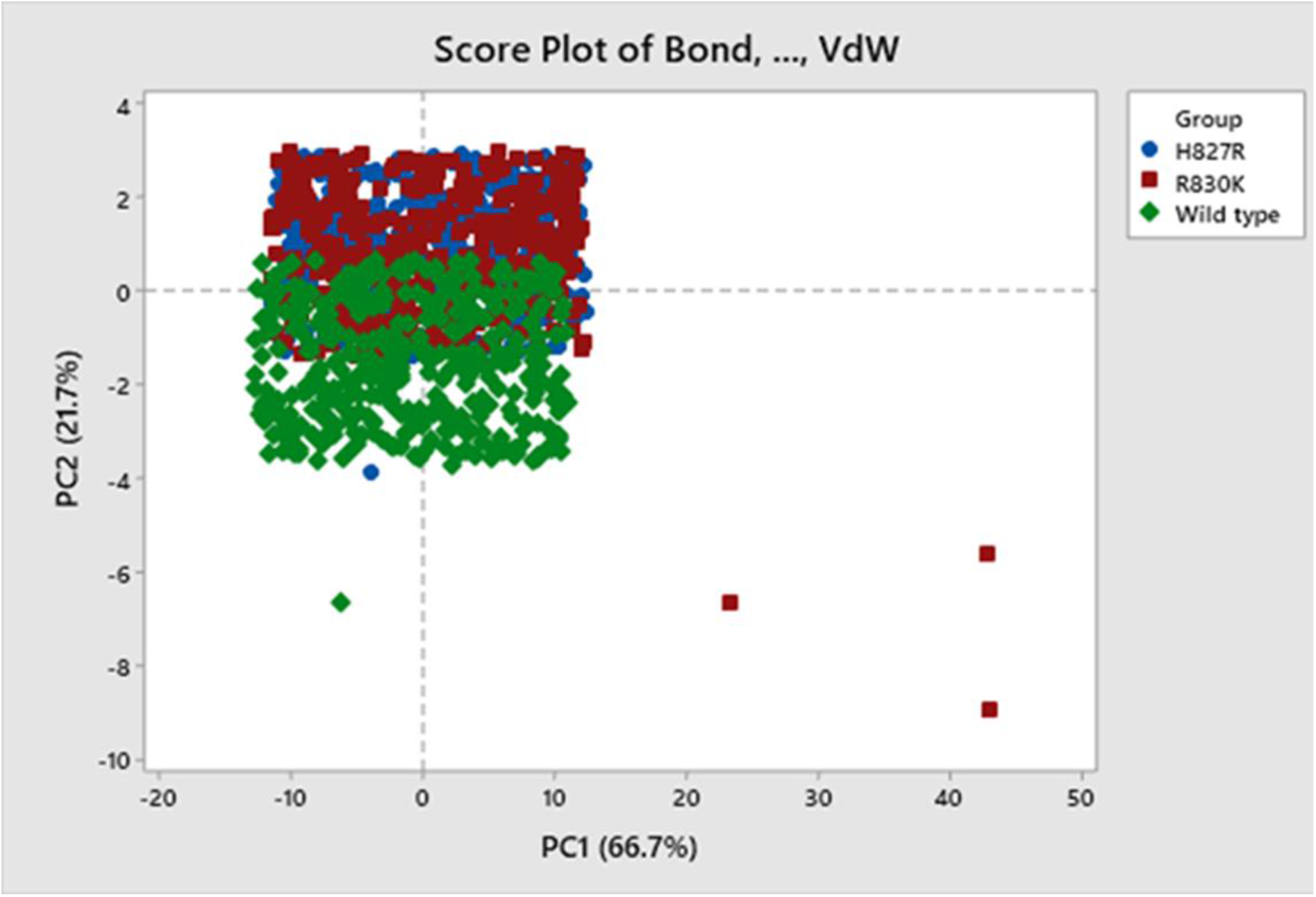
**Principal Component Analysis. (**The PCA model generated score plot consists of three different clusters: wild type CYLD-ubiquitin complex (green), mutant (H827R) CYLD-ubiquitin complex (blue) and mutant (R830K) CYLD-ubiquitin complex (red) where each dot indicates one time point.)

### Protein-protein network analysis

The functional interaction pattern of CYLD protein with other proteins in different biological pathways was predicted using the STRING database **(Fig. 10)**. CYLD functionally interacts with TRAF2, TRAF6, IKBKG, RNF31, TNFRSF1A, DDX58, RIPK1, BIRC3, UBC, UBE2K, UBE2S, RPS27A, UBA52, RAD18, RPL8, RPS16, RPL19, RPL35, RPS12, and RPL18A and thereby play various significant biological roles (**S9 Table).** This interaction pattern of CYLD may be disturbed if any deleterious change occurs in CYLD protein. The data about degree of connectivity, average shortest path length, betweenness centrality, closeness centrality of all the related protein of CYLD were predicted by Cytoscape (**S8 Table)**. Highest number of interactions was seen with both UBC (Ubiquitin C) and UBA52 (Ubiquitin A-52 residues ribosomal protein fusion product 1) with degree 20. Mutation can hamper all those interactions and therefore highlighted the deleterious effect of nsSNPs of CYLD.

**Fig. 10:**
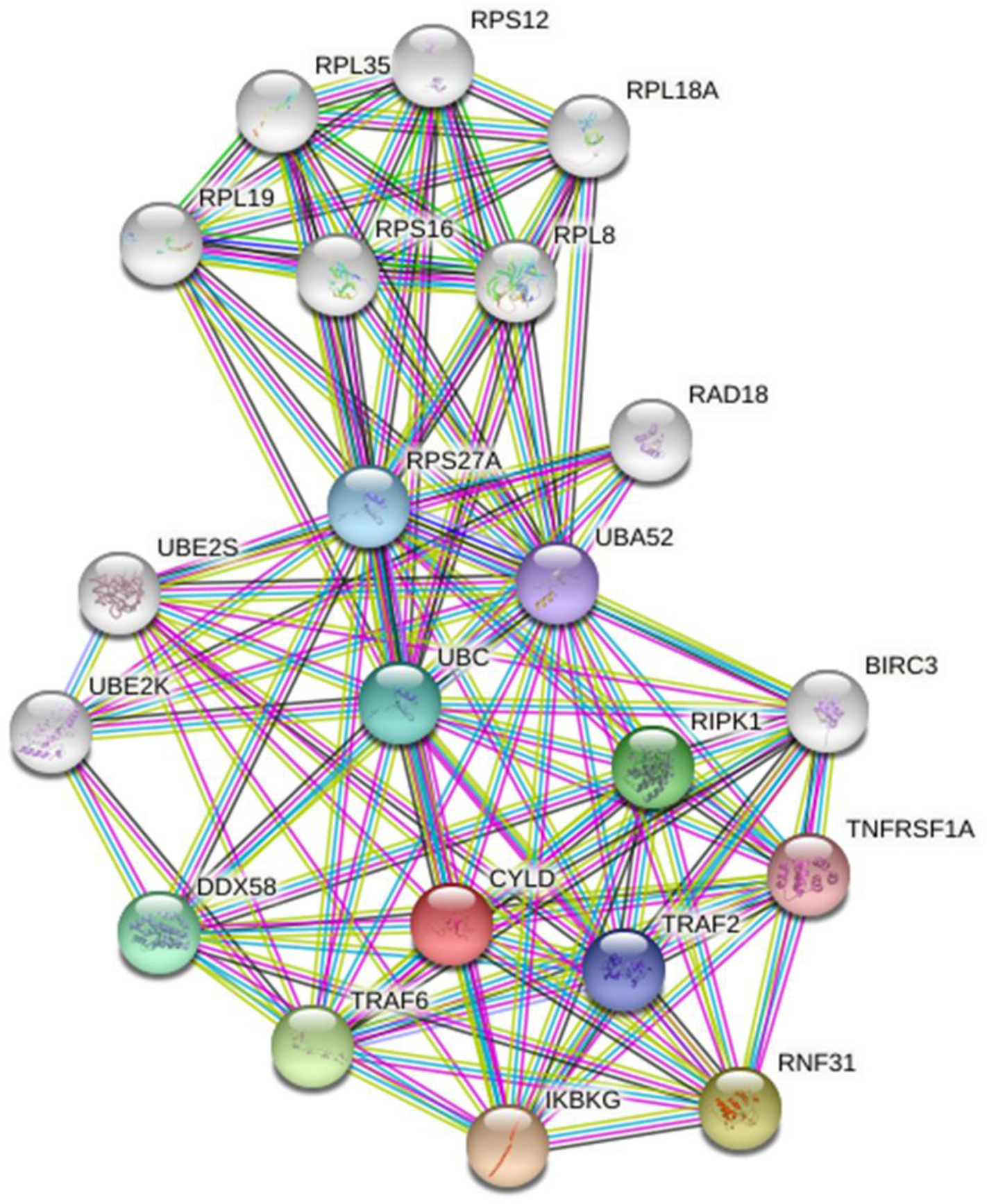
Protein-protein interaction network of CYLD protein constructed by the STRING database

## Discussion

CYLD known as a deubiquitinase gene that exhibits tumor suppression activity in humans (2). Mutation in CYLD is generally associated with many cancer types such as familial cylindroma, melanoma, salivary gland tumor, breast cancer, etc (2). Investigation of the impact of point mutation on the structural and functional activity of CYLD protein is a difficult task. Application of various bioinformatics tools make this analysis easier. In this study, we exploited the damaging consequences of nsSNPs of CYLD to study the effect on its structure and function using different computational approaches. We started our analysis by retrieving 446 nsSNPs recorded in NCBI database for CYLD gene. Subsequently, we examined these nsSNPs using seven different computational methods: PANTHER, PROVEAN, PREDICT SNP, POLYPHEN 2, PHD SNP, PON P2, SIFT for the screening out of high risk nsSNPs. Each algorithm ranked nsSNPs based on their deleterious effect taking into consideration parameters such as sequence homology, structural homology, conservancy, biological and physical characteristics of amino acids. The integration of different algorithms often serves as powerful tools to prioritize the functional SNP candidates (65). Considering this, we focused on 18 significant nsSNPs of CYLD commonly predicted as deleterious by all the seven tools. InterPro, domain identification program revealed that two nsSNPs were located at Cap-Gly domain required for the interaction with NEMO/IKKγ and TRAF2 (17) whereas rest of the nsSNPs were positioned on the USP domain responsible for its deubiquitinase activity (18) . A study reported that mutation in conserved regions lead to the greater reduction in protein stability compared to non-conserved regions (66) . Therefore, we analyzed the conservation profile of our targeted nsSNPs from there we only considered highly conserved residues with the help of Consurf server. Several studies demonstrate that alteration in protein stability due to SNP can cause degradation, misfolding, and coagulation in a protein leading to structural and functional impairments (67, 68) and we have found 14 destabilizing residues among 18 nsSNPs in our study when we used I-mutant and SDM tools .

Next, we approached to determine the profile of structural modifications caused by these destabilizing nsSNPs through the comparative structural analysis for both native and mutant protein models using Phyre 2 homology model prediction server. TM align tool determined the structural deviations of mutant models in comparison with the native protein. According to studies (46, 69) TM-score determines the topological similarity whereas RMSD indicates average distance between α-carbon backbones of wild type and mutant proteins .Greater RMSD value signifies greater deviation and lower TM score means higher dissimilarities between wild and mutant protein models. We furthered 13 nsSNPs based on higher RMSD value and lower TM score and we found in one study (70) that they also selected nsSNPs based on higher RMSD. In our study, among 13 nsSNPs, highest RMSD value (2.21) was found in highly conserved H827R, and lowest TM score (0.84714) was displayed by highly conserved R830K. I-TASSER generated confidence score by remodeling more reliable wild type and mutant type proteins. We also investigated relative terms in Swiss Model such as solvation, torsion, qmean, Cβ value comparing wild type and mutant models. Project Hope program provides deep insight on the detrimental effect of point mutation on the structural configuration of a protein. Analysis showed that wild type residues replaced by smaller mutants result in the empty space formation due to the loss of significant interactions in case of R830K. Besides, misfolding and repulsion can cause when charge was added to H827R. The influence of deleterious nsSNPs on the energy minimization state of the CYLD protein determination is fundamental as protein achieve its stable conformation with lower energy after energy minimization according to a study (71). On the contrary, structural changes due to mutation can restraint the protein to be stable easily. Findings showed that the total energy for the native CYLD protein was -20130.191 kj/mol after energy minimization. H827R mutant showed remarkable increase in energy -15956.584 kj/mol than wild type.

Furthermore, we performed molecular docking between CYLD PDB id:2VHF (583aa-956aa) and ubiquitin as binding interactions pattern among them has significant role in tumor suppressor activity of CYLD (10). Studies showed that decreasing binding affinity due to mutation signifies impairment of the binding interaction pattern (72, 73). Similarly, our analysis also revealed that 4 nsSNPs: L781P, P698T, H827R and R830K mutant complex showed lowest binding affinity of -13.4, -13.0-12.8 and -12.7 kj/mol respectively when compared with wild type (-14.6 kj/mol). We observed higher dissociation constant for these 4 nsSNPs (**S6 Table**) compared to the native CYLD which also substantiated weak binding interactions between ubiquitin and CYLD mutant protein complex. Mutation 3D specifically verified that mutation in H827 and R830 can have strong association with cancer whereas no association was found for L781P and P698T.

We performed MD simulations to evaluate the dynamic behavior of our CYLD-ubiquitin complex in an aqueous environment for 100 ns. Simulation was executed with a time step interval of 2.5 femtoseconds (74, 75). This analysis mainly focused on the relative structural deviation of the H827R and R830K in comparison to wild type CYLD protein. Mentionable variations in RMSD (root mean square deviation) value were observed in H827R and R830K compared with the wild type protein. Wild type CYLD exhibited variations in RMSD value up to 10.5 ns and then became stable within range between 2.9 to 4.4 Å during the simulation time frame. In case of H827R, we found highest peak at 80 ns with RMSD value 6.275 Å indicated that mutant H827R became unstable throughout the whole simulation period. On the other hand, we found highest peak at 71ns with RMSD value 6.087 Å in case of R830K. Average RMSD value for mutants H827R (4.9575 Å) and R830K (5.278 Å) were much higher than the native CYLD (3.388 Å). These results indicated that H827R and R830K lead to the structural variation of the CYLD protein as higher RMSD value signifies structural distance of protein or protein complex. After that, we analyzed root-mean-square fluctuation (RMSF) of CYLD and its two mutants to evaluate mutation-causing fluctuations in structural part of a protein comparing with the actual structure of a protein. We observed higher residual fluctuation in H827R rather than R830K when referenced with native CYLD. We found highest RMSF 9.66 Å at positions LYS179 (762aa of CYLD) for H827R. In case of R830K, higher RMSF value 8.22 was found at 316 (899 aa) residue.

Rg (Radius of gyration) analysis determined the compactness of CYLD protein thereby signified the folding rate as well as stability of that protein. We found that Rg values of CYLD protein complex ranged from 25.27 Å to 26.38 Å where as in case of H827R and R830K, Rg ranged between 25.17 Å to 26.71 Å and 24.849 Å to 25.99 Å respectively. H827R mutant showed higher Rg value and R830K showed less Rg value compared with wild type. From this we can hypothesize that, the compactness of the CYLD protein is probably affected by mutation at position 827 than at 830. Finally, H-bond analysis of CYLD was performed. A study revealed that if any alteration occur in H-bond formation, it can affect the folding and stability of a protein (76). Average H-bond of native CYLD-ubiquitin displayed ∼ 392 H-bonds whereas the average H-bond generated by CYLD-ubiquitin in mutants R830K and H827R was calculated to be ∼ 389 and ∼ 386 respectively. Loss of H-bond in mutant complex signified its weak binding interaction with ubiquitin as well as its structural deformation. We found several studies (70, 77–79) where they did not performed molecular docking and MD simulations for observing changes in interaction pattern as well as stability of a protein after point mutation. Principal component analysis obtained from MD simulation hints at the aberrant structural and functional activity of CYLD due to the point mutation at 827 and 830 position. In several studies (58, 80) they found greater deviation in structure and energy profile by comparing wild and mutants and we also found deviation in structure and energy profile by comparing wild type CYLD with H827R and R830K mutants. We also examined the interacting partners of CYLD in various biological pathways through network analysis and suggested that H827R and R830K mutants can disturb those pathways.

CYLD performs its tumor suppressor activity by disassembling k-63 ubiquitin chain (20, 81, 82) where interaction between C-terminal USP domain and ubiquitin chain is the prerequisite for this function. Mutation in USP domain can interrupt their Lys63-linked polyubiquitin cleavage activity resulting in cancer (1). In our current study, we tried to shortlist deleterious SNP disrupting the total catalytic activity and binding affinity of USP domain of CYLD and their strong association with cancer.

Throughout the study, a consistent workflow was developed for the reproducibility of this in silico deleterious SNPs prediction and multiple algorithms, tools were used to assess each step to increase the accuracy of the approaches by removing the artifacts from each tool. As SNPs in the genome are being considered as critical in regard to functional and structural effects of proteins involving cellular metabolism, gene expression and disease susceptibility etc. This computational prediction- based approach would provide deep insights and faster outcomes for experimental validation.

In conclusion, Mutation in tumor suppressor CYLD has been linked to a variety of cancers. Therefore, determination of the effect of point mutations on the structural and functional activities of the CYLD protein is a challenging task. The use of numerous bioinformatics tools simplifies this assessment. In our present study, we employed multiple computational tools to investigate the harmful consequences of the mutant variant of CYLD on its structure and function. As mutant CYLD associated with different cancer types, our results will be useful in the development of future diagnostic and research on CYLD mutations. Our findings however required in vitro and in vivo experimental validation.

## Acknowledgments

We acknowledge the Department of Biotechnology and Genetic Engineering and Computational Biology and Chemistry Lab, Noakhali Science and Technology University, for providing the research work support.

## Author’s Contributions

Conceptualization: N.M.B, M.R.A, H.A.R, S.G Data curation: A.S.R, T.F.

Formal analysis: A.S.R, T.F, M.A.M., D.A.S, N.J.A, M.K.I

Funding acquisition:

Methodology: N.M.B, M.R.A, H.A.R, S.G Writing – original draft: A.S.R, T.F, H.A.R

Writing – review & editing: M.R.A, N.M.B, M.S.H.

## Supporting Information

**S1 Fig:**
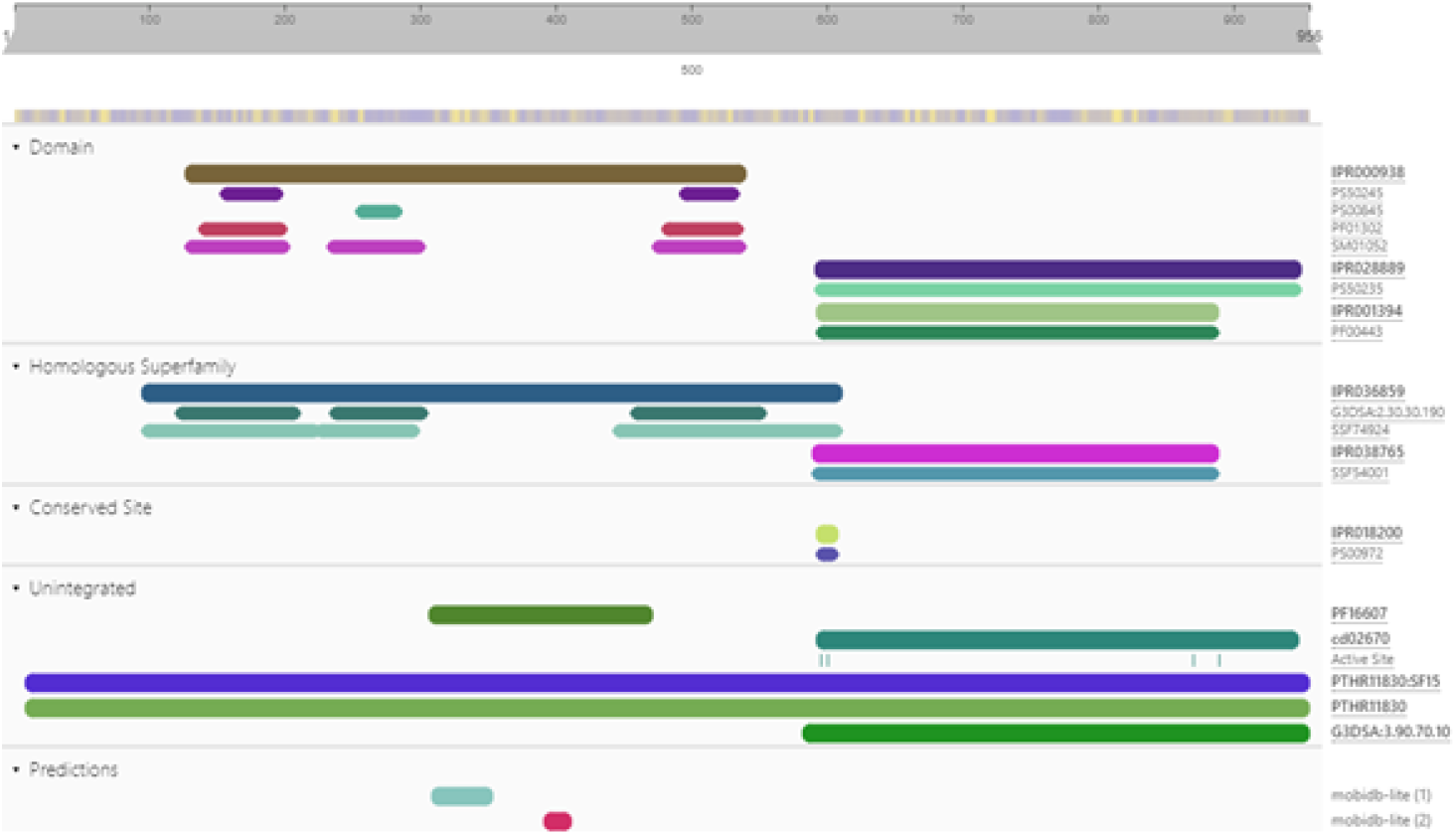
Domain identification of CYLD using InterPro server. CYLD contains two functional domains namely: Cap-Gly domain (127-540 Amino acid) and USP domain (592-950 AA).

**S2 Fig:**
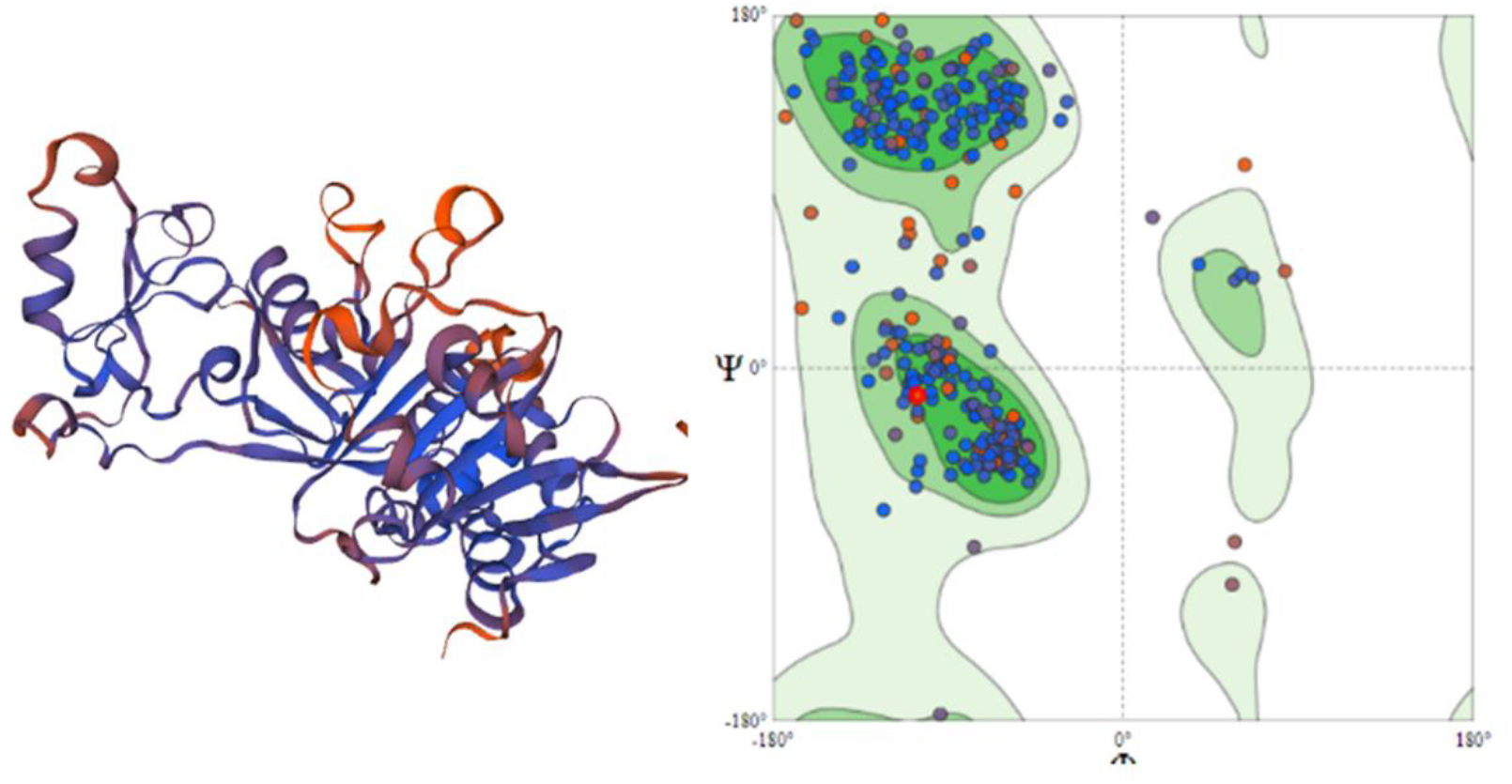
The 3D structure of CYLD protein (USP domain) constructed by SWISS MODEL and structural quality assessment using Ramachandran plot

**S1 Table:**
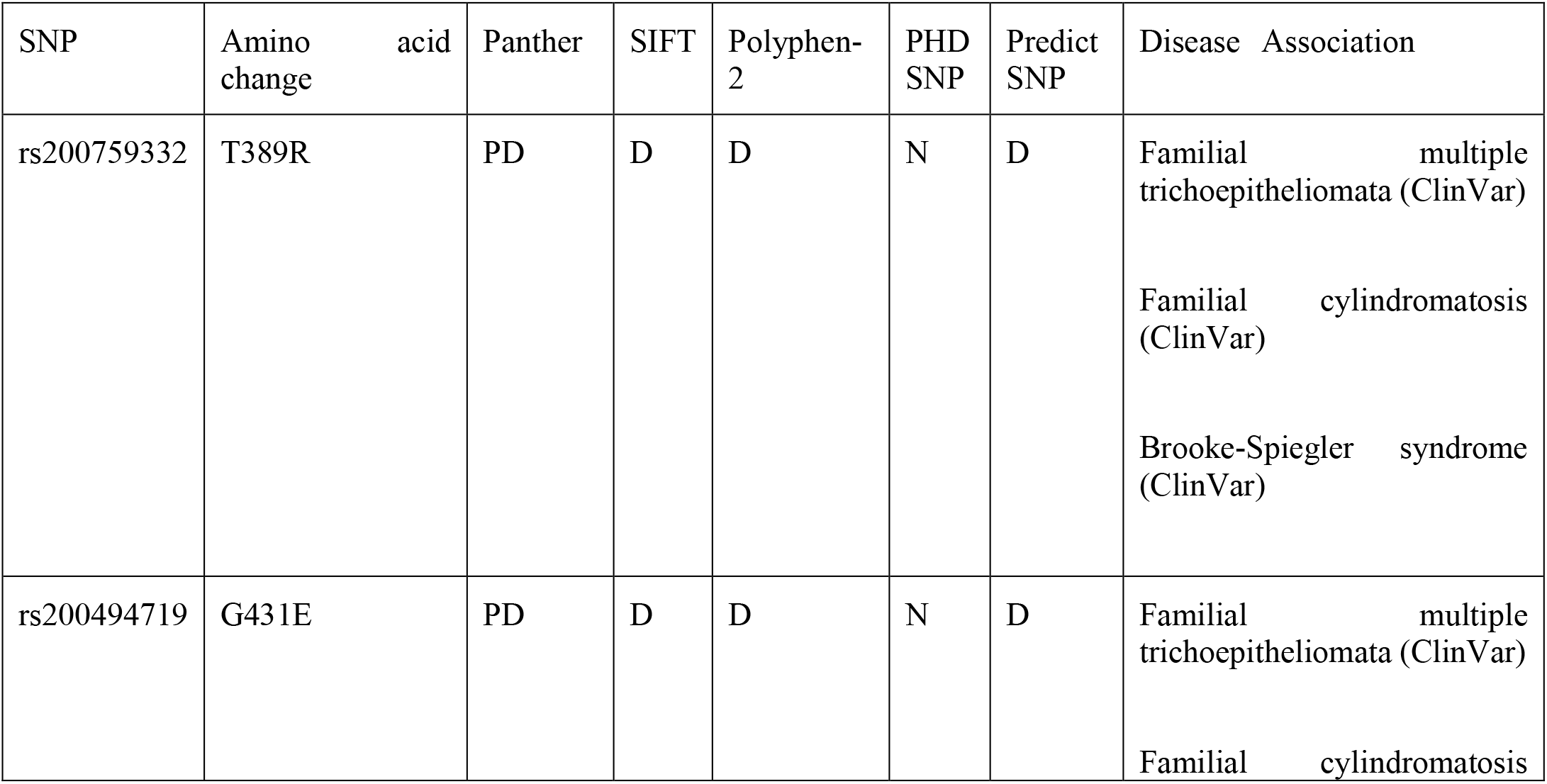
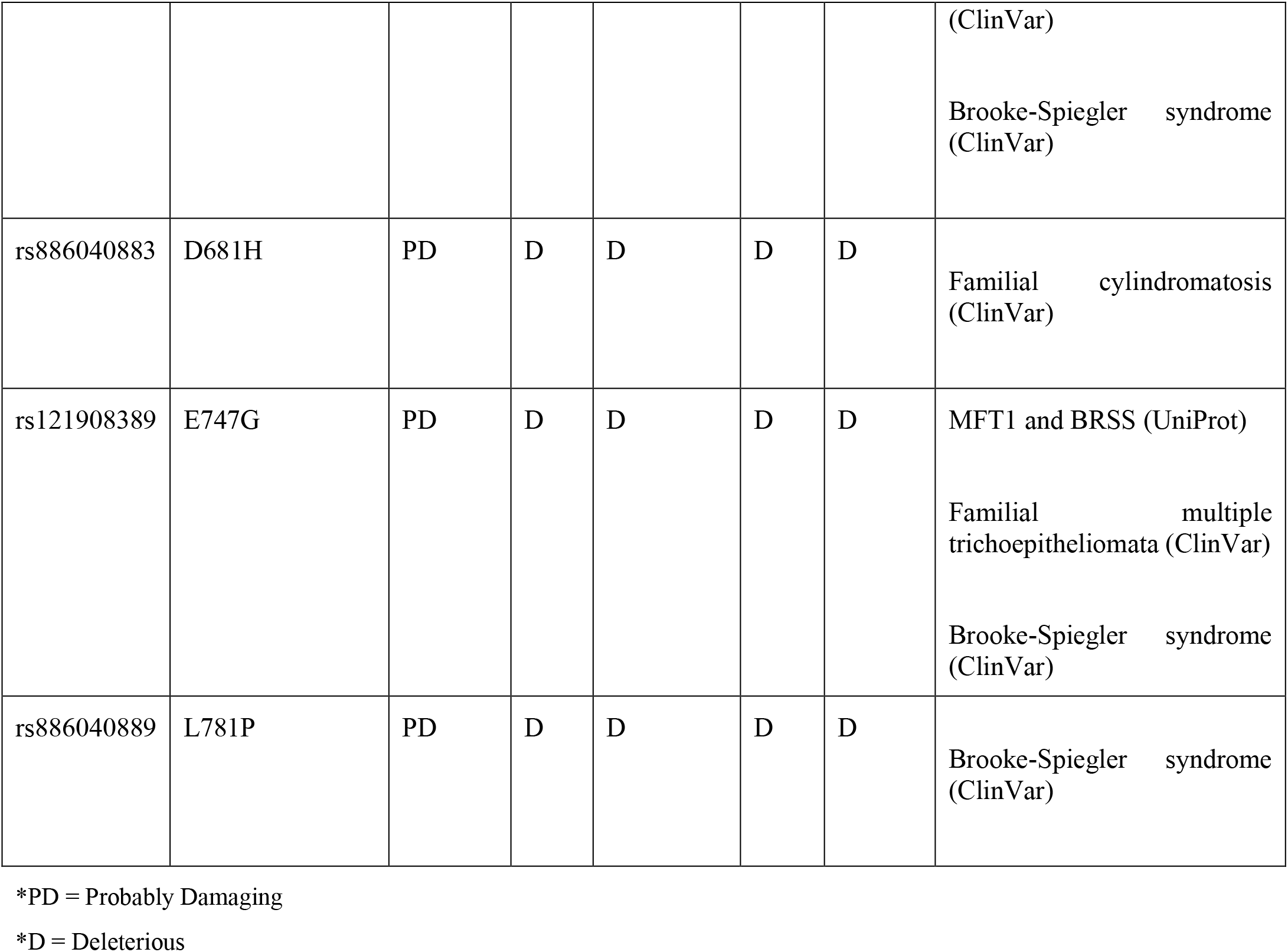
Prediction of known disease-associated variants.

**S2 Table:**
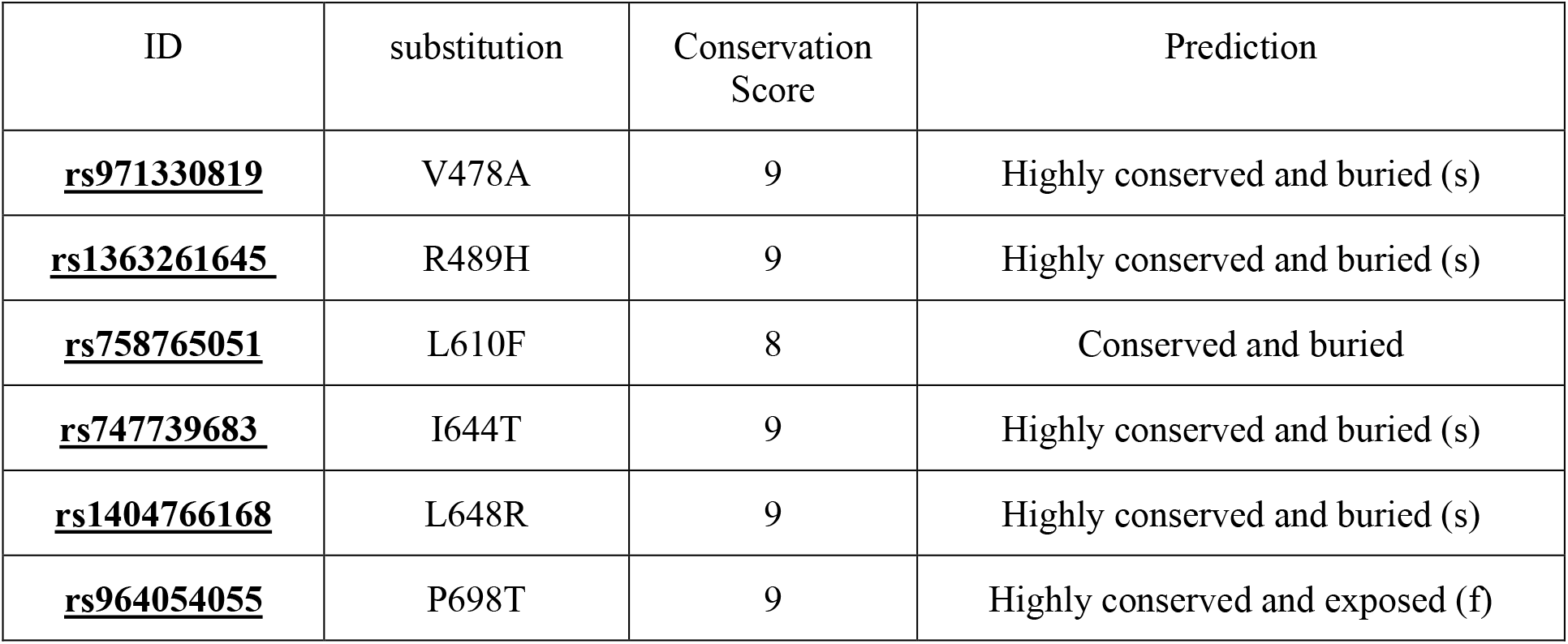
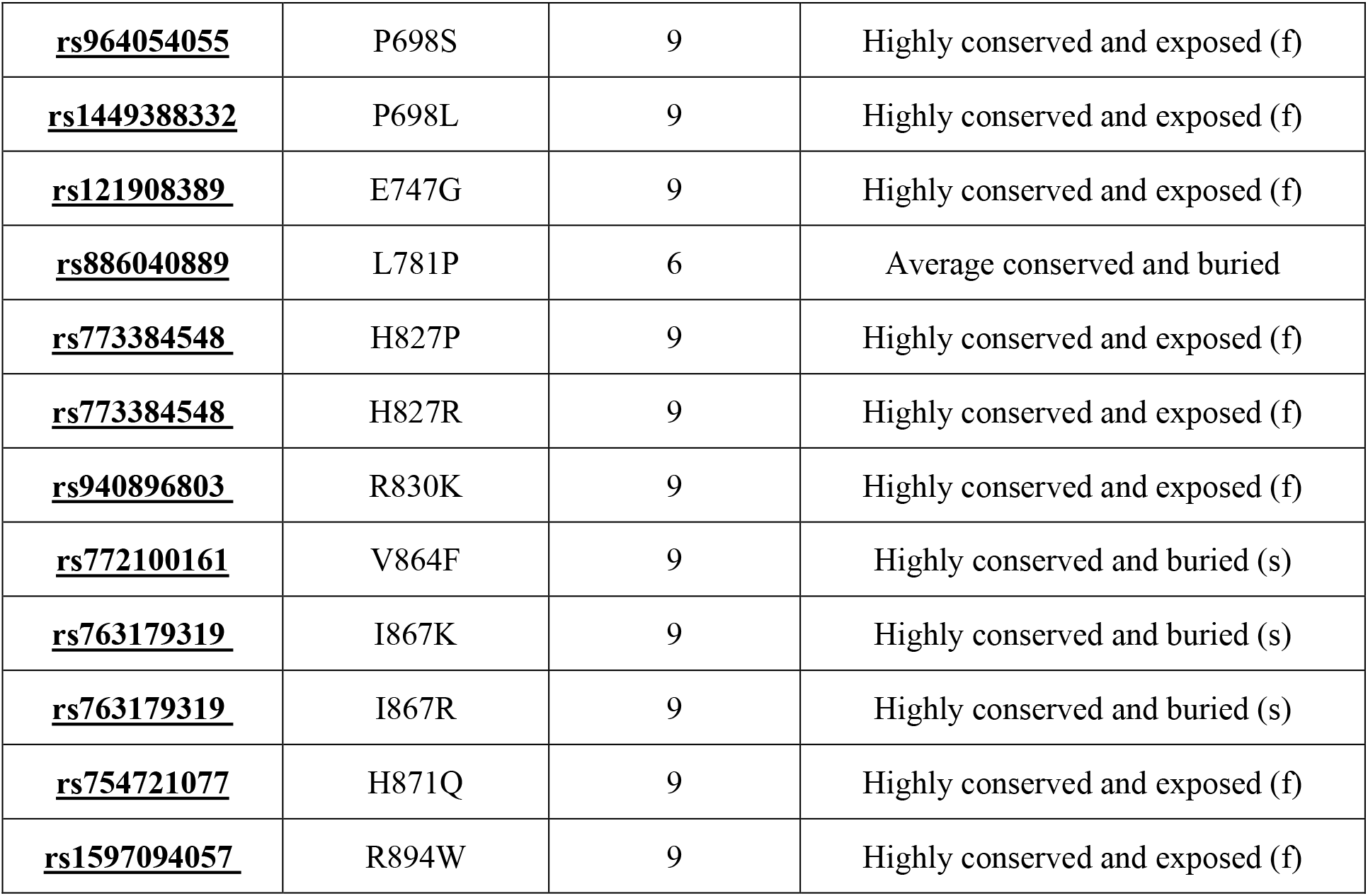
Conservancy analysis using Consurf server.

| ID | substitution | Conservation Score | Prediction |
| --- | --- | --- | --- |
| **<u>rs971330819</u>** | V478A | 9 | Highly conserved and buried (s) |
| **<u>rs1363261645</u>** | R489H | 9 | Highly conserved and buried (s) |
| **<u>rs758765051</u>** | L610F | 8 | Conserved and buried |
| **<u>rs747739683</u>** | I644T | 9 | Highly conserved and buried (s) |
| **<u>rs1404766168</u>** | L648R | 9 | Highly conserved and buried (s) |
| **<u>rs964054055</u>** | P698T | 9 | Highly conserved and exposed (f) |
| <u>**rs964054055**</u> | P698S | 9 | Highly conserved and exposed (f) |
| <u>**rs1449388332**</u> | P698L | 9 | Highly conserved and exposed (f) |
| <u>**rs121908389**</u> | E747G | 9 | Highly conserved and exposed (f) |
| <u>**rs886040889**</u> | L781P | 6 | Average conserved and buried |
| <u>**rs773384548**</u> | H827P | 9 | Highly conserved and exposed (f) |
| <u>**rs773384548**</u> | H827R | 9 | Highly conserved and exposed (f) |
| <u>**rs940896803**</u> | R830K | 9 | Highly conserved and exposed (f) |
| <u>**rs772100161**</u> | V864F | 9 | Highly conserved and buried (s) |
| <u>**rs763179319**</u> | I867K | 9 | Highly conserved and buried (s) |
| <u>**rs763179319**</u> | I867R | 9 | Highly conserved and buried (s) |
| <u>**rs754721077**</u> | H871Q | 9 | Highly conserved and exposed (f) |
| <u>**rs1597094057**</u> | R894W | 9 | Highly conserved and exposed (f) |

**S3 Table:**
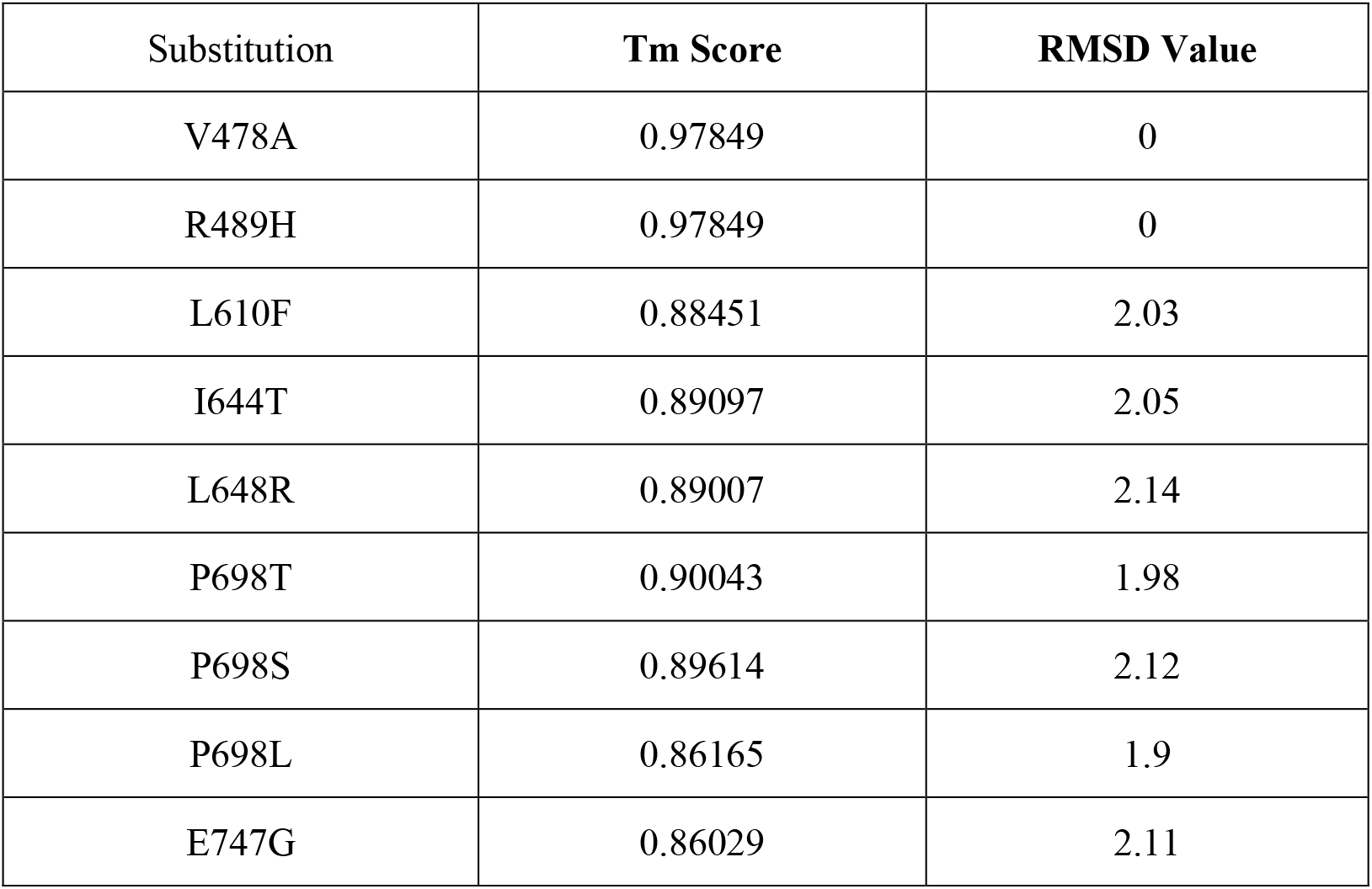
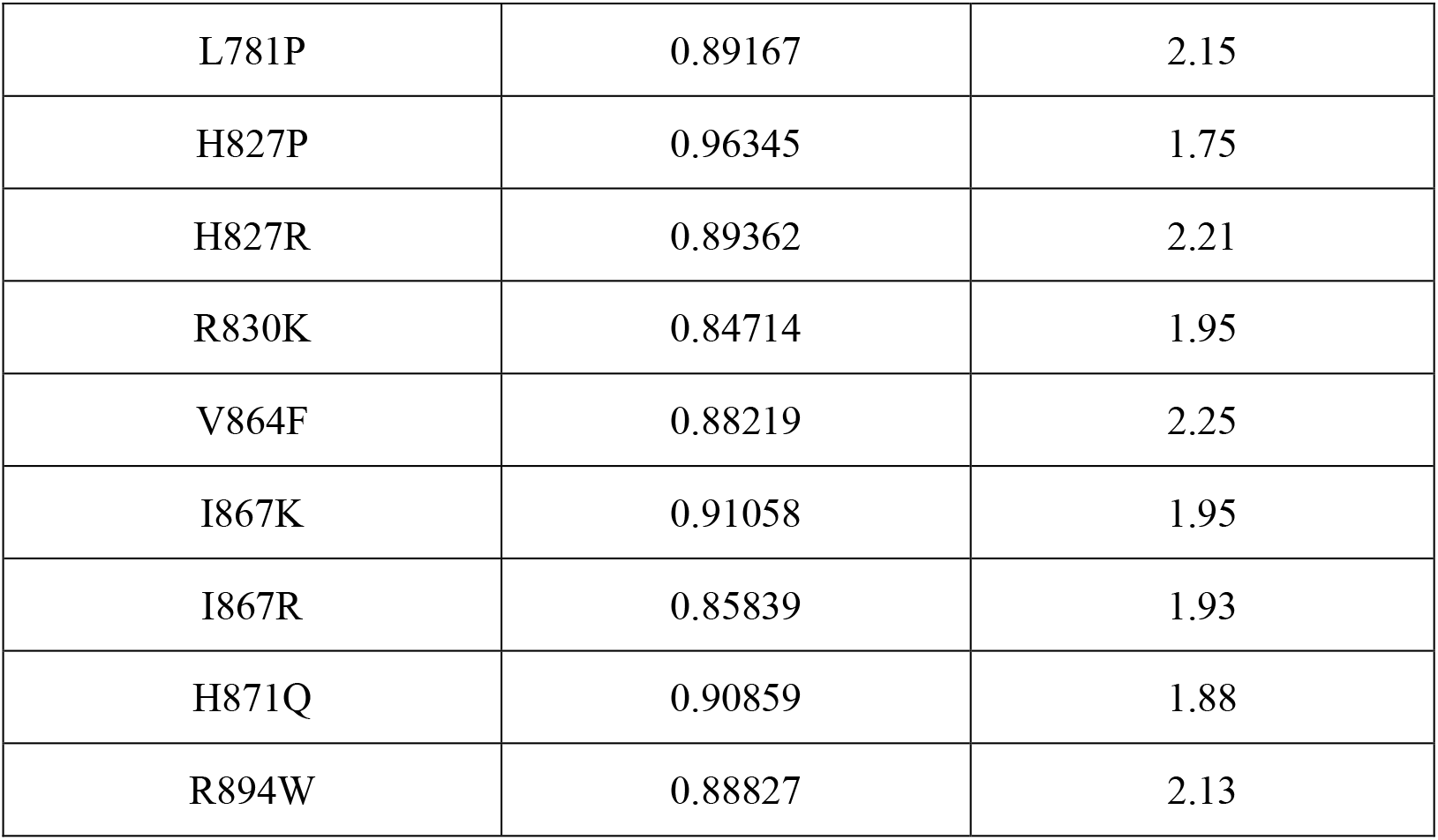
Tm score and RMSD value predicted by TM align tool.

**S4 Table:**
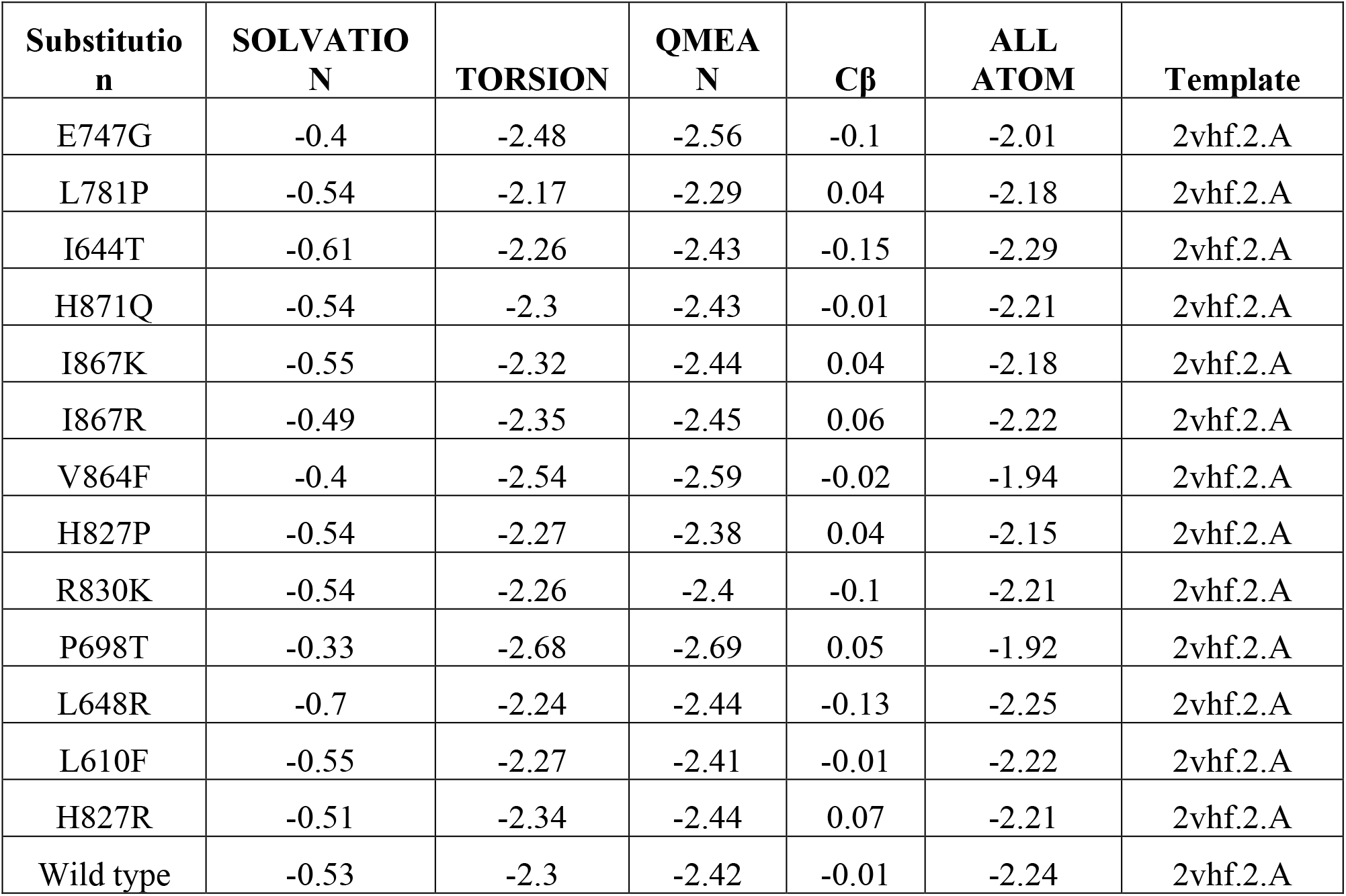
Swiss model Result.

**S5 Table:**
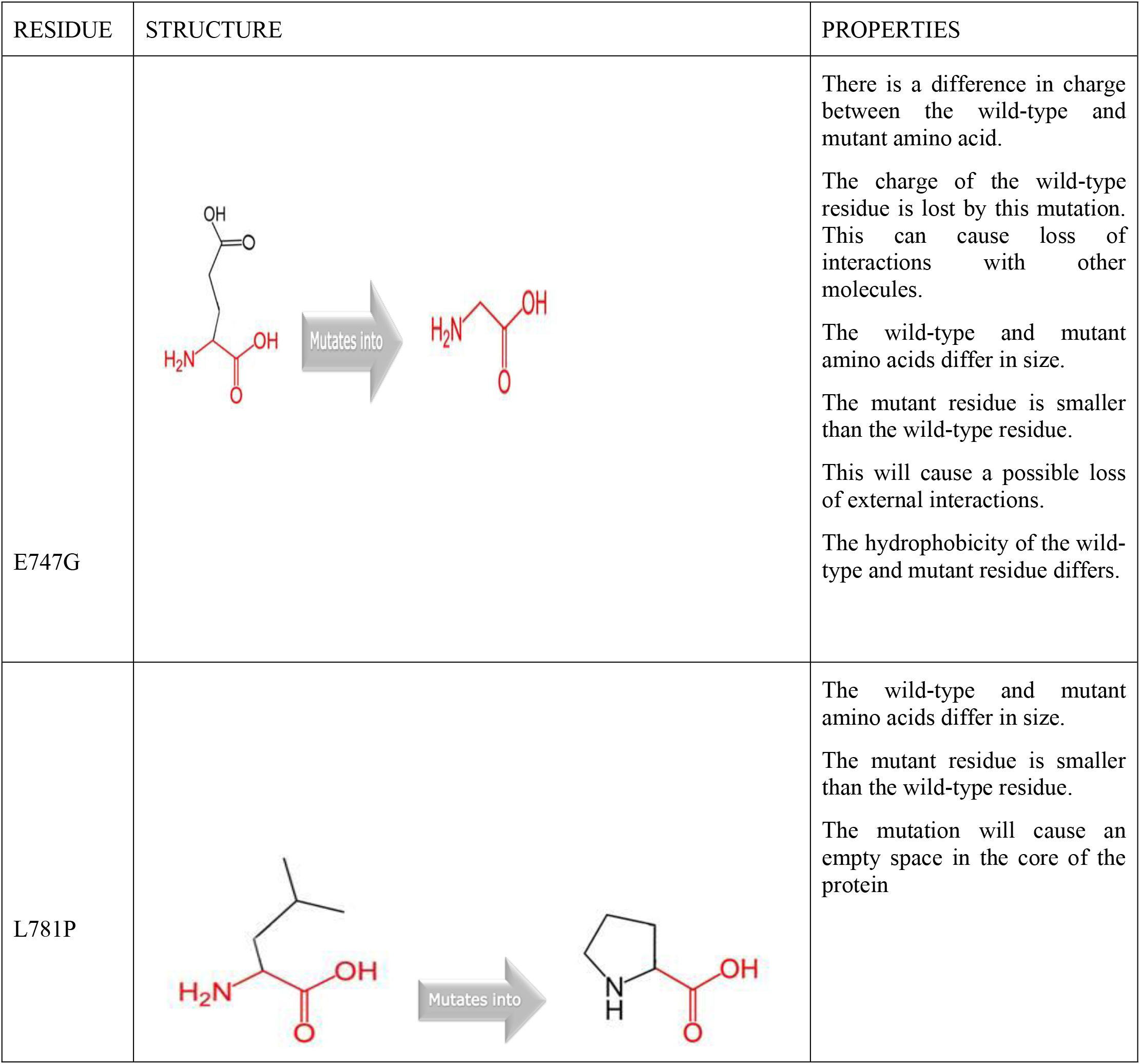
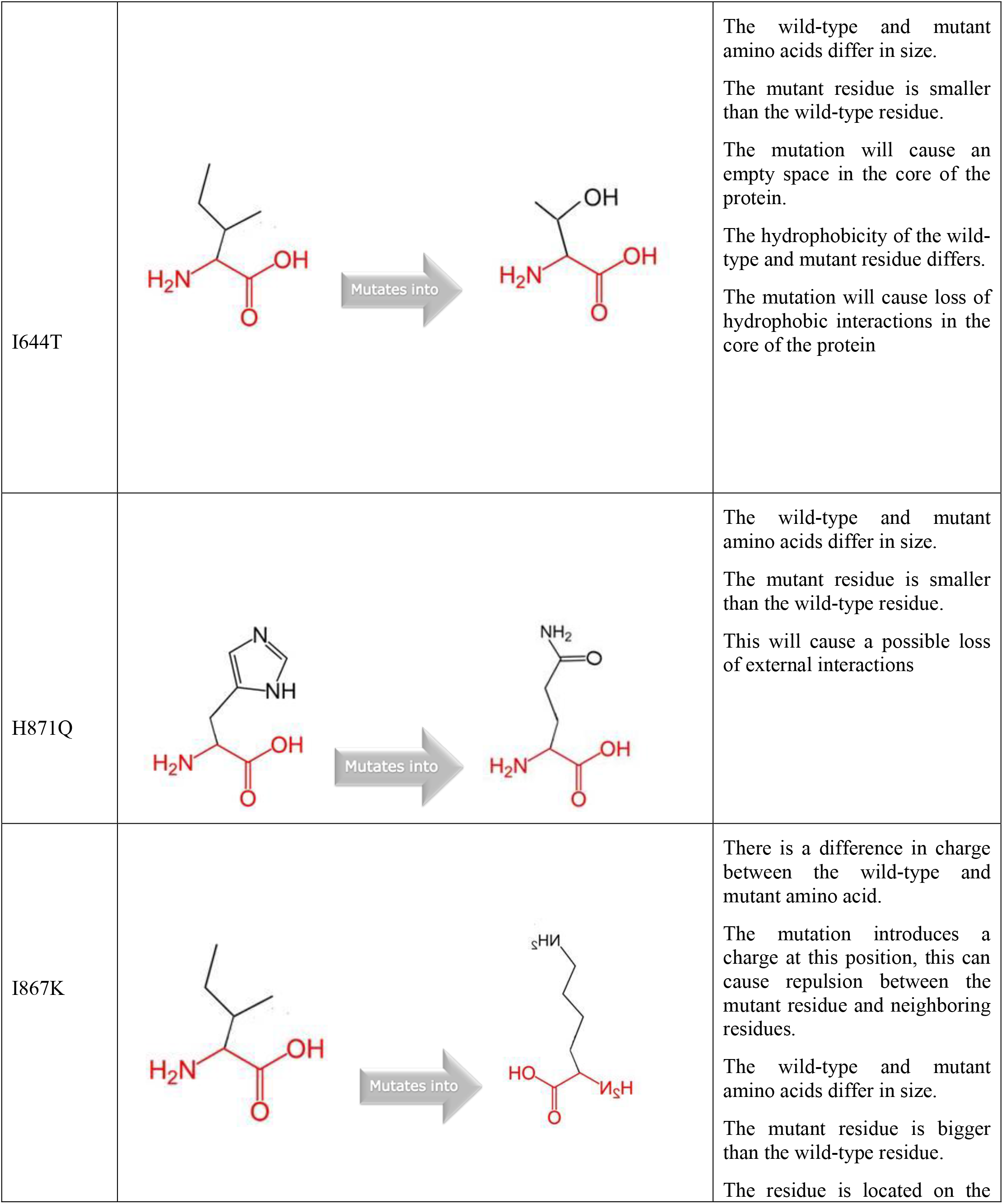
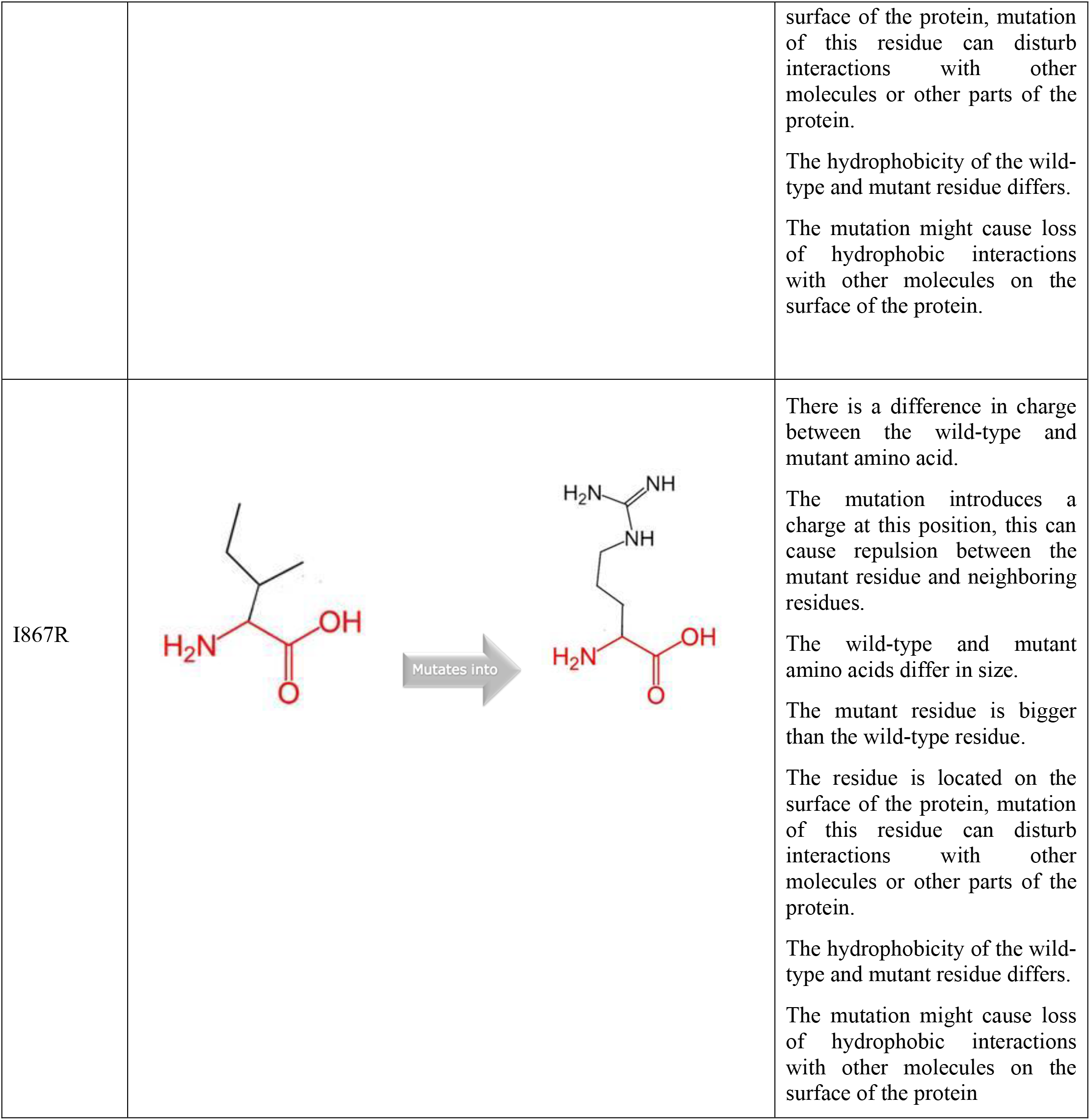
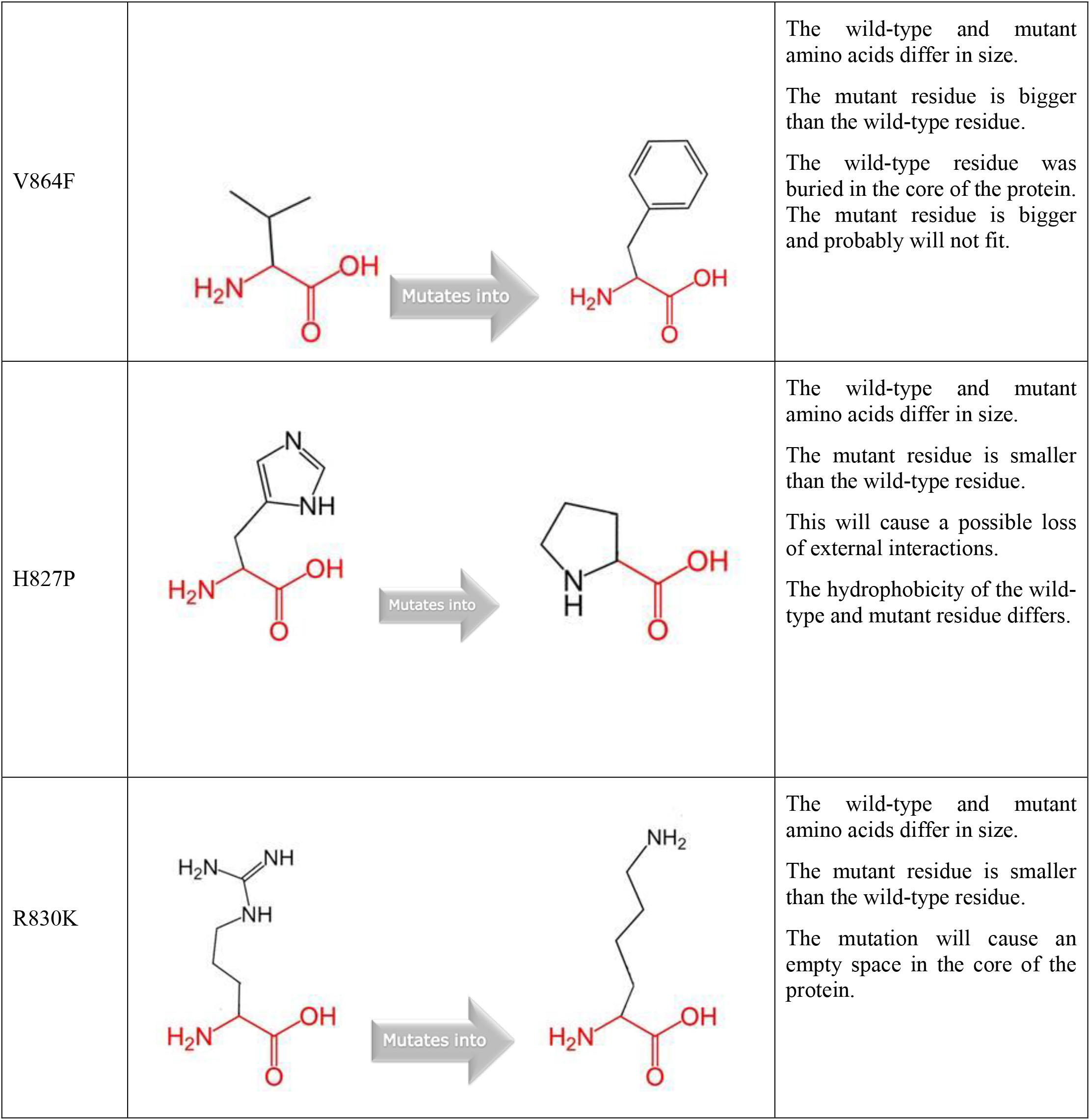
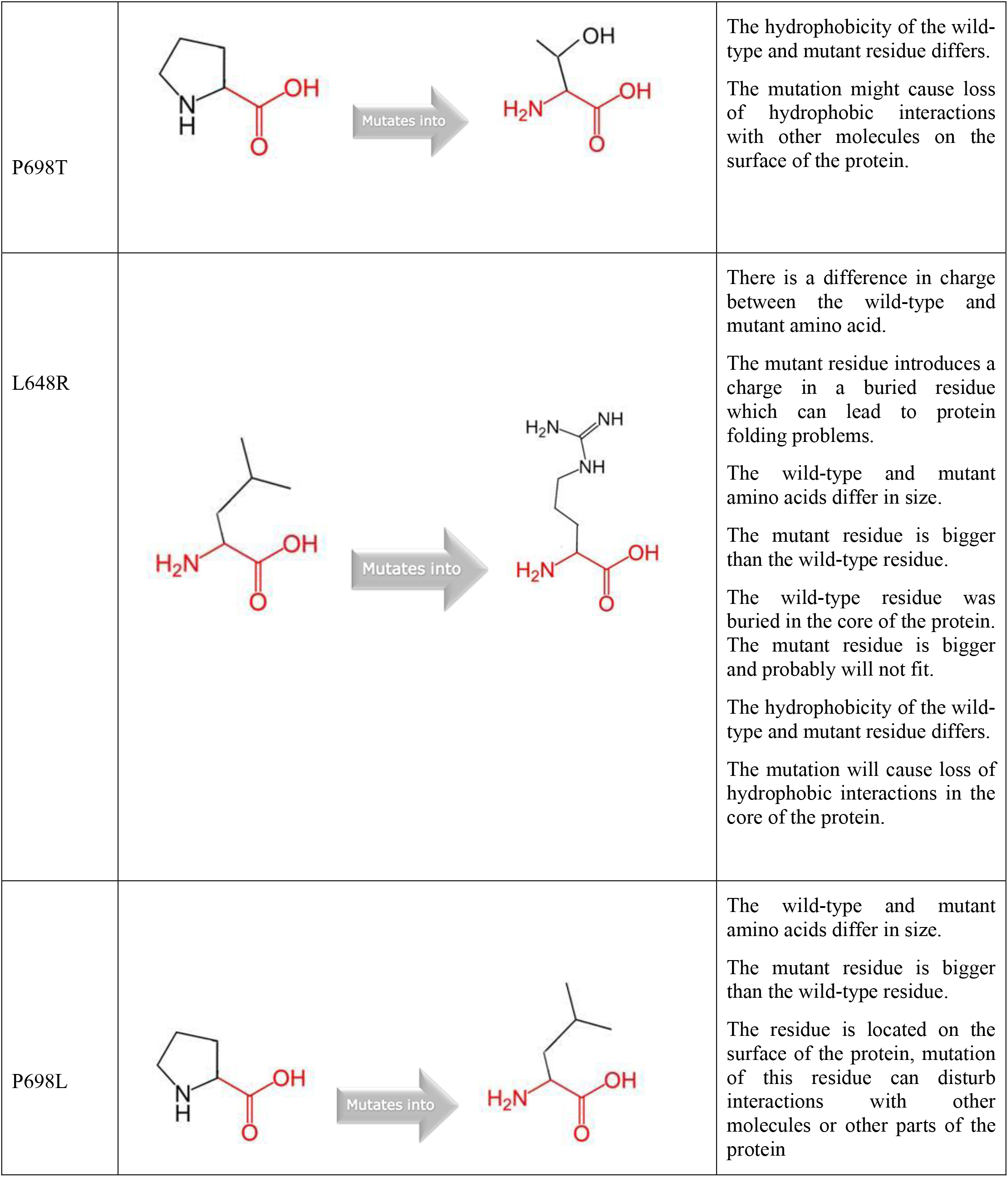
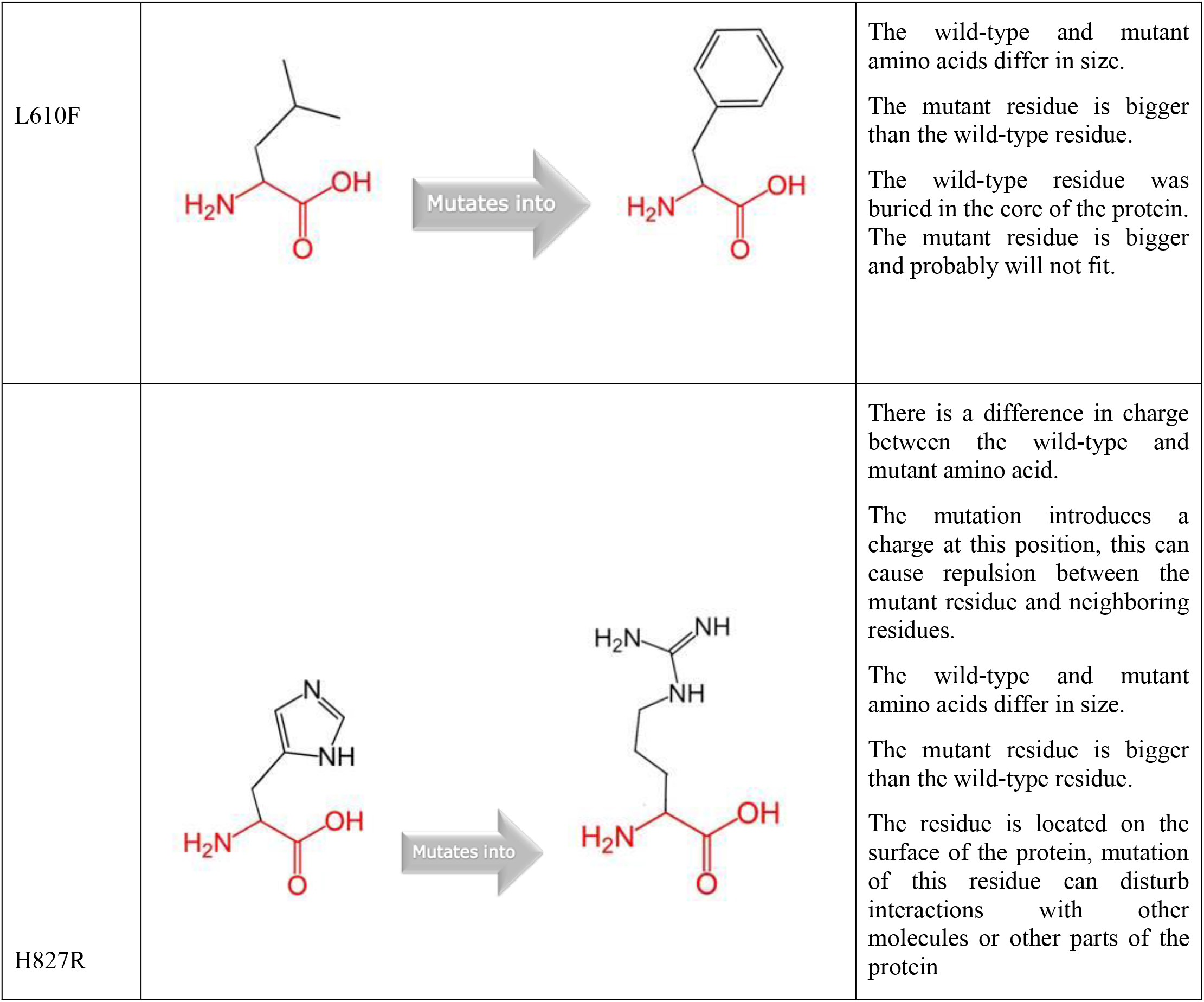
Structural effect of the point mutation predicted by HOPE project.

**S6 Table:**
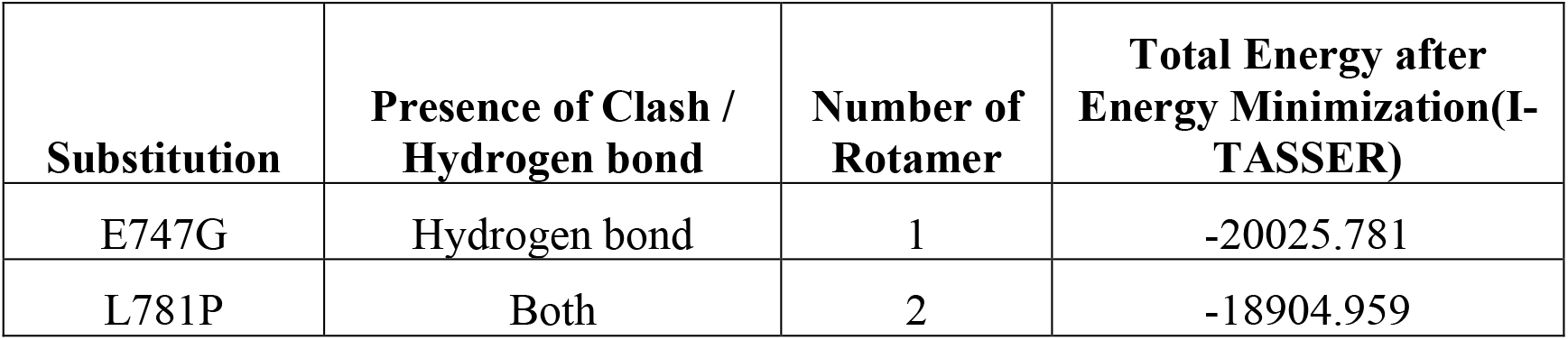
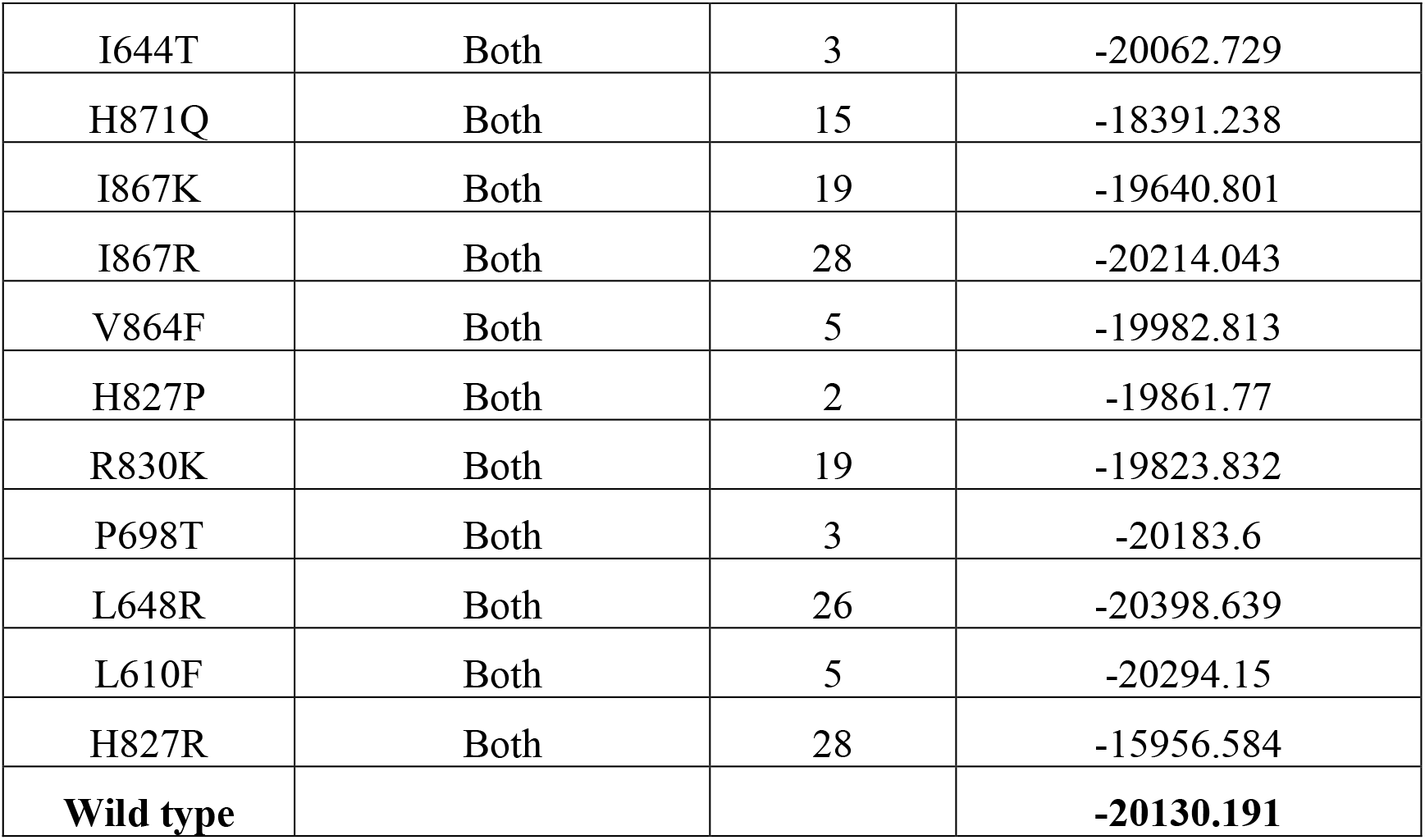
Energy minimization state of wild and 13 mutant variants determined by Swiss PDB Viewer.

**S7 Table:**
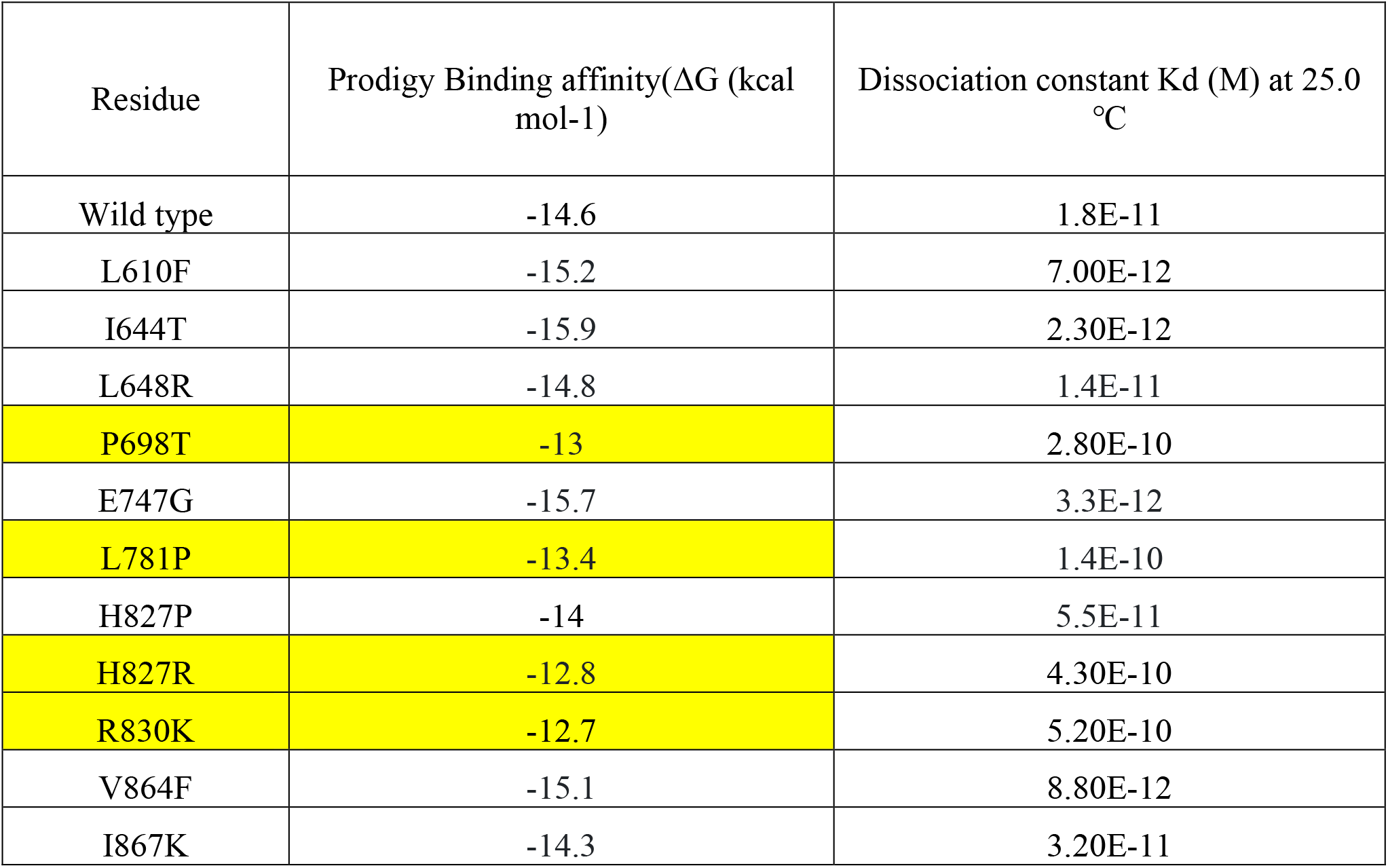
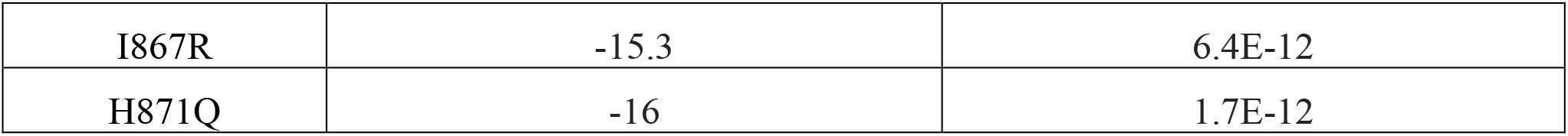
Binding affinity and dissociation constant of all docking complex shown in the table.

**S8 Table:**
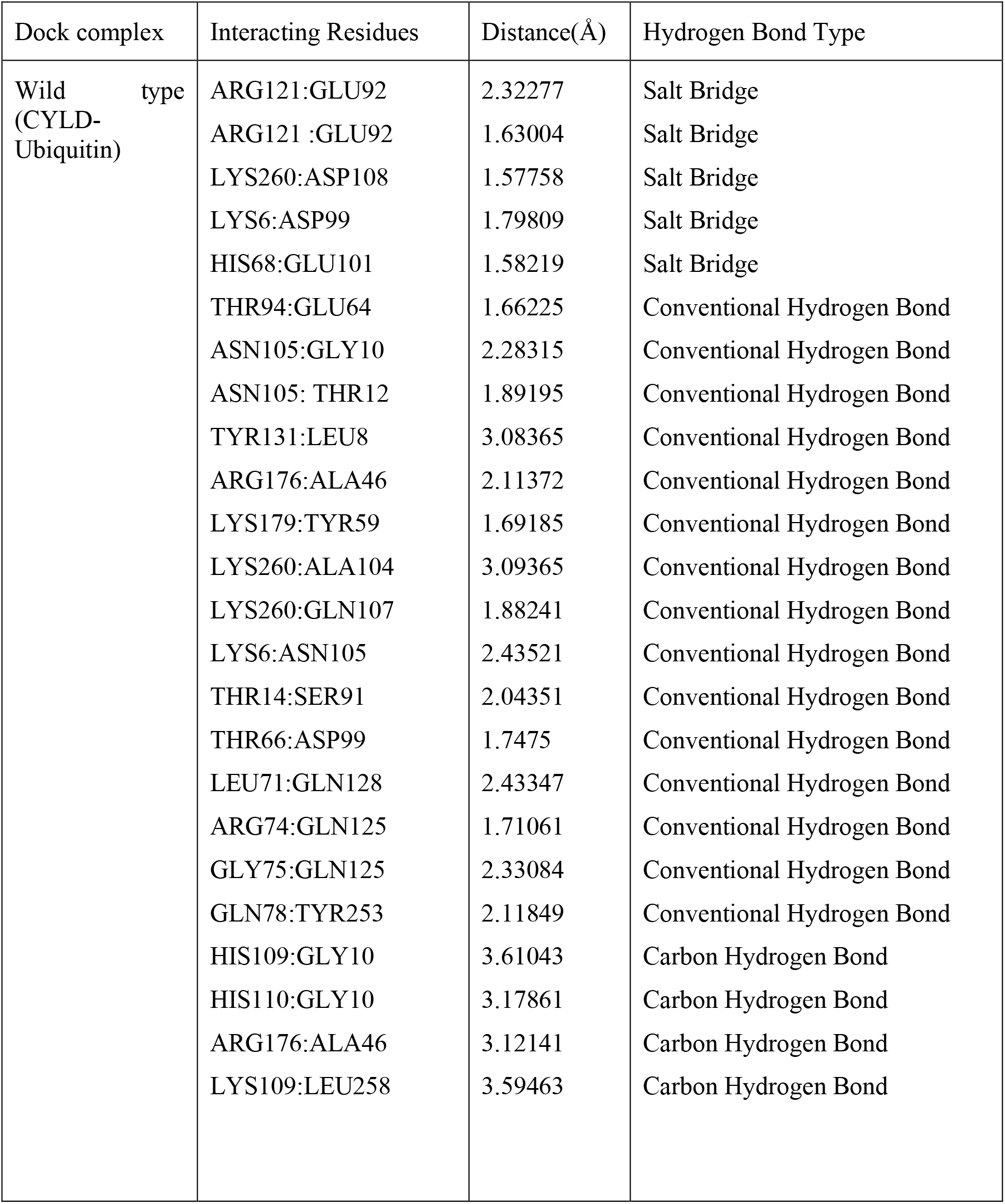
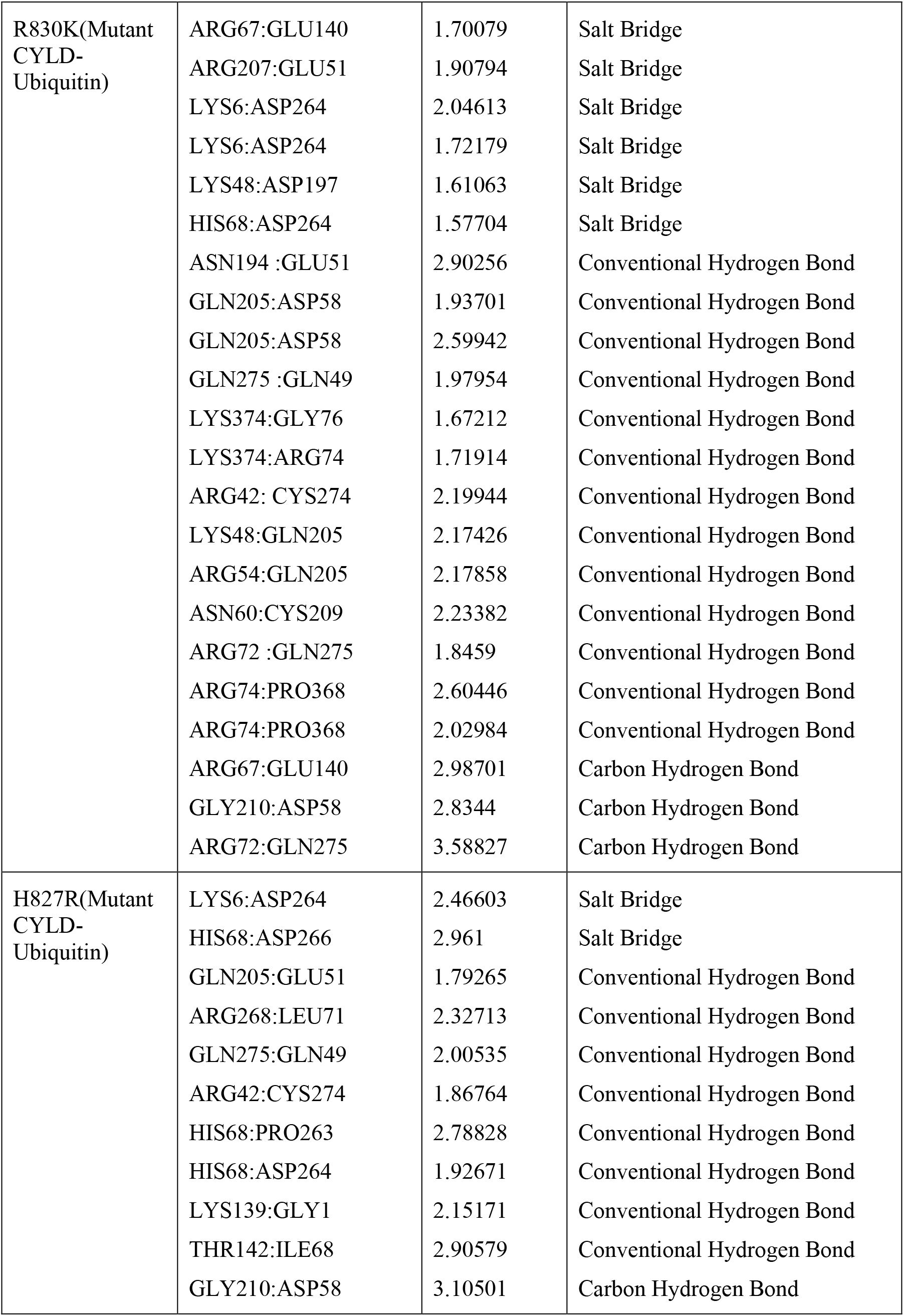
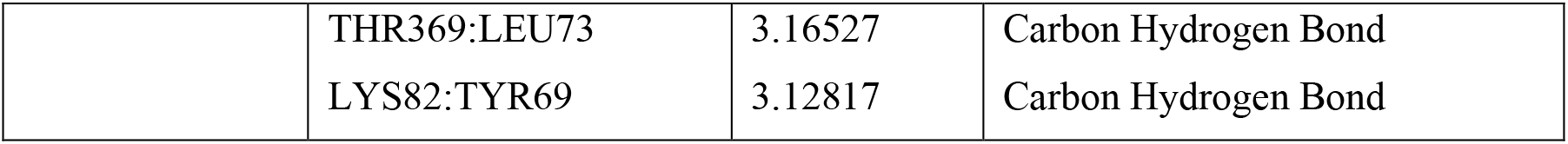
Hydrogen bond interactions between amino acid residues of CYLD (wild & Mutant variant)-Ubiquitin docking complexes given below:

| Dock complex | Interacting Residues | Distance(Å) | Hydrogen Bond Type |
| --- | --- | --- | --- |
| Wild type<br>(CYLD-Ubiquitin) | ARG121:GLU92 | 2.32277 | Salt Bridge |
|  | ARG121 :GLU92 | 1.63004 | Salt Bridge |
|  | LYS260:ASP108 | 1.57758 | Salt Bridge |
|  | LYS6:ASP99 | 1.79809 | Salt Bridge |
|  | HIS68:GLU101 | 1.58219 | Salt Bridge |
|  | THR94:GLU64 | 1.66225 | Conventional Hydrogen Bond |
|  | ASN105:GLY10 | 2.28315 | Conventional Hydrogen Bond |
|  | ASN105: THR12 | 1.89195 | Conventional Hydrogen Bond |
|  | TYR131:LEU8 | 3.08365 | Conventional Hydrogen Bond |
|  | ARG176:ALA46 | 2.11372 | Conventional Hydrogen Bond |
|  | LYS179:TYR59 | 1.69185 | Conventional Hydrogen Bond |
|  | LYS260:ALA104 | 3.09365 | Conventional Hydrogen Bond |
|  | LYS260:GLN107 | 1.88241 | Conventional Hydrogen Bond |
|  | LYS6:ASN105 | 2.43521 | Conventional Hydrogen Bond |
|  | THR14:SER91 | 2.04351 | Conventional Hydrogen Bond |
|  | THR66:ASP99 | 1.7475 | Conventional Hydrogen Bond |
|  | LEU71:GLN128 | 2.43347 | Conventional Hydrogen Bond |
|  | ARG74:GLN125 | 1.71061 | Conventional Hydrogen Bond |
|  | GLY75:GLN125 | 2.33084 | Conventional Hydrogen Bond |
|  | GLN78:TYR253 | 2.11849 | Conventional Hydrogen Bond |
|  | HIS109:GLY10 | 3.61043 | Carbon Hydrogen Bond |
|  | HIS110:GLY10 | 3.17861 | Carbon Hydrogen Bond |
|  | ARG176:ALA46 | 3.12141 | Carbon Hydrogen Bond |
|  | LYS109:LEU258 | 3.59463 | Carbon Hydrogen Bond |
| R830K(Mutant CYLD-Ubiquitin) | ARG67:GLU140 | 1.70079 | Salt Bridge |
|  | ARG207:GLU51 | 1.90794 | Salt Bridge |
|  | LYS6:ASP264 | 2.04613 | Salt Bridge |
|  | LYS6:ASP264 | 1.72179 | Salt Bridge |
|  | LYS48:ASP197 | 1.61063 | Salt Bridge |
|  | HIS68:ASP264 | 1.57704 | Salt Bridge |
|  | ASN194 :GLU51 | 2.90256 | Conventional Hydrogen Bond |
|  | GLN205:ASP58 | 1.93701 | Conventional Hydrogen Bond |
|  | GLN205:ASP58 | 2.59942 | Conventional Hydrogen Bond |
|  | GLN275 :GLN49 | 1.97954 | Conventional Hydrogen Bond |
|  | LYS374:GLY76 | 1.67212 | Conventional Hydrogen Bond |
|  | LYS374:ARG74 | 1.71914 | Conventional Hydrogen Bond |
|  | ARG42: CYS274 | 2.19944 | Conventional Hydrogen Bond |
|  | LYS48:GLN205 | 2.17426 | Conventional Hydrogen Bond |
|  | ARG54:GLN205 | 2.17858 | Conventional Hydrogen Bond |
|  | ASN60:CYS209 | 2.23382 | Conventional Hydrogen Bond |
|  | ARG72 :GLN275 | 1.8459 | Conventional Hydrogen Bond |
|  | ARG74:PRO368 | 2.60446 | Conventional Hydrogen Bond |
|  | ARG74:PRO368 | 2.02984 | Conventional Hydrogen Bond |
|  | ARG67:GLU140 | 2.98701 | Carbon Hydrogen Bond |
|  | GLY210:ASP58 | 2.8344 | Carbon Hydrogen Bond |
|  | ARG72:GLN275 | 3.58827 | Carbon Hydrogen Bond |
| H827R(Mutant CYLD-Ubiquitin) | LYS6:ASP264 | 2.46603 | Salt Bridge |
|  | HIS68:ASP266 | 2.961 | Salt Bridge |
|  | GLN205:GLU51 | 1.79265 | Conventional Hydrogen Bond |
|  | ARG268:LEU71 | 2.32713 | Conventional Hydrogen Bond |
|  | GLN275:GLN49 | 2.00535 | Conventional Hydrogen Bond |
|  | ARG42:CYS274 | 1.86764 | Conventional Hydrogen Bond |
|  | HIS68:PRO263 | 2.78828 | Conventional Hydrogen Bond |
|  | HIS68:ASP264 | 1.92671 | Conventional Hydrogen Bond |
|  | LYS139:GLY1 | 2.15171 | Conventional Hydrogen Bond |
|  | THR142:ILE68 | 2.90579 | Conventional Hydrogen Bond |
|  | GLY210:ASP58 | 3.10501 | Carbon Hydrogen Bond |
|  | THR369:LEU73 | 3.16527 | Carbon Hydrogen Bond |
|  | LYS82:TYR69 | 3.12817 | Carbon Hydrogen Bond |

**S9 Table:**
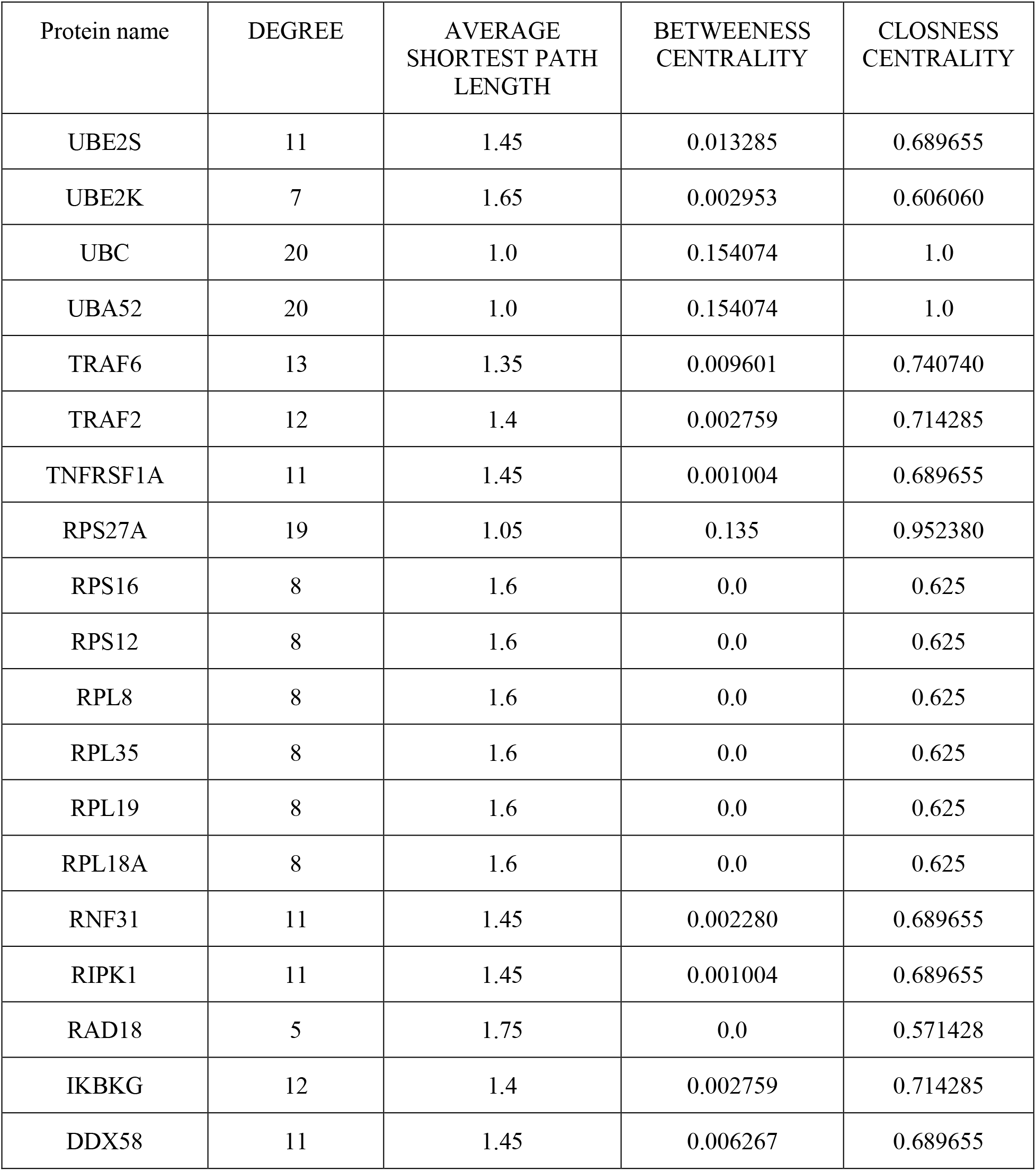
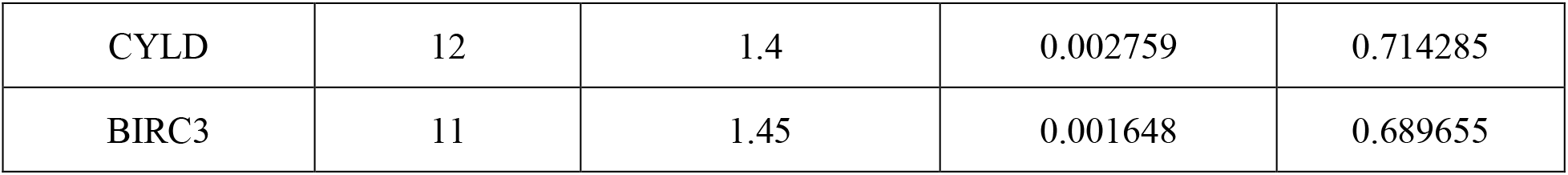
Protein-protein interaction analysis using Cytoscape.

**S10 Table:**
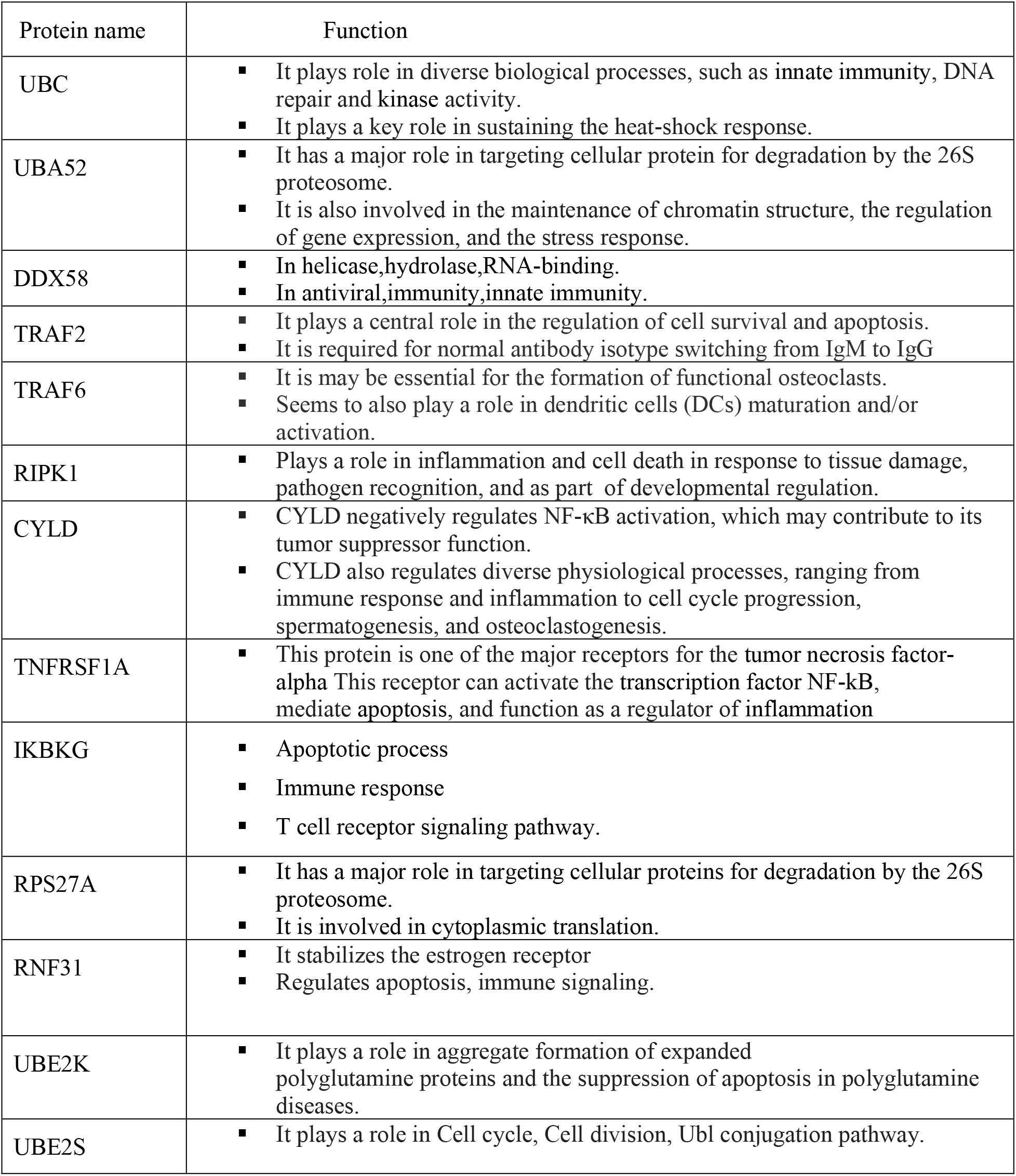
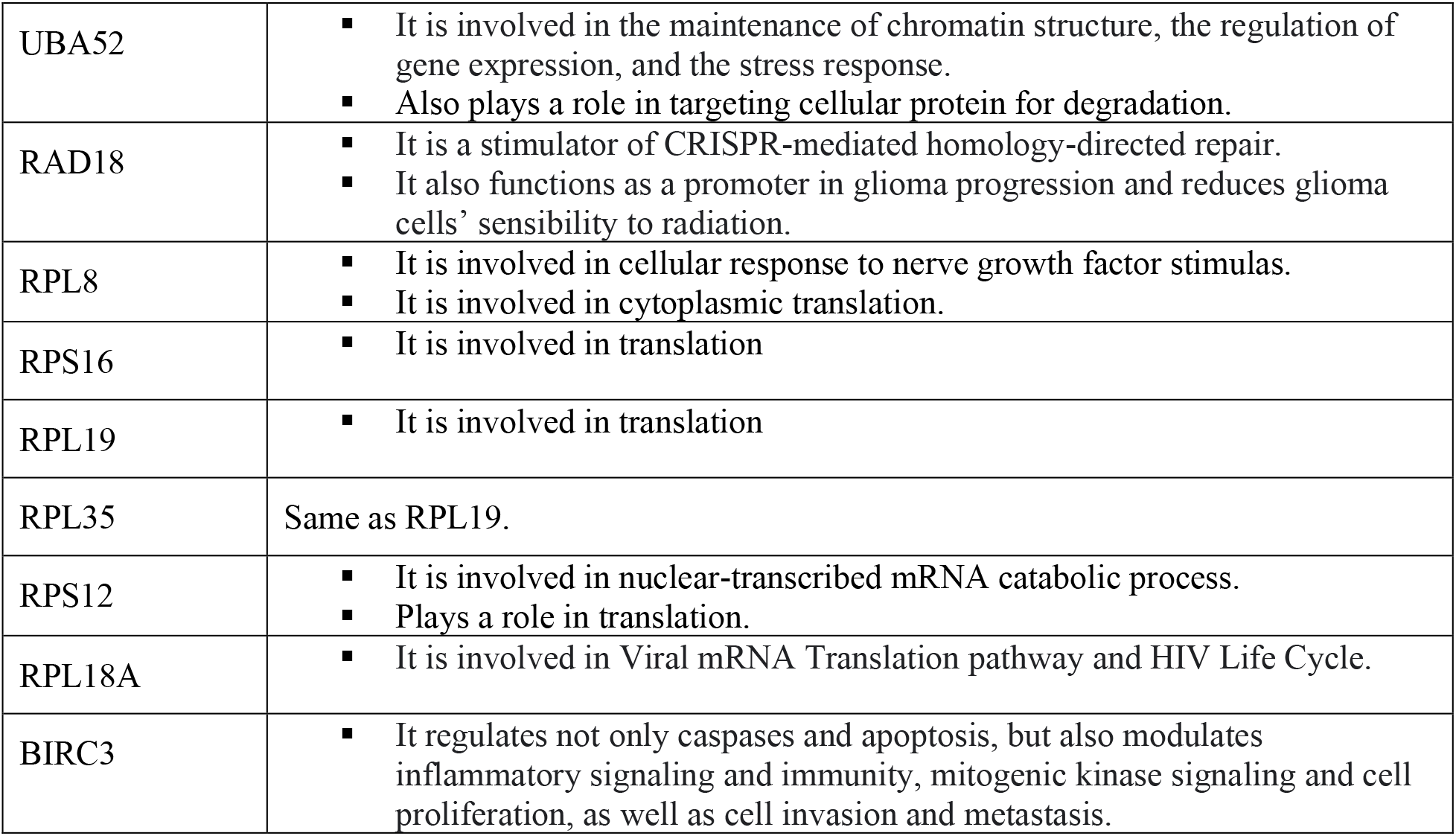
Functions of proteins interacting with CYLD in the PPI network.

