## Supplementary figures and tables for "A Computational Approach for Structural and Functional Analyses of Disease-associated Mutations in the Human *CYLD* Gene"

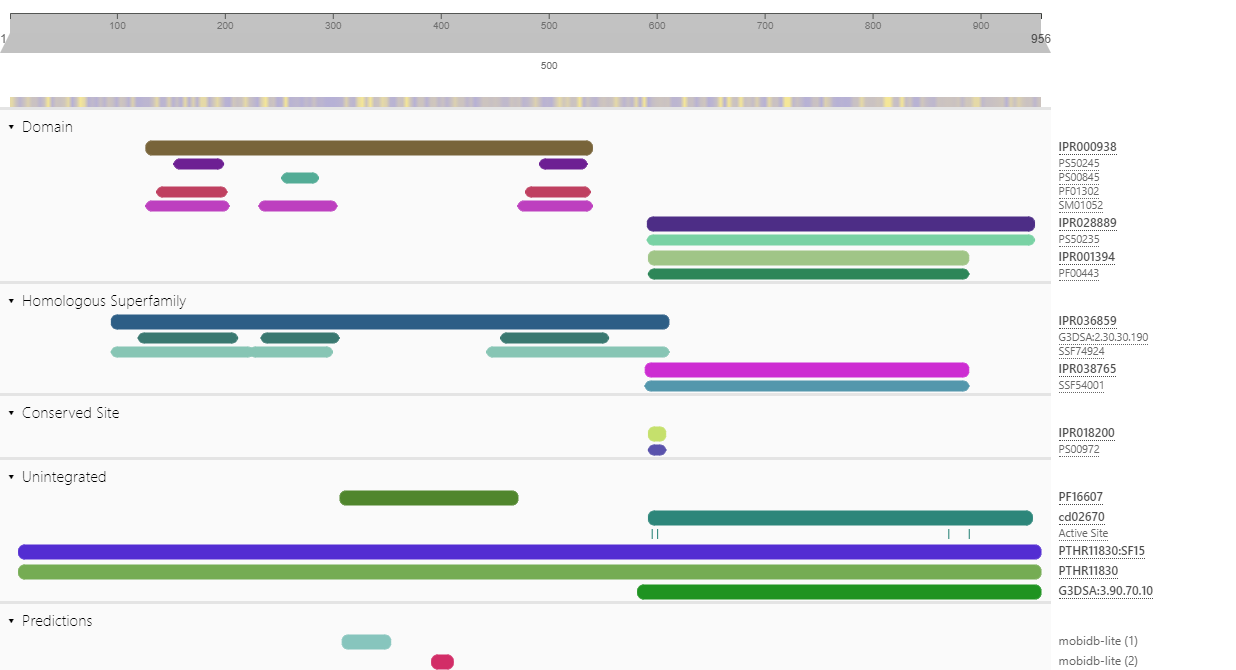

**S1_Fig: Domain identification of CYLD using InterPro server. (**CYLD contains two functional domains namely: Cap-Gly domain (127-540 Amino acid) and USP domain (592-950 AA)).

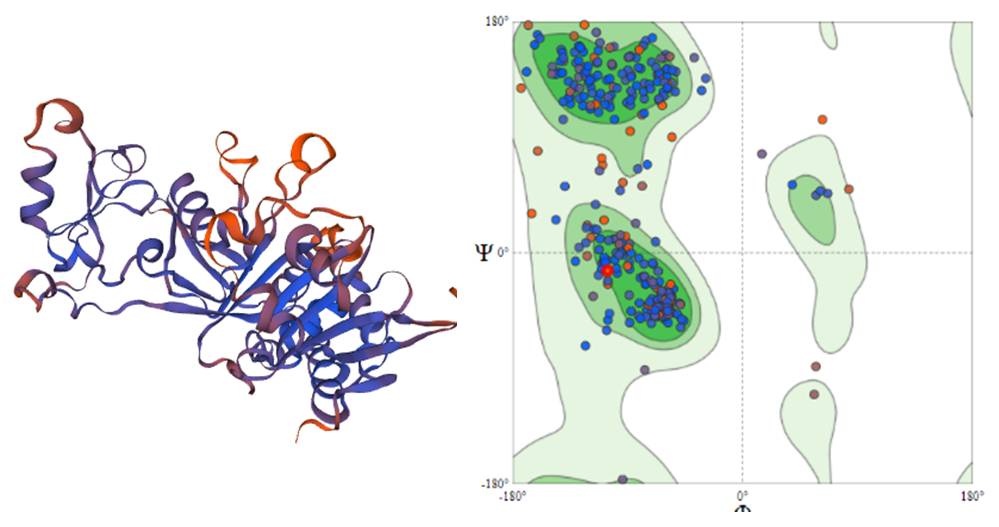

**S2_Fig: The 3D structure of CYLD protein (USP domain) constructed by SWISS MODEL and structural quality assessment using Ramachandran plot**

**S1 Table: Prediction of known disease-associated variants**

| SNP | Amino acid change | Panther | SIFT | Polyphen-2 | PHD SNP | Predict SNP | Disease Association |
| --- | --- | --- | --- | --- | --- | --- | --- |
| rs200759332 | T389R | PD | D | D | N | D | Familial multiple trichoepitheliomata (ClinVar)  Familial cylindromatosis (ClinVar)  Brooke-Spiegler syndrome (ClinVar) |
| rs200494719 | G431E | PD | D | D | N | D | Familial multiple trichoepitheliomata (ClinVar)  Familial cylindromatosis (ClinVar)  Brooke-Spiegler syndrome (ClinVar) |
| rs886040883 | D681H | PD | D | D | D | D | Familial cylindromatosis (ClinVar) |
| rs121908389 | E747G | PD | D | D | D | D | MFT1 and BRSS (UniProt)  Familial multiple trichoepitheliomata (ClinVar)  Brooke-Spiegler syndrome (ClinVar) |
| rs886040889 | L781P | PD | D | D | D | D | Brooke-Spiegler syndrome (ClinVar) |

*PD = Probably Damaging

*D = Deleterious

**S2 Table: Conservancy analysis using Consurf server**

| ID | substitution | Conservation Score | Prediction |
| --- | --- | --- | --- |
| [**rs971330819**](https://www.ncbi.nlm.nih.gov/snp/rs971330819) | V478A | 9 | Highly conserved and buried (s) |
| [**rs1363261645**](https://www.ncbi.nlm.nih.gov/snp/rs1363261645) | R489H | 9 | Highly conserved and buried (s) |
| [**rs758765051**](https://www.ncbi.nlm.nih.gov/snp/rs758765051) | L610F | 8 | Conserved and buried |
| [**rs747739683**](https://www.ncbi.nlm.nih.gov/snp/rs747739683) | I644T | 9 | Highly conserved and buried (s) |
| [**rs1404766168**](https://www.ncbi.nlm.nih.gov/snp/rs1404766168) | L648R | 9 | Highly conserved and buried (s) |
| [**rs964054055**](https://www.ncbi.nlm.nih.gov/snp/rs964054055) | P698T | 9 | Highly conserved and exposed (f) |
| [**rs964054055**](https://www.ncbi.nlm.nih.gov/snp/rs964054055) | P698S | 9 | Highly conserved and exposed (f) |
| [**rs1449388332**](https://www.ncbi.nlm.nih.gov/snp/rs1449388332) | P698L | 9 | Highly conserved and exposed (f) |
| [**rs121908389**](https://www.ncbi.nlm.nih.gov/snp/rs121908389) | E747G | 9 | Highly conserved and exposed (f) |
| [**rs886040889**](https://www.ncbi.nlm.nih.gov/snp/rs886040889) | L781P | 6 | Average conserved and buried |
| [**rs773384548**](https://www.ncbi.nlm.nih.gov/snp/rs773384548) | H827P | 9 | Highly conserved and exposed (f) |
| [**rs773384548**](https://www.ncbi.nlm.nih.gov/snp/rs773384548) | H827R | 9 | Highly conserved and exposed (f) |
| [**rs940896803**](https://www.ncbi.nlm.nih.gov/snp/rs940896803) | R830K | 9 | Highly conserved and exposed (f) |
| [**rs772100161**](https://www.ncbi.nlm.nih.gov/snp/rs772100161) | V864F | 9 | Highly conserved and buried (s) |
| [**rs763179319**](https://www.ncbi.nlm.nih.gov/snp/rs763179319) | I867K | 9 | Highly conserved and buried (s) |
| [**rs763179319**](https://www.ncbi.nlm.nih.gov/snp/rs763179319) | I867R | 9 | Highly conserved and buried (s) |
| [**rs754721077**](https://www.ncbi.nlm.nih.gov/snp/rs754721077) | H871Q | 9 | Highly conserved and exposed (f) |
| [**rs1597094057**](https://www.ncbi.nlm.nih.gov/snp/rs1597094057) | R894W | 9 | Highly conserved and exposed (f) |

**S3 Table: Tm score and RMSD value predicted by TM align tool**

| Substitution | **Tm Score** | **RMSD Value** |
| --- | --- | --- |
| V478A | 0.97849 | 0 |
| R489H | 0.97849 | 0 |
| L610F | 0.88451 | 2.03 |
| I644T | 0.89097 | 2.05 |
| L648R | 0.89007 | 2.14 |
| P698T | 0.90043 | 1.98 |
| P698S | 0.89614 | 2.12 |
| P698L | 0.86165 | 1.9 |
| E747G | 0.86029 | 2.11 |
| L781P | 0.89167 | 2.15 |
| H827P | 0.96345 | 1.75 |
| H827R | 0.89362 | 2.21 |
| R830K | 0.84714 | 1.95 |
| V864F | 0.88219 | 2.25 |
| I867K | 0.91058 | 1.95 |
| I867R | 0.85839 | 1.93 |
| H871Q | 0.90859 | 1.88 |
| R894W | 0.88827 | 2.13 |

**S4 Table: Swiss model Result**

| **Substitution** | **SOLVATION** | **TORSION** | **QMEAN** | **Cβ** | **ALL ATOM** | **Template** |
| --- | --- | --- | --- | --- | --- | --- |
| E747G | -0.4 | -2.48 | -2.56 | -0.1 | -2.01 | 2vhf.2.A |
| L781P | -0.54 | -2.17 | -2.29 | 0.04 | -2.18 | 2vhf.2.A |
| I644T | -0.61 | -2.26 | -2.43 | -0.15 | -2.29 | 2vhf.2.A |
| H871Q | -0.54 | -2.3 | -2.43 | -0.01 | -2.21 | 2vhf.2.A |
| I867K | -0.55 | -2.32 | -2.44 | 0.04 | -2.18 | 2vhf.2.A |
| I867R | -0.49 | -2.35 | -2.45 | 0.06 | -2.22 | 2vhf.2.A |
| V864F | -0.4 | -2.54 | -2.59 | -0.02 | -1.94 | 2vhf.2.A |
| H827P | -0.54 | -2.27 | -2.38 | 0.04 | -2.15 | 2vhf.2.A |
| R830K | -0.54 | -2.26 | -2.4 | -0.1 | -2.21 | 2vhf.2.A |
| P698T | -0.33 | -2.68 | -2.69 | 0.05 | -1.92 | 2vhf.2.A |
| L648R | -0.7 | -2.24 | -2.44 | -0.13 | -2.25 | 2vhf.2.A |
| L610F | -0.55 | -2.27 | -2.41 | -0.01 | -2.22 | 2vhf.2.A |
| H827R | -0.51 | -2.34 | -2.44 | 0.07 | -2.21 | 2vhf.2.A |
| Wild type | -0.53 | -2.3 | -2.42 | -0.01 | -2.24 | 2vhf.2.A |

**S5 Table: Structural effect of the point mutation predicted by HOPE project**

| RESIDUE | STRUCTURE | PROPERTIES |
| --- | --- | --- |
| E747G | 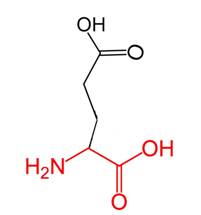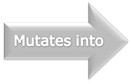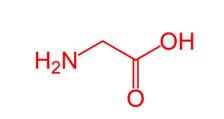 | There is a difference in charge between the wild-type and mutant amino acid.  The charge of the wild-type residue is lost by this mutation. This can cause loss of interactions with other molecules.  The wild-type and mutant amino acids differ in size.  The mutant residue is smaller than the wild-type residue.  This will cause a possible loss of external interactions.  The hydrophobicity of the wild-type and mutant residue differs. |
| L781P | 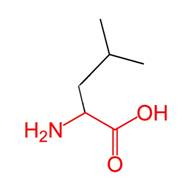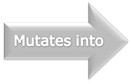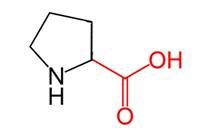 | The wild-type and mutant amino acids differ in size.  The mutant residue is smaller than the wild-type residue.  The mutation will cause an empty space in the core of the protein |
| I644T | 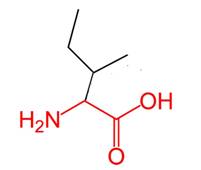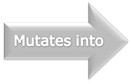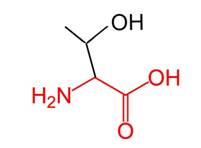 | The wild-type and mutant amino acids differ in size.  The mutant residue is smaller than the wild-type residue.  The mutation will cause an empty space in the core of the protein.  The hydrophobicity of the wild-type and mutant residue differs.  The mutation will cause loss of hydrophobic interactions in the core of the protein |
| H871Q | 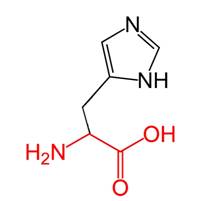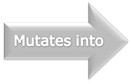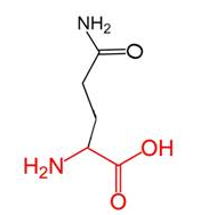 | The wild-type and mutant amino acids differ in size.  The mutant residue is smaller than the wild-type residue.  This will cause a possible loss of external interactions |
| I867K | 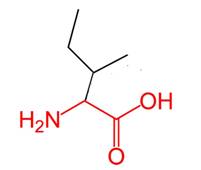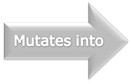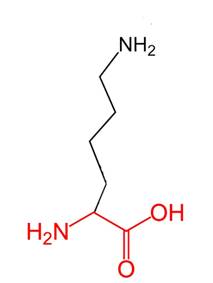 | There is a difference in charge between the wild-type and mutant amino acid.  The mutation introduces a charge at this position, this can cause repulsion between the mutant residue and neighboring residues.  The wild-type and mutant amino acids differ in size.  The mutant residue is bigger than the wild-type residue.  The residue is located on the surface of the protein, mutation of this residue can disturb interactions with other molecules or other parts of the protein.  The hydrophobicity of the wild-type and mutant residue differs.  The mutation might cause loss of hydrophobic interactions with other molecules on the surface of the protein. |
| I867R | 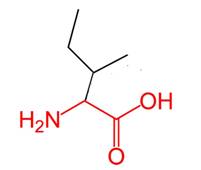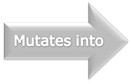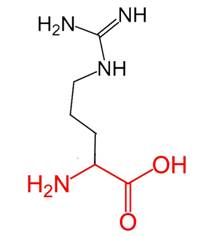 | There is a difference in charge between the wild-type and mutant amino acid.  The mutation introduces a charge at this position, this can cause repulsion between the mutant residue and neighboring residues.  The wild-type and mutant amino acids differ in size.  The mutant residue is bigger than the wild-type residue.  The residue is located on the surface of the protein, mutation of this residue can disturb interactions with other molecules or other parts of the protein.  The hydrophobicity of the wild-type and mutant residue differs.  The mutation might cause loss of hydrophobic interactions with other molecules on the surface of the protein |
| V864F | 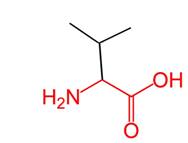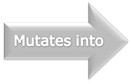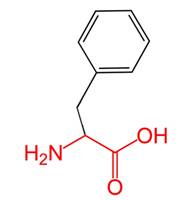 | The wild-type and mutant amino acids differ in size.  The mutant residue is bigger than the wild-type residue.  The wild-type residue was buried in the core of the protein. The mutant residue is bigger and probably will not fit. |
| H827P | 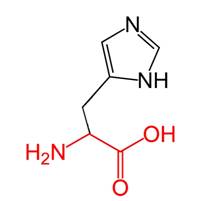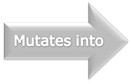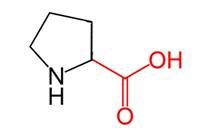 | The wild-type and mutant amino acids differ in size.  The mutant residue is smaller than the wild-type residue.  This will cause a possible loss of external interactions.  The hydrophobicity of the wild-type and mutant residue differs. |
| R830K | 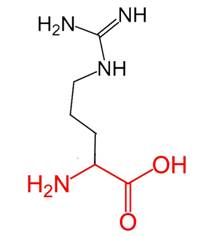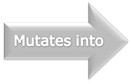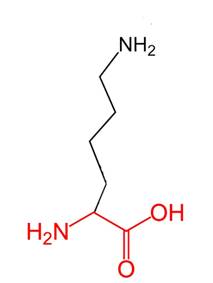 | The wild-type and mutant amino acids differ in size.  The mutant residue is smaller than the wild-type residue.  The mutation will cause an empty space in the core of the protein. |
| P698T | 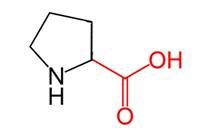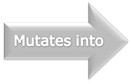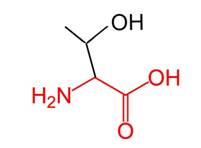 | The hydrophobicity of the wild-type and mutant residue differs.  The mutation might cause loss of hydrophobic interactions with other molecules on the surface of the protein. |
| L648R | 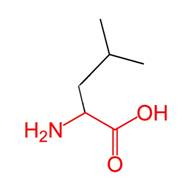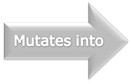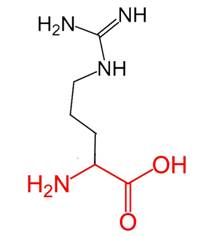 | There is a difference in charge between the wild-type and mutant amino acid.  The mutant residue introduces a charge in a buried residue which can lead to protein folding problems.  The wild-type and mutant amino acids differ in size.  The mutant residue is bigger than the wild-type residue.  The wild-type residue was buried in the core of the protein. The mutant residue is bigger and probably will not fit.  The hydrophobicity of the wild-type and mutant residue differs.  The mutation will cause loss of hydrophobic interactions in the core of the protein. |
| P698L | 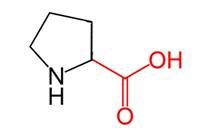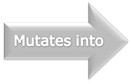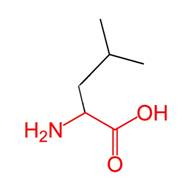 | The wild-type and mutant amino acids differ in size.  The mutant residue is bigger than the wild-type residue.  The residue is located on the surface of the protein, mutation of this residue can disturb interactions with other molecules or other parts of the protein |
| L610F | 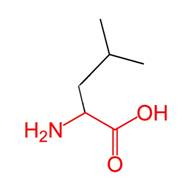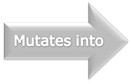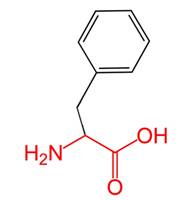 | The wild-type and mutant amino acids differ in size.  The mutant residue is bigger than the wild-type residue.  The wild-type residue was buried in the core of the protein. The mutant residue is bigger and probably will not fit. |
| H827R | 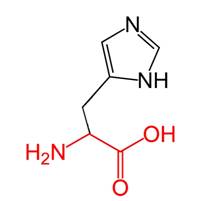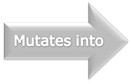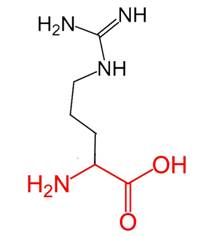 | There is a difference in charge between the wild-type and mutant amino acid.  The mutation introduces a charge at this position, this can cause repulsion between the mutant residue and neighboring residues.  The wild-type and mutant amino acids differ in size.  The mutant residue is bigger than the wild-type residue.  The residue is located on the surface of the protein, mutation of this residue can disturb interactions with other molecules or other parts of the protein |

**S6 Table: Energy minimization state of wild and 13 mutant variants determined by Swiss PDB Viewer**

| **Substitution** | **Presence of Clash / Hydrogen bond** | **Number of Rotamer** | **Total Energy after Energy Minimization(I-TASSER)** |
| --- | --- | --- | --- |
| E747G | Hydrogen bond | 1 | -20025.781 |
| L781P | Both | 2 | -18904.959 |
| I644T | Both | 3 | -20062.729 |
| H871Q | Both | 15 | -18391.238 |
| I867K | Both | 19 | -19640.801 |
| I867R | Both | 28 | -20214.043 |
| V864F | Both | 5 | -19982.813 |
| H827P | Both | 2 | -19861.77 |
| R830K | Both | 19 | -19823.832 |
| P698T | Both | 3 | -20183.6 |
| L648R | Both | 26 | -20398.639 |
| L610F | Both | 5 | -20294.15 |
| H827R | Both | 28 | -15956.584 |
| **Wild type** |  |  | **-20130.191** |

**S7 Table: Binding affinity and dissociation constant of all docking complex shown in the table**

| Residue | Prodigy Binding affinity(ΔG (kcal mol-1) | Dissociation constant Kd (M) at 25.0 ℃ |
| --- | --- | --- |
| Wild type | -14.6 | 1.8E-11 |
| L610F | -15.2 | 7.00E-12 |
| I644T | -15.9 | 2.30E-12 |
| L648R | -14.8 | 1.4E-11 |
| P698T | -13 | 2.80E-10 |
| E747G | -15.7 | 3.3E-12 |
| L781P | -13.4 | 1.4E-10 |
| H827P | -14 | 5.5E-11 |
| H827R | -12.8 | 4.30E-10 |
| R830K | -12.7 | 5.20E-10 |
| V864F | -15.1 | 8.80E-12 |
| I867K | -14.3 | 3.20E-11 |
| I867R | -15.3 | 6.4E-12 |
| H871Q | -16 | 1.7E-12 |

**S8 Table: Hydrogen bond interactions between amino acid residues of CYLD (wild & Mutant variant)-Ubiquitin docking complexes given below:**

| Dock complex | Interacting Residues | Distance(Å) | Hydrogen Bond Type |
| --- | --- | --- | --- |
| Wild type (CYLD-Ubiquitin) | ARG121:GLU92  ARG121 :GLU92  LYS260:ASP108  LYS6:ASP99  HIS68:GLU101  THR94:GLU64  ASN105:GLY10  ASN105: THR12  TYR131:LEU8  ARG176:ALA46  LYS179:TYR59  LYS260:ALA104  LYS260:GLN107  LYS6:ASN105  THR14:SER91  THR66:ASP99  LEU71:GLN128  ARG74:GLN125  GLY75:GLN125  GLN78:TYR253  HIS109:GLY10  HIS110:GLY10  ARG176:ALA46  LYS109:LEU258 | 2.32277  1.63004  1.57758  1.79809  1.58219  1.66225  2.28315  1.89195  3.08365  2.11372  1.69185  3.09365  1.88241  2.43521  2.04351  1.7475  2.43347  1.71061  2.33084  2.11849  3.61043  3.17861  3.12141  3.59463 | Salt Bridge  Salt Bridge  Salt Bridge  Salt Bridge  Salt Bridge  Conventional Hydrogen Bond  Conventional Hydrogen Bond  Conventional Hydrogen Bond  Conventional Hydrogen Bond  Conventional Hydrogen Bond  Conventional Hydrogen Bond  Conventional Hydrogen Bond  Conventional Hydrogen Bond  Conventional Hydrogen Bond  Conventional Hydrogen Bond  Conventional Hydrogen Bond  Conventional Hydrogen Bond  Conventional Hydrogen Bond  Conventional Hydrogen Bond  Conventional Hydrogen Bond  Carbon Hydrogen Bond  Carbon Hydrogen Bond  Carbon Hydrogen Bond  Carbon Hydrogen Bond |
| R830K(Mutant CYLD-Ubiquitin) | ARG67:GLU140  ARG207:GLU51  LYS6:ASP264  LYS6:ASP264  LYS48:ASP197  HIS68:ASP264  ASN194 :GLU51  GLN205:ASP58  GLN205:ASP58  GLN275 :GLN49  LYS374:GLY76  LYS374:ARG74  ARG42: CYS274  LYS48:GLN205  ARG54:GLN205  ASN60:CYS209  ARG72 :GLN275  ARG74:PRO368  ARG74:PRO368  ARG67:GLU140  GLY210:ASP58  ARG72:GLN275 | 1.70079  1.90794  2.04613  1.72179  1.61063  1.57704  2.90256  1.93701  2.59942  1.97954  1.67212  1.71914  2.19944  2.17426  2.17858  2.23382  1.8459  2.60446  2.02984  2.98701  2.8344  3.58827 | Salt Bridge  Salt Bridge  Salt Bridge  Salt Bridge  Salt Bridge  Salt Bridge  Conventional Hydrogen Bond  Conventional Hydrogen Bond  Conventional Hydrogen Bond  Conventional Hydrogen Bond  Conventional Hydrogen Bond  Conventional Hydrogen Bond  Conventional Hydrogen Bond  Conventional Hydrogen Bond  Conventional Hydrogen Bond  Conventional Hydrogen Bond  Conventional Hydrogen Bond  Conventional Hydrogen Bond  Conventional Hydrogen Bond  Carbon Hydrogen Bond  Carbon Hydrogen Bond  Carbon Hydrogen Bond |
| H827R(Mutant CYLD-Ubiquitin) | LYS6:ASP264  HIS68:ASP266  GLN205:GLU51  ARG268:LEU71  GLN275:GLN49  ARG42:CYS274  HIS68:PRO263  HIS68:ASP264  LYS139:GLY1  THR142:ILE68  GLY210:ASP58  THR369:LEU73  LYS82:TYR69 | 2.46603  2.961  1.79265  2.32713  2.00535  1.86764  2.78828  1.92671  2.15171  2.90579  3.10501  3.16527  3.12817 | Salt Bridge  Salt Bridge  Conventional Hydrogen Bond  Conventional Hydrogen Bond  Conventional Hydrogen Bond  Conventional Hydrogen Bond  Conventional Hydrogen Bond  Conventional Hydrogen Bond  Conventional Hydrogen Bond  Conventional Hydrogen Bond  Carbon Hydrogen Bond  Carbon Hydrogen Bond  Carbon Hydrogen Bond |

**S9 Table: Protein-protein interaction analysis using Cytoscape**

| Protein name | DEGREE | AVERAGE SHORTEST PATH LENGTH | BETWEENESS CENTRALITY | CLOSNESS CENTRALITY |
| --- | --- | --- | --- | --- |
| UBE2S | 11 | 1.45 | 0.013285 | 0.689655 |
| UBE2K | 7 | 1.65 | 0.002953 | 0.606060 |
| UBC | 20 | 1.0 | 0.154074 | 1.0 |
| UBA52 | 20 | 1.0 | 0.154074 | 1.0 |
| TRAF6 | 13 | 1.35 | 0.009601 | 0.740740 |
| TRAF2 | 12 | 1.4 | 0.002759 | 0.714285 |
| TNFRSF1A | 11 | 1.45 | 0.001004 | 0.689655 |
| RPS27A | 19 | 1.05 | 0.135 | 0.952380 |
| RPS16 | 8 | 1.6 | 0.0 | 0.625 |
| RPS12 | 8 | 1.6 | 0.0 | 0.625 |
| RPL8 | 8 | 1.6 | 0.0 | 0.625 |
| RPL35 | 8 | 1.6 | 0.0 | 0.625 |
| RPL19 | 8 | 1.6 | 0.0 | 0.625 |
| RPL18A | 8 | 1.6 | 0.0 | 0.625 |
| RNF31 | 11 | 1.45 | 0.002280 | 0.689655 |
| RIPK1 | 11 | 1.45 | 0.001004 | 0.689655 |
| RAD18 | 5 | 1.75 | 0.0 | 0.571428 |
| IKBKG | 12 | 1.4 | 0.002759 | 0.714285 |
| DDX58 | 11 | 1.45 | 0.006267 | 0.689655 |
| CYLD | 12 | 1.4 | 0.002759 | 0.714285 |
| BIRC3 | 11 | 1.45 | 0.001648 | 0.689655 |

**S10 Table: Functions of proteins interacting with CYLD in the PPI network**

| Protein name | Function |
| --- | --- |
| UBC | - It plays role in diverse biological processes, such as innate immunity, DNA repair and kinase activity. - It plays a key role in sustaining the heat-shock response. |
| UBA52 | - It has a major role in targeting cellular protein for degradation by the 26S proteosome. - It is also involved in the maintenance of chromatin structure, the regulation of gene expression, and the stress response. |
| DDX58 | - In helicase,hydrolase,RNA-binding. - In antiviral,immunity,innate immunity. |
| TRAF2 | - It plays a central role in the regulation of cell survival and apoptosis. - It is required for normal antibody isotype switching from IgM to IgG |
| TRAF6 | - It is may be essential for the formation of functional osteoclasts. - Seems to also play a role in dendritic cells (DCs) maturation and/or activation. |
| RIPK1 | - Plays a role in inflammation and cell death in response to tissue damage, pathogen recognition, and as part of developmental regulation. |
| CYLD | - CYLD negatively regulates NF-κB activation, which may contribute to its tumor suppressor function. - CYLD also regulates diverse physiological processes, ranging from immune response and inflammation to cell cycle progression, spermatogenesis, and osteoclastogenesis. |
| TNFRSF1A | - This protein is one of the major receptors for the tumor necrosis factor-alpha This receptor can activate the transcription factor NF-kB, mediate apoptosis, and function as a regulator of inflammation |
| IKBKG | - Apoptotic process - Immune response - T cell receptor signaling pathway. |
| RPS27A | - It has a major role in targeting cellular proteins for degradation by the 26S proteosome. - It is involved in cytoplasmic translation. |
| RNF31 | - It stabilizes the estrogen receptor - Regulates apoptosis, immune signaling. |
| UBE2K | - It plays a role in aggregate formation of expanded polyglutamine proteins and the suppression of apoptosis in polyglutamine diseases. |
| UBE2S | - It plays a role in Cell cycle, Cell division, Ubl conjugation pathway. |
| UBA52 | - It is involved in the maintenance of chromatin structure, the regulation of gene expression, and the stress response. - Also plays a role in targeting cellular protein for degradation. |
| RAD18 | - It is a stimulator of CRISPR-mediated homology-directed repair. - It also functions as a promoter in glioma progression and reduces glioma cells’ sensibility to radiation. |
| RPL8 | - It is involved in cellular response to nerve growth factor stimulas. - It is involved in cytoplasmic translation. |
| RPS16 | - It is involved in translation |
| RPL19 | - It is involved in translation |
| RPL35 | Same as RPL19. |
| RPS12 | - It is involved in nuclear-transcribed mRNA catabolic process. - Plays a role in translation. |
| RPL18A | - It is involved in Viral mRNA Translation pathway and HIV Life Cycle. |
| BIRC3 | - It regulates not only caspases and apoptosis, but also modulates inflammatory signaling and immunity, mitogenic kinase signaling and cell proliferation, as well as cell invasion and metastasis. |
